# Terminal differentiation and persistence of effector regulatory T cells essential for the prevention of intestinal inflammation

**DOI:** 10.1101/2022.05.16.492030

**Authors:** Stanislav Dikiy, Andrew G. Levine, Paolo Giovanelli, Zhong-Min Wang, Giorgi Beroshvili, Yuri Pritykin, Chirag Krishna, Ariella Glasner, Christina S. Leslie, Alexander Y. Rudensky

**Author notes:** Senior author (Christina S. Leslie, Alexander Y. Rudensky). Department of Immunology and Microbiology, Scripps Research, 10550 North Torrey Pines Road, La Jolla, CA 92037, USA.

## Abstract

Regulatory T (Treg) cells represent a specialized CD4^+^ T cell lineage with essential anti-inflammatory functions. Recent studies of the adaptations of Treg cells to non-lymphoid tissues which enable their specialized immunosuppressive and tissue supportive functions raise questions about the underlying mechanisms of these adaptations and whether they represent stable differentiation or reversible activation states. Using novel genetic tools, we characterized the transcriptional programs of distinct colonic effector Treg cell types. We found that attenuated T cell receptor (TCR) signaling and acquisition of substantial TCR independent functionality appears to facilitate the terminal differentiation of a population of colonic effector Treg cells distinguished by stable expression of immunomodulatory cytokine interleukin-10 (IL-10). Functional studies revealed that this subset of effector Treg cells, but not their expression of IL-10, was indispensable for colonic health. These findings suggest core features of terminal differentiation of effector Treg cells in non-lymphoid tissues and their function therein.

## Introduction

Regulatory T (Treg) cells, a specialized suppressive subset of T cells, are defined by sustained expression of the lineage-specifying transcription factor (TF) Foxp3 ^1,2^. Treg cells are first and foremost essential for the prevention and control of autoimmunity as congenital deficiency due to inactivating mutations in *Foxp3,* or experimental depletion of these cells results in fulminant and ultimately fatal systemic autoimmune and inflammatory disease ^3–9^. However, more recent work has uncovered additional essential functions of Treg cells, including but not limited to modulating organismal metabolism, supporting tissue growth and regeneration, and enforcing tolerance to innocuous environmental and microbiota derived antigens ^10–18^. However, in cancerous settings, both the immunomodulatory and tissue supportive functions of Treg cells promote tumor growth and subvert anti-tumor immunity ^19–22^.

The processes governing Treg cell generation from thymocytes and naïve CD4^+^ T cells in the thymus and periphery, respectively, have been well characterized ^1^. However, the activation and subsequent differentiation of mature Treg cells upon entering secondary lymphoid organs (SLOs) and particularly non-lymphoid tissues, which result in a manifest diversification of Treg cell states and function, have only recently begun to be explored. Distinct Treg cell populations in the periphery have been described as having ‘resting’ versus ‘activated’ or ‘effector’ phenotypes. The former share features with naïve or central memory CD4^+^ T cells, while the latter have more potent suppressor activity, preferentially traffic to non-lymphoid tissue, and have been shown to expand under inflammatory conditions in response to TCR and cytokine stimulation ^23,24^. Exploring this distinction, several recent studies have leveraged genomic techniques to assess changes in gene expression and chromatin accessibility to identify transcription factors associated with the transition of Treg cells within the SLOs to those carrying out various functions in non-lymphoid tissues. One pair of studies suggests that stepwise expression of the TFs Nfil3, Batf, and Gata3 facilitates the acquisition of ‘effector’ molecules and migration to non-lymphoid tissues in both mouse and human Treg cells, while another study points to increased expression of Rorα and Gata3 as drivers of a ‘non-lymphoid’ Treg cell phenotype ^25–27^.

However, it remains unclear whether mature Treg cells are truly undergoing differentiation in the periphery into multiple effector states, akin to conventional CD4^+^ T cells. Alternatively, these cells may merely adopt distinct transient gene expression programs specified by ongoing environmental conditioning. In the latter scenario, Treg cells, retaining plasticity, would return to a ‘resting’ state upon the withdrawal of these conditioning stimuli and potentially adopt a different state of activation in response to distinct stimuli. Previous studies have suggested that this indeed occurs for Treg cells under a variety of perturbations including systemic autoimmunity, viral infection, as well as within tumors ^21,24^. At the same time, recent work has identified a population of Treg cells which durably express the TF T-bet and specifically control type I immune responses ^28^. Importantly, these cells were inefficient at controlling two other major types of immune responses, suggesting a highly specialized function with diminished plasticity and therefore potential terminal differentiation of this population.

A caveat of many comparative studies of ‘effector’ and ‘resting’ Treg cells is that they contrasted Treg cells residing in SLOs with those in non-lymphoid organs, or those within SLOs bearing markers suggesting preferential retention versus exit from that tissue ^23,25,27^. Since T cells in SLOs are overall more quiescent than those from non-lymphoid tissues, quiescence can be readily conflated with a less differentiated state in at least some of these studies ^29^. For this reason, we wished to study Treg cell differentiation specifically within a non-lymphoid tissue by contrasting the populations found therein. Additionally, we wanted to directly identify a Treg cell effector state using a validated functional molecule uncoupled from preferential homing to lymphoid versus non-lymphoid tissues. Using a fate-mapping approach tied to expression of the immunomodulatory cytokine interleukin (IL)-10, we found distinct, specialized populations of Treg cells in the colonic lamina propria, which were apparently terminally differentiated, and acquired a specialized gene expression program at least partly on the basis of attenuating signaling through the T cell receptor (TCR). Specific depletion of this specialized IL-10 expressing colonic Treg cell subset revealed its essential function in preventing colonic inflammation, while induced ablation of IL-10 itself suggested its redundancy for Treg cell function in this context.

## Results

### Identification of a stable effector Treg cell population

In order to track the fate of effector Treg cells, we generated a genetic mouse model to allow the identification, labeling, and tracking of effector Treg cells. As their distinguishing feature we chose the major immunosuppressive cytokine IL-10 as Treg cells have been shown to specifically deploy this cytokine to modulate various immune responses in diverse settings ^30–32^. We generated an IL-10 reporter allele by inserting the coding sequence for a tdTomato fluorescent reporter, fused by a T2A self-cleaving peptide with a tamoxifen-inducible Cre recombinase into the 3’ UTR of the *Il10* gene, with independent translation driven by an IRES sequence (Extended Data Fig. 1a). In *Il10^tdTomato-CreER^* mice, cells expressing IL-10 were readily detectable due to tdTomato expression and harbored a tamoxifen-inducible Cre recombinase. To enable tracking of Treg cells, we crossed these mice to *Foxp3^Thy1.1^* reporter mice ^33^. Analysis of tdTomato fluorescence demonstrated increased IL-10 expression by Treg cells versus conventional activated or naïve CD4^+^ T cells, as well as by activated (CD44^hi^CD62L^−^) versus resting (CD44^lo^CD62L^+^) Treg cells across a variety of lymphoid and non-lymphoid tissues (Fig. 1a). Importantly, high frequencies of tdTomato (IL-10)^+^ colonic macrophages and plasma cells, populations known to be important *in vivo* sources of IL-10, as well as differential intensity of tdTomato fluorescence among distinct IL-10 expressing cell types, suggested that the *Il10^tdTomato-CreER^* allele allows for a sensitive and faithful reporting of endogenous IL-10 expression (Extended Data Fig. 1b-d).

**Fig. 1.**
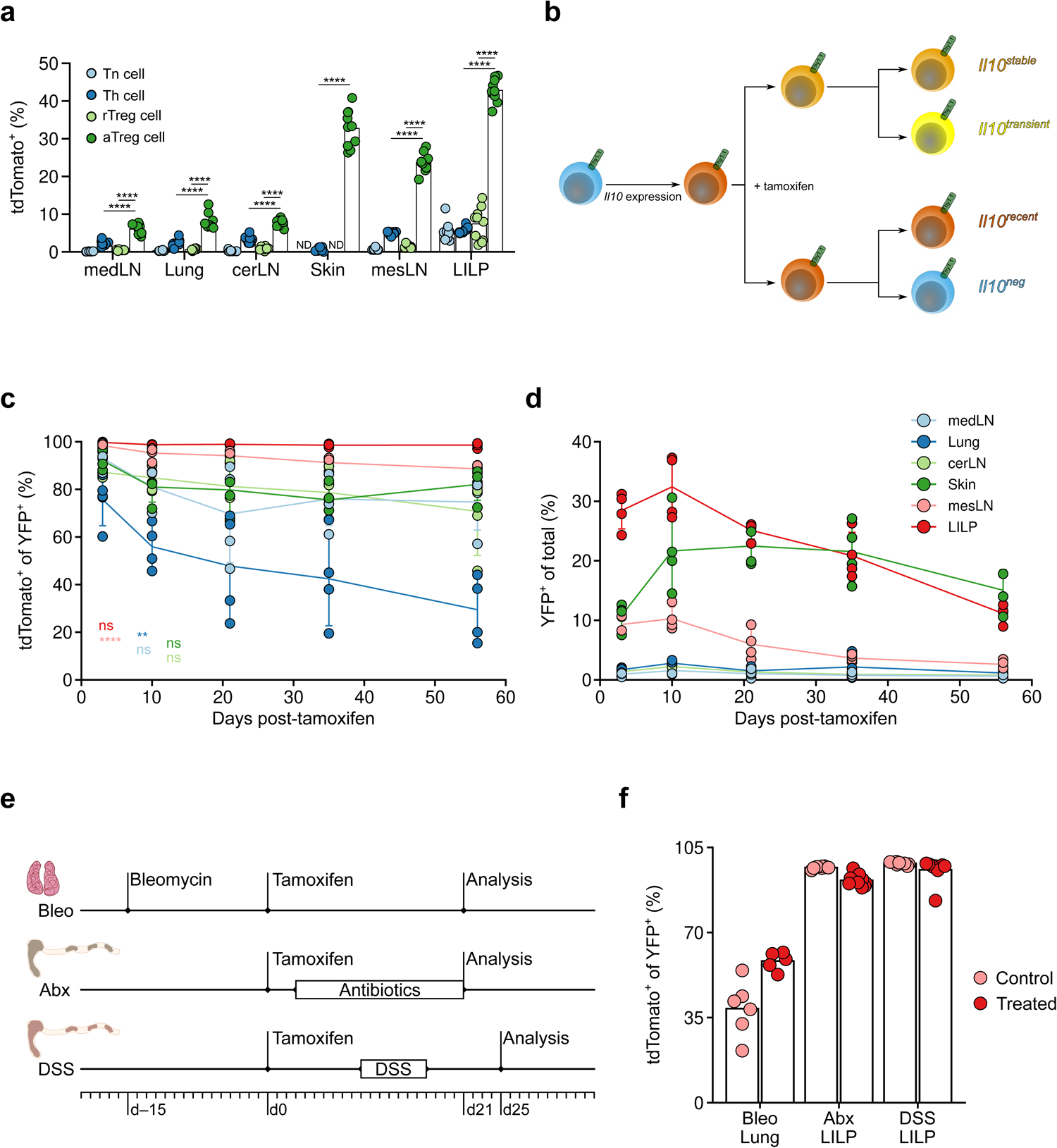
A population of colonic Treg cells stably expresses IL-10. a. Frequencies of tdTomato^+^ cells among naïve (Tn) (CD4^+^TCRβ^+^CD44^lo^Thy1.1^−^) and activated helper (Th) CD4^+^ T cells (CD4^+^TCRβ^+^CD44^hi^Thy1.1^−^), resting (rTreg) (Thy1.1^+^CD4^+^TCRβ^+^CD62L^+^), and activated (aTreg) (Thy1.1^+^CD4^+^TCRβ^+^CD62L^−^) Treg cells isolated from indicated tissues of 15-week-old *Il10^FM^* mice. ND: Tn and rTreg cells were not detected. Each point represents an individual mouse (n = 10) and data are representative of two independent experiments. Unpaired two-sided t-tests with Holm’s correction for multiple comparisons. b-d. *Il10^FM^* mice were treated at 7 weeks of age with tamoxifen and analyzed 3 to 56 days later. Mice analyzed together were littermates and treated at different times with tamoxifen in order be analyzed on the same day. Schematic of IL-10 fate-mapping experiments and all possible combinations of tdTomato and YFP expression by Treg cells therein (b). Frequencies of tdTomato^+^ cells among YFP^+^ (c) and YFP^+^ among all (d) Treg cells (Thy1.1^+^CD4^+^TCRβ^+^) isolated from indicated tissues of mice at indicated days after tamoxifen treatment. Each point represents and individual mouse, lines indicate mean per tissue across time points, bars indicate standard deviation, and data are pooled from two independent experiments (n = 4 mice per time point, with different mice for each time point). ANOVA with Holm’s correction for multiple comparisons. e, f. Analysis of stability of IL-10 expressing Treg cells with perturbations. For bleomycin challenge, 7-week-old *Il10^FM^* mice were treated intranasally with bleomycin or vehicle 15 days before treatment with tamoxifen and then analyzed 21 days later (Bleo). For microbiota depletion, 7-to 10-week-old *Il10^FM^* mice were treated with tamoxifen, then 3 days later placed on antibiotics-laden or control drinking water, and then analyzed 18 days later (Abx). For chemically induced colitis, 7-to 10-week-old *Il10^FM^* female mice were treated with tamoxifen, then 10 days later placed on drinking water containing 3% DSS or vehicle for 7 days, and analyzed 25 days post-tamoxifen (DSS). Experimental schematics (e) and frequencies of tdTomato^+^ cells among YFP^+^ Treg cells (Thy1.1^+^CD4^+^TCRβ^+^) isolated from the lungs (Bleo) or large intestine lamina propria (Abx, DSS) of challenged and littermate control *Il10^FM^* mice (f). Each point represents an individual mouse (n = 6 per group for Bleo, 8-9 per group for Abx and DSS) and data are pooled from two independent experiments for each perturbation. *P*-value > 0.05 ns, < 0.01 **, < 0.0001 ****

To assess the stability of IL-10 expression using genetic fate-mapping in Treg cells we introduced a *Gt(ROSA)26Sor^LSL-YFP^* recombination reporter allele into *Il10^tdTomato-CreER^Foxp3^Thy1.1^* mice (hereafter referred to as *Il10^FM^* mice). This enabled fate-mapping through activation of the Cre recombinase in IL-10 expressing cells by punctual tamoxifen administration, inducing their irreversible marking by YFP expression ^34^. Using *Il10^FM^* mice, we sought to identify Treg cells which were expressing IL-10 at the time of tamoxifen treatment and maintained (*Il10^stable^*) or lost IL-10 expression (*Il10^transient^*), acquired IL-10 expression after tamoxifen administration (*Il10^recent^*), or were not expressing IL-10 at the time of tamoxifen treatment or analysis (*Il10^neg^*, Fig. 1b). Analysis of YFP and tdTomato expression at different time points after tamoxifen administration revealed that all IL-10 expressing Treg cells in the colon stably maintained IL-10 expression, i.e. all tagged YFP^+^ cells continued to be tdTomato^+^, over at least 8 weeks (Fig. 1c). This was in contrast to cells from tissues other than the colon and from the colon draining mesenteric lymph node, where variable proportions of tagged YFP^+^ cells lost IL-10 expression (Fig. 1c). Even though they underwent gradual turnover, colonic *Il10^stable^* Treg cells were long lived, with a sizable population of YFP^+^ cells persisting at least 8 weeks after tamoxifen-induced tagging (Fig. 1d). Analysis of other major IL-10 expressing cell types in the colon suggested that the observed stability was not an intrinsic feature of IL-10 expression in that tissue, as fate-mapped YFP^+^ plasma cells, macrophages, and conventional T cells were prone to lose IL-10 expression to varying degrees (Extended Data Fig. 1e).

The unique stability of IL-10 expression by colonic Treg cells, combined with their persistence, suggested that they might represent a specialized, terminally differentiated effector Treg cell population. In considering this notion, we entertained several possibilities. First, Treg cells in the colon may stably maintain IL-10 expression because the intestinal microbiota is a cause of constant stimulation for colon resident immune cells: a variable absent in other ‘microbe-poor’ tissues such as the lung ^35–37^. Thus, the unique phenotype of colonic Treg cells could be explained by the fact that these cells, unlike those in the lung, are exposed to chronic inflammatory stimuli and thus the apparent stability of IL-10 expression reflects ongoing albeit inherently reversible activation of these cells rather than their terminal differentiation. To address this possibility, we induced chronic inflammation in the lungs of *Il10^FM^* mice by intranasal bleomycin challenge. In order to assess the stability of IL-10 expression by Treg cells in the presence of this sustained inflammatory stimulation, *Il10^FM^* mice were treated with tamoxifen 15 days after bleomycin challenge to ‘fate-map’ IL-10 expressing cells and analyzed for stability of IL-10 expression 21 days later (Fig. 1e). Although bleomycin challenge somewhat increased the proportion of Treg cells expressing IL-10, they were still prone to lose IL-10 expression even if to a lesser extent (Fig. 1f).

To directly test the role of the intestinal microbiota in driving or maintaining the *Il10^stable^* state, we administered tamoxifen to *Il10^FM^* mice 3 days prior to treating them with broad-spectrum antibiotics in their drinking water (Fig. 1e). YFP^+^ Treg cells in the colons of antibiotic treated *Il10^FM^* mice maintained stable IL-10 expression comparably with cells in littermate controls receiving vehicle without antibiotics (Fig. 1f). These results suggest that *Il10^stable^* cells, once differentiated, largely maintain their phenotypic state even if the intestinal microbiota, which facilitates IL-10 expression in Treg cells, is depleted (data not shown). Finally, we examined whether IL-10 expressing Treg cells would maintain IL-10 expression during a severe inflammatory insult. To test this, we first labeled pre-existing IL-10 expressing cells by administering tamoxifen to *Il10^FM^* mice. 10 days later we induced colitis in these mice by providing DSS in the drinking water for 7 days and analyzed cells in the colonic lamina propria 8 days after DSS withdrawal (Fig. 1e). Despite pronounced inflammation and intestinal Treg cell expansion, pre-existing YFP^+^ Treg cells retained IL-10 expression. Together, these results suggest that rather than being a product of ongoing stimulation, *Il10^stable^* Treg cells in the colon represent a terminally differentiated cell state robust to environmental perturbations.

### Transcriptomic analysis of colonic Il10^stable^ Treg cells

To elucidate the distinguishing molecular features associated with durable IL-10 expression in colonic Treg cells, we performed RNA-seq analysis on *Il10^neg^*, *Il10^recent^*, and *Il10^stable^* Treg cells isolated from the colonic lamina propria of *Il10^FM^* mice 21 days after tamoxifen induced labeling. Differential gene expression analysis contrasting *Il10^neg^* and *Il10^stable^* cells revealed divergent expression of various immunomodulatory and tissue supportive mediators, suggesting that these Treg cell populations participate in distinct regulatory and physiological processes (Fig. 2a). This observation, combined with differential expression of genes encoding receptors for chemokines and other external stimuli, and molecules involved in intercellular and extracellular matrix interactions, overall suggested distinct specialized niches for the generation, residence, and function of colonic *Il10^neg^* and *Il10^stable^* cell populations (Fig. 2a). Indeed, on the basis of differential TF expression, these populations seemed to represent the previously described Helios^hi^ and Rorγ(t)^hi^ colonic Treg cells, respectively (Fig. 2b), which have been shown to expand in response to distinct cues and to contribute to the control of distinct types of immune responses ^38–41^.

**Fig. 2.**
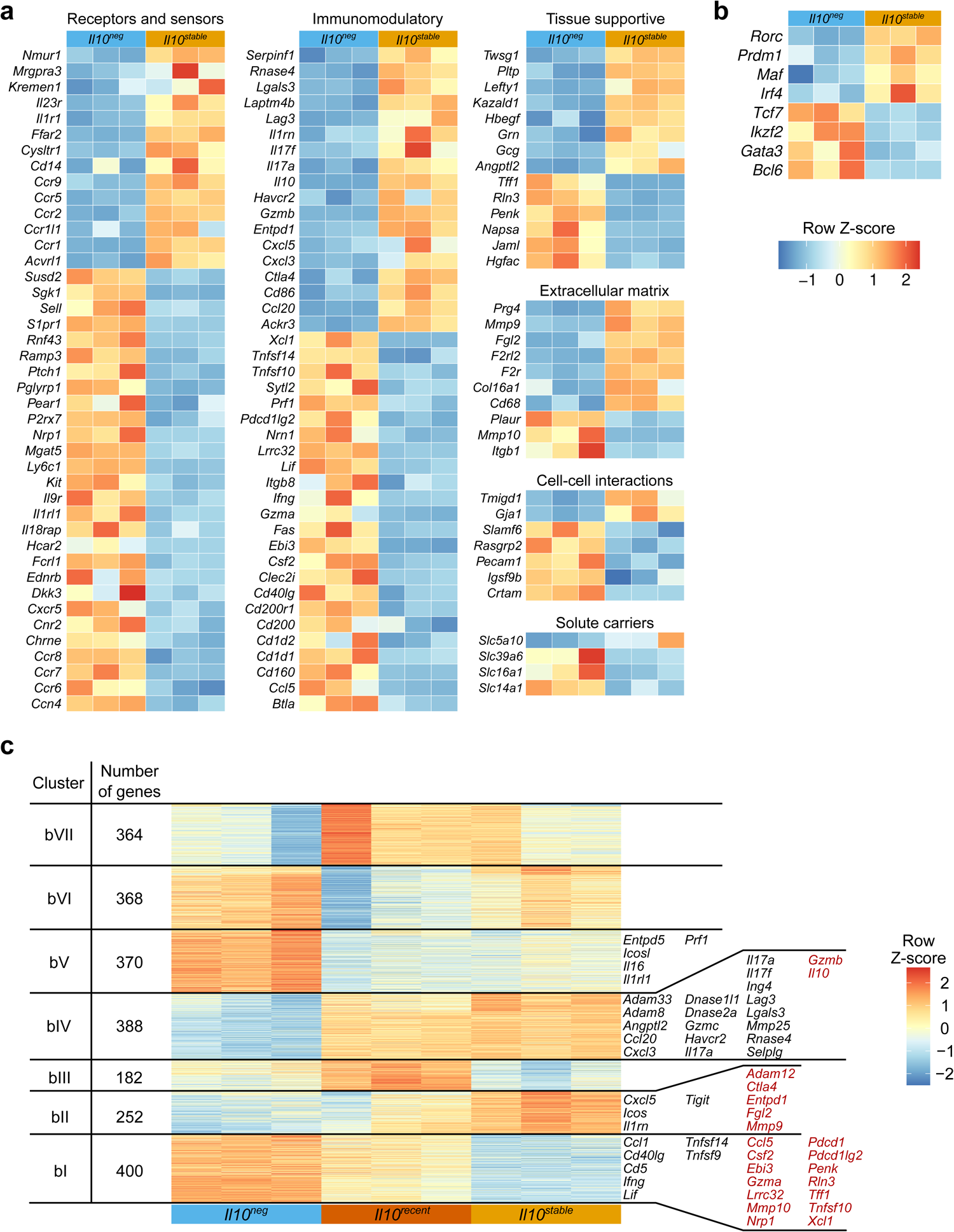
Distinct transcriptional features of *Il10^neg^* and *Il10^stable^* cells. a-c. RNA was isolated from *Il10^neg^*, *Il10^recent^*, and *Il10^stable^* Treg cells, as defined in Fig. 1b isolated from the colonic lamina propria of 10-week-old *Il10^FM^* mice treated 21 days prior with tamoxifen and sequenced. a, b. Coloring depicts Z-score normalized log_2_ transformed gene FPKM counts for individual *Il10^neg^* and *Il10^stable^* samples. Genes shown are all significantly differentially expressed between *Il10^neg^* and *Il10^stable^* (log_2_ FC > 1 and adjusted *p*-value < 0.05), annotated as encoding cell surface or secreted proteins, and manually categorized (a) or select differentially expressed (adjusted *p*-value < 0.05) TFs (b). c. K-means clustering was performed on Z-score normalized log_2_ transformed gene FPKM counts for genes significantly differentially expressed in any pairwise comparison (p-value < 0.05). Genes of interest differentially expressed between *Il10^stable^* versus *Il10^neg^* (black) or *Il10^stable^* versus *Il10^neg^* and *Il10^recent^* (red) within clusters I, II, IV, and V are indicated. See Methods for details. Negative binomial fitting with Wald’s significance test and the Benjamini & Hochberg correction for multiple comparisons. Significance testing and correction was performed on all genes.

To gain further insight into the differentiation process of *Il10^stable^* Treg cells, we incorporated into our analysis the *Il10^recent^* Treg cell population—cells that gained IL-10 expression only after the tamoxifen ‘time-stamping’ treatment 21 days prior—and performed single-cell RNA-seq (scRNA-seq) on colonic tdTomato^+^ and tdTomato^−^ Treg cells (Fig. 2c, 3a-e, 4a-c, and Extended Data Fig. 2a-c). Reasoning that precursors of these tissue Treg cells might be located in the draining lymph nodes, as suggested by recent reports, we also analyzed the corresponding Treg cells from the colonic mesenteric lymph node ^25,27^. First, we wished to reconcile our bulk RNA-seq and scRNA-seq analyses. We performed k-means clustering on the bulk RNA-seq data to identify gene expression clusters that best distinguish *Il10^neg^, Il10^recent^*, and *Il10^stable^* Treg cell populations (Fig. 2c). This revealed two gene clusters, one including the *Il10* gene itself, which increased in expression in *Il10^stable^* versus both *Il10^neg^* and *Il10^recent^* Treg cells (Fig. 2c, clusters bII and bIV). Conversely, many of the genes that were highly expressed in *Il10^neg^* cells lost expression in *Il10^stable^* versus *Il10^recent^* cells (Fig. 2c, cluster bI). Next, we mapped expression of the bulk RNA-seq gene (bI-VII) clusters to our scRNA-seq cell (sc0-10) clusters (Fig. 3c and Extended Data Fig. 2b-c). This analysis suggested the *Il10^stable^* cells were mostly present in sc0 and the small sc10 cluster, as this cell cluster had high expression of genes belonging to gene clusters bII and bIV (Fig. 3c and Extended Data Fig. 2b). Meanwhile, *Il10^neg^* cells seemed to comprise cell clusters sc1, sc2, sc3, and sc7 as these had the highest expression of bulk gene clusters bI, bV, and bVI (Fig. 3c and Extended Data Fig. 2b). Finally, *Il10^recent^* cells were apparently present within the cell cluster sc8, as these cells highly expressed genes from bulk clusters bIII and bVII, but also were present among the tdTomato^+^ cells in clusters sc1, sc2, and sc3. These cells, and bulk sequenced *Il10^recent^* cells both had intermediate expression of genes from bulk clusters bI and bII (Fig. 3c and Extended Data Fig. 2b). Altogether, this comparison of the scRNA-seq and bulk RNA-seq datasets pinpointed within the scRNA-seq dataset cell subsets corresponding to the sorted populations subjected to bulk RNA-seq analysis (Fig. 3d). Importantly, many of the bulk gene clusters showed statistically significant enrichment in the differentially highly expressed genes of each scRNA-seq subset used to identify cells as representing the *Il10^neg^*, *Il10^recent^*, or *Il10^stable^* bulk populations (Extended Data Fig. 2c).

**Fig. 3.**
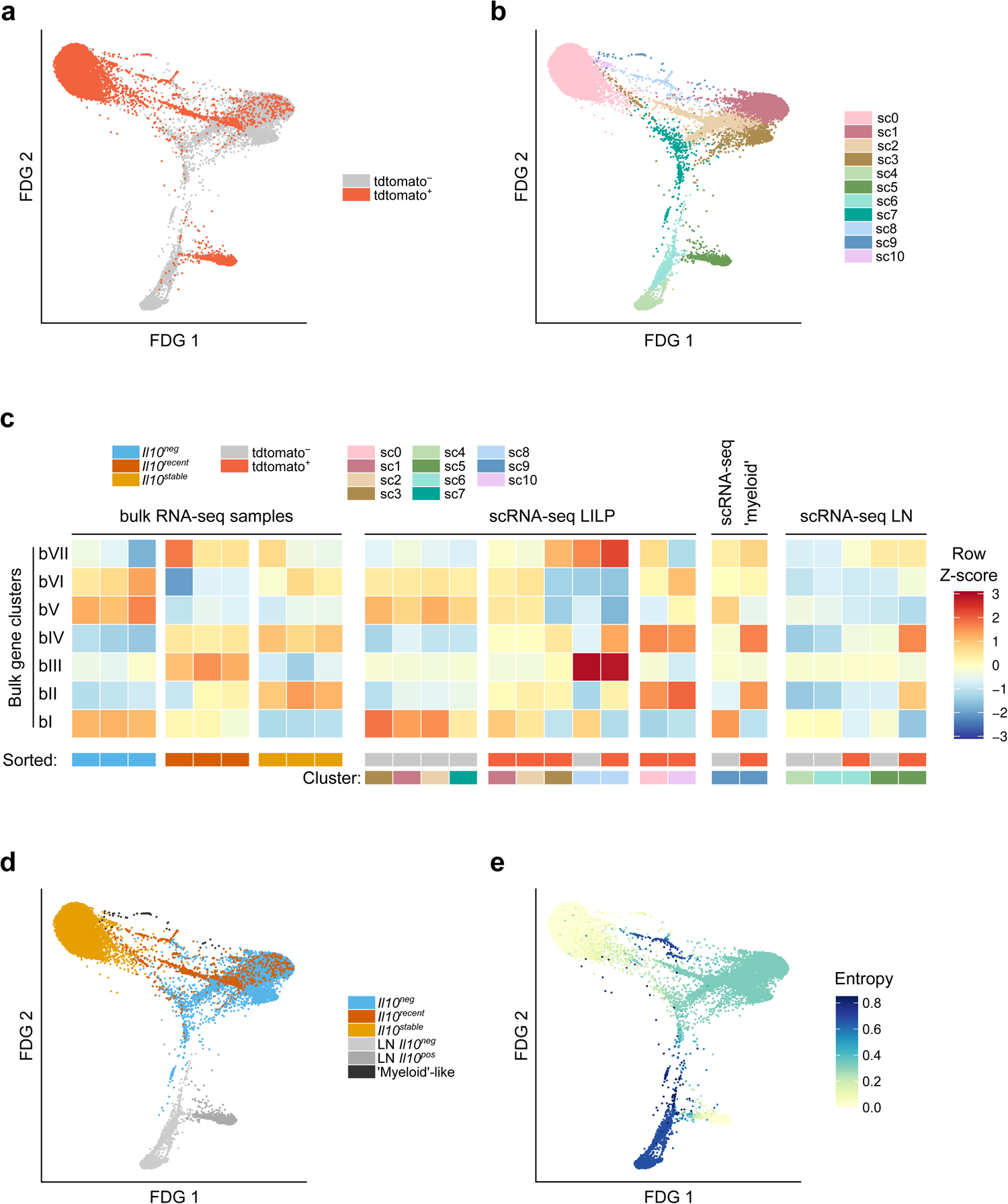
RNA-seq analysis of colonic Treg cells suggests terminal differentiation of *Il10^stable^*cells. a-e. tdTomato^+^ and tdTomato^−^ Treg cells (Thy1.1^+^CD4^+^TCRβ^+^) were separately sorted from the colon (LILP) and mesenteric lymph node (LN) of *Il10^FM^* mice and processed for scRNA-seq. See Methods for details. Additionally, bulk RNA-seq analysis was conducted on *Il10^neg^*, *Il10^recent^*, and *Il10^stable^* Treg cells, as defined in Fig. 1b, sorted from the colons of 10-week-old *Il10^FM^* mice treated 21 days prior with tamoxifen. a. 2D force-directed graph layout of tdTomato^+^ (red) and tdTomato^−^ (grey) Treg cells from the LILP and LN. b. Nearest-neighbor clustering was used to identify 11 distinct cell clusters. 2D force-directed graph layout of Treg cells from the LILP and LN, colored according to cluster. c. Integration of bulk and single-cell RNA-sequencing. Mean log_2_ transformed FPKM counts were computed for each k-means gene cluster for each bulk RNA-seq sample and mean expression was then Z-score normalized across samples per cluster. Bulk RNA-seq data are from RNA-seq analysis presented in Fig. 2 and 3. The scRNA-seq cells were scored for expression of bulk RNA-seq gene clusters, the 11 nearest-neighbor cell clusters were then separated as originating from the tdTomato^+^ or tdTomato^−^ sample, and manually organized. Per-subset expression score was Z-normalized across cell clusters per gene cluster. For clarity, scRNA-seq populations with very few cells are not depicted. Colored boxes indicate bulk RNA-seq samples sorted as *Il10^neg^*, *Il10^recent^*, or *Il10^stable^* Treg cell populations, and scRNA-seq populations by cluster and cell sample origin (tdTomato^+^ or tdTomato^−^). See Methods for details. d, e. Plots depicting 2D force-directed graph layout for all tdTomato^+^ and tdTomato^−^ LILP and LN scRNA-seq cells. Coloring indicates manually determined similarity of gene expression to the bulk-sorted *l10^neg^* (blue), *Il10^recent^* (orange), and *Il10^stable^* (yellow) Treg cell populations, or whether cells derive from the LN (light greys) or belong to a colonic ‘myeloid-like’ T cell population (dark grey) (d). Shading (cornsilk/low to indigo/high) indicates entropy, as determined by the Palantir algorithm (e). See Methods for details.

To explore potential developmental relations between cells belonging to these clusters, we used the Palantir algorithm, which assigns entropy measures to cells, indicative of differentiation potential ^42^. As a ‘starting cell’ for this analysis, we chose a random cell in cluster sc4, which was a cluster of lymph node cells enriched for markers of quiescence and naivete (*Lef1, Bcl2, Sell, S1pr1*). The relatively high entropy values in clusters sc1, sc2, sc3, and especially sc8 were consistent with the notion that these tdTomato^+^ cells might be in the process of differentiating into the *Il10^stable^* (sc0) cells (Fig. 3e). Interestingly, the colonic high entropy cluster (sc8) was enriched in cell cycle related genes, a feature also apparent in the bulk RNA-seq analysis of *Il10^recent^* cells (Fig. 4a-b). Altogether, these analyses suggested that within the colon, *Il10^recent^* cells underwent further differentiation into *Il10^stable^* cells and that this process was associated with cell division.

**Fig. 4.**
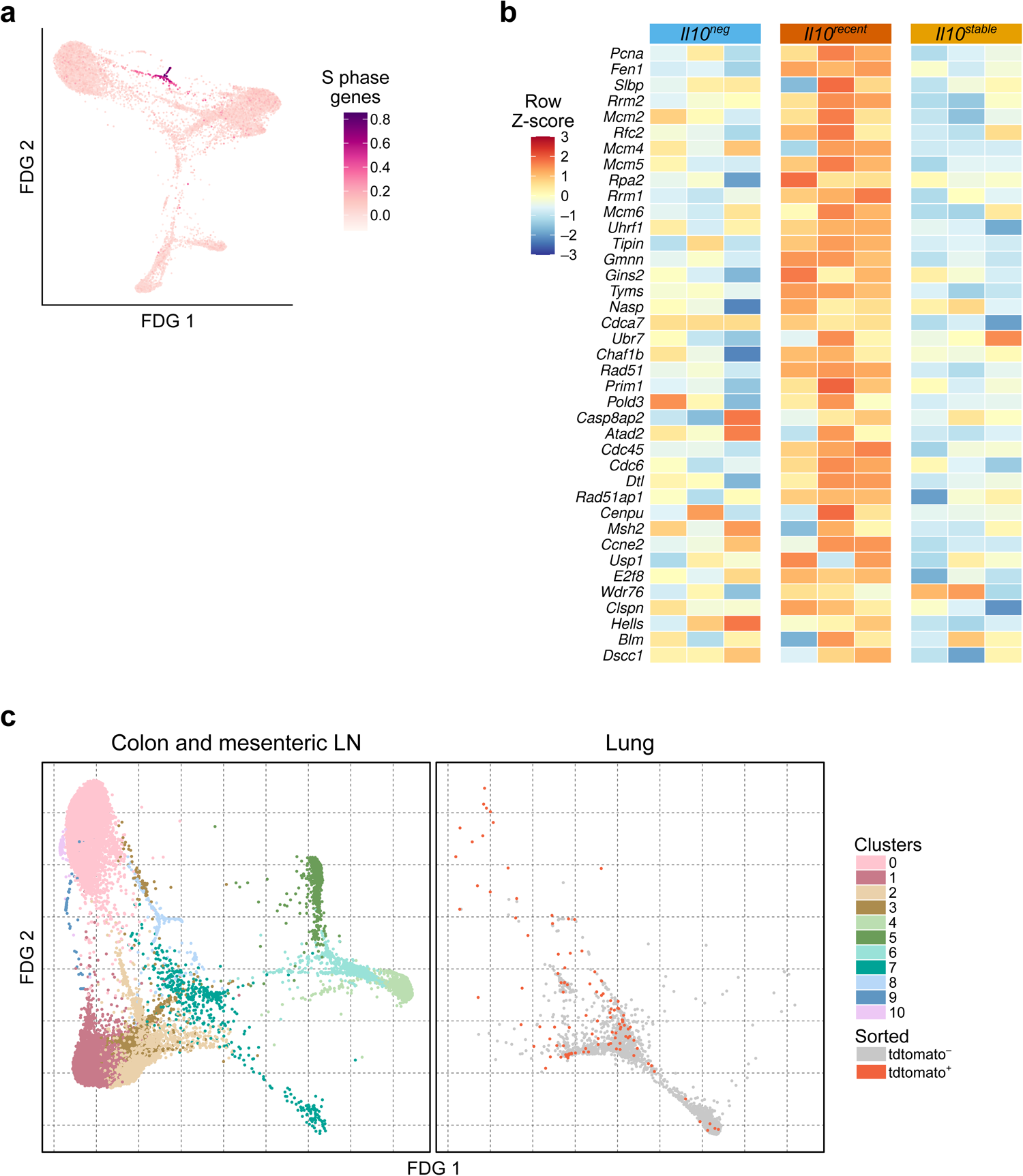
Characterization of IL-10 expressing and non-expressing Treg cells by scRNA-seq. a. tdTomato^+^ and tdTomato^−^ Treg cells (Thy1.1^+^CD4^+^TCRβ^+^) were separately sorted from the colon (LILP) and mesenteric lymph node (LN) of *Il10^FM^* mice and scRNA-seq was performed. 2D force-directed graph layout of all tdTomato^+^ and tdTomato^−^ LILP and LN scRNA-seq cells. Shading indicates gene expression score for genes associated with the S phase of the cell cycle. b. Data are from RNA-seq analysis presented in Fig. 2 and 3. Heatmap showing log_2_ transformed, row Z-score normalized FPKM counts for genes associated with the S phase of the cell cycle among bulk RNA-seq samples. c. tdTomato^+^ and tdTomato^−^ Treg cells (Thy1.1^+^CD4^+^TCRβ^+^) were separately sorted from the colon lamina propria (LILP), mesenteric lymph node (LN) and lung of *Il10^FM^* mice and scRNA-seq was performed. Data are from scRNA-seq analysis presented in Fig. 3 and Extended Data Fig. 2, with the addition of cells from the lung. 2D force-directed graph layout of all tdTomato^+^ and tdTomato^−^ scRNA-seq cells. Left: coloring indicates assigned nearest-neighbor cluster for each cell when clustering LILP and LN cells, as in Fig. 3a. Right: coloring indicates cells sorted as tdTomato^+^ or tdTomato^−^ from the lung.

Our fate-mapping analysis of IL-10 expressing Treg cells showed the existence of *Il10^transient^* cells in lungs (Fig. 1c). This suggested that in contrast to the colon, IL-10 expressing Treg cells in the lung do not undergo a similar terminal differentiation process. This notion was supported by scRNA-seq analysis, since both tdTomato^+^ and tdTomato^−^ Treg cells from the lung tended to cluster away from colonic Treg cells with the lowest entropy values, i.e. the most differentiated states. Moreover, both lung Treg cell populations tended to cluster together, suggesting their overall similarity despite the difference in IL-10 expression (Fig. 4c). The presence of a small number of lung tdTomato^+^ cells clustering with colonic cell cluster sc0 might be a reflection of the existence of a small population of terminally differentiated *Il10^stable^* cells in the lung, which may be generated directly in the lung, or result from migration of colonic *Il10^stable^* Treg cells or their immediate local precursors, a possibility suggested by a recent study of colonic Treg cell emigration^43^.

### Identification of transcriptional programs active in colonic Treg cells

Next, we sought to identify the transcriptional regulators and corresponding upstream signaling pathways controlling the differentiation of *Il10^stable^* colonic Treg cells by performing ATAC-seq on the same populations subjected to bulk RNA-seq analysis: *Il10^stable^*, *Il10^recent^*, and *Il10^neg^* colonic Treg cells isolated 21 days after tamoxifen induced labeling. Many of the differentially accessible peaks in the ATAC-seq atlas which distinguished *Il10^stable^* from *Il10^neg^* Treg cells were also similarly differentially accessible between *Il10^stable^* and *Il10^recent^* populations (Pearson’s r: 0.602, p-value: < 2.2 × 10^−16^), suggesting that identifying the pathways and transcription factors (TFs) converging on these peaks might reveal which TFs differentially specify the *Il10^stable^* versus *Il10^neg^* state, as well as TFs facilitating the differentiation of *Il10^recent^* into *Il10^stable^* Treg cells (Fig. 5a).

**Fig. 5.**
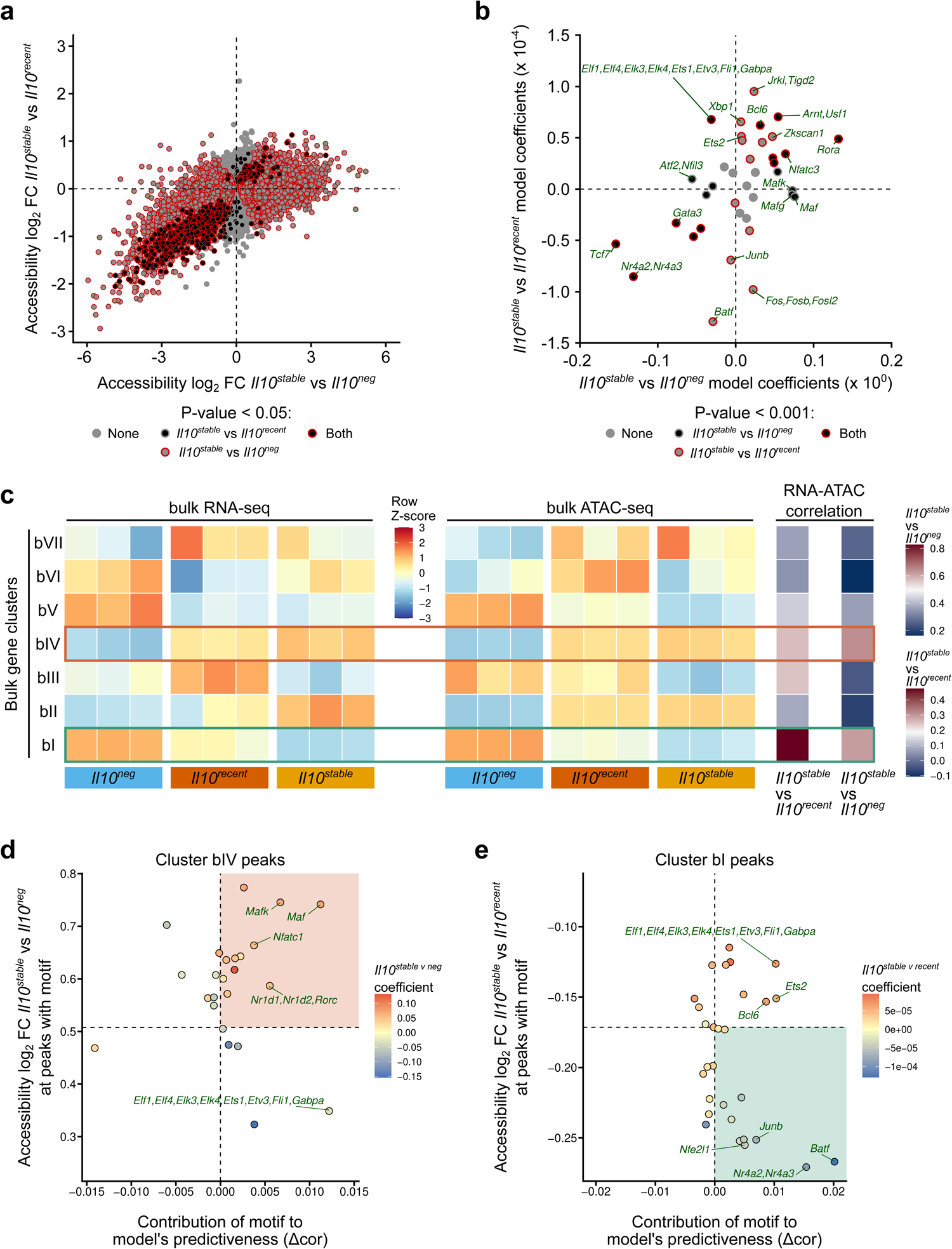
ATAC-seq analysis of TF motifs associated with differential chromatin accessibility in colonic IL-10 expressing and non-expressing Treg cell subsets. *Il10^neg^*, *Il10^recent^*, and *Il10^stable^* Treg cells, as defined in Fig. 1b, were sorted from the LILP of 10-week-old *Il10^FM^* mice treated 21 days prior with tamoxifen and subjected to ATAC-seq analysis. See Methods for details. a. Plot showing mean log_2_ fold change (FC) peak accessibility for *Il10^stable^* versus *Il10^neg^* (x-axis) and *Il10^stable^* versus *Il10^recent^* (y-axis) samples. Coloring indicates peaks significantly differentially accessible (adjusted *p*-value < 0.05) in neither comparison (grey), between *Il10^stable^* versus *Il10^neg^* samples (red outline), between *Il10^stable^* versus *Il10^recent^* samples (black fill), or in both comparisons (black fill with red outline). Negative binomial fitting with Wald’s significance test and the Benjamini & Hochberg correction for multiple comparisons. Significance testing and correction was performed on all peaks. b. Linear ridge regression models generated for *Il10^stable^* versus *Il10^neg^* (svn) and *Il10^stable^* versus *Il10^recent^* (svr) chromatin accessibility changes as a function of motif presence/absence in all peaks. Plots showing per-motif coefficients for the svn (y-axis) and svr (x-axis) models and coloring indicates motifs with significantly contributing (*p*-value < 0.001) coefficients: in neither model (grey), in the svn model (black fill), in the svr model (red outline), or in both models (black fill with red outline). Motifs with coefficients > mean ± SD and *p*-values < 0.001 are labeled. See Methods for details. c. Mean log_2_ transformed FPKM counts were computed for each k-means gene cluster for each bulk RNA-seq sample (Fig. 2 and 3). Mean expression was then Z-score normalized across samples per cluster. Mean log_2_ transformed FPKM accessibility counts of peaks associated with genes in each k-means cluster were calculated per ATAC-seq sample. Mean accessibility was Z-score normalized across samples per cluster. Shading (blue/low – yellow/mid – red/high) indicates Z-score normalized expression or accessibility count means. Pearson’s correlation coefficients were calculated for expression FC versus accessibility FC for the genes and associated peaks in each cluster, for the svn and svr comparisons. Shading (blue/low – white/mid – red/high) indicates correlation coefficient, centered on the correlation coefficient for the genome-wide gene expression and accessibility linear regression for each comparison. Clusters with the highest correlation for comparison (green – Cluster bI – svr, orange – Cluster bIV – svn) are identified by boxes. d, e. Correlation coefficients for predicted versus actual log_2_ FC peak accessibility for the svn (d) and svr (e) models at peaks associated with cluster bIV (d) and cluster bI (e) genes were determined. Corresponding correlation coefficients were then determined for models with each motif individually withheld. X-axes show difference between original correlation coefficients for predicted versus actually log_2_ FC and motif withheld correlation coefficient (Δcor). Y-axes show log_2_ FC peak accessibility for cluster bIV (d) and cluster bI (e) associated peaks containing each motif. Horizontal dashed line indicates log_2_ FC peak accessibility for all cluster bIV (d) and cluster bI (e) associated peaks. Shading indicates per-motif coefficients for the svn (d) and svr (e) models. Motifs with Δcor > 0 + SD, or top 3 motifs with the highest Δcor are labeled. Quadrants with motifs positively contributing to the model’s predictiveness and associated with above-average increased (d) or decreased (e) accessibility are highlighted in orange (d) or green (e).

To do this, we first identified motifs within each peak in our peak atlas corresponding to TFs expressed in any of the three cell populations. We then fit separate linear ridge regressions on this peak by motif matrix to generate models that would predict accessibility differences at each peak in both the *Il10^stable^* versus *Il10^neg^* (svn model) and *Il10^stable^* versus *Il10^recent^* (svr model) comparisons ^44–46^. Each unique motif is represented by a term in these models, whose coefficient indicates to what extent that motif contributes to predicting accessibility changes. Thus, large positive or negative coefficient values suggest that the presence of a given motif is more strongly associated with increased or decreased accessibility at peaks harboring that motif, ultimately implying increased or decreased activity of the corresponding TFs at those loci in the analyzed cell populations. Comparing the coefficients from the two models revealed motifs associated with accessibility changes in a statistically significant manner in the svn model, the svr model, or both (Fig. 5b and data not shown). Coefficients for motifs for TFs differentially expressed between *Il10^neg^* and *Il10^stable^* cells, such as Gata3 and Maf corresponded with their differential expression, suggesting biologically meaningful changes in TF activity revealed by this analysis (Fig. 2b & 5b). The TF Blimp1 (encoded by the *Prdm1* gene) antagonizes the repressive activity of Bcl6; therefore, increased Blimp1 activity would manifest in increased chromatin accessibility at Bcl6 target genes. Accordingly, we found an enrichment of Bcl6 motifs at sites characterized by increased accessibility in *Il10^stable^* cells, consistent with the increased expression of Blimp1 in *Il10^stable^* versus *Il10^neg^* cells (Fig. 2b). Notably, motifs of TFs which act in T cells as downstream effectors of TCR stimulation (AP-1 family members Junb, Fos, Fosb, Fosl2; Batf; and Nr4a2/3) were associated with a loss of accessibility in *Il10^stable^* relative to *Il10^recent^* cells, as well as to *Il10^neg^* cells (Fig. 5b). This suggested that attenuation of TCR stimulation or signaling was associated with the terminal differentiation of Treg cells into the *Il10^stable^* state.

We focused our subsequent analysis on the bulk RNA-seq gene clusters bI and bIV, which were of particular interest as differential expression of the constitutive genes continued to increase (bIV) or decrease (bI) alongside sustained *Il10* gene expression (Fig. 5c). Thus, expression of these genes was specifically gained or lost, respectively, during terminal differentiation of IL-10 expressing Treg cells. A similar pattern was observed for differential chromatin accessibility of the *cis*-regulatory elements (ATAC-seq peaks) associated with these genes (Fig. 5c). Therefore, we reasoned that elucidation of TF motifs associated with chromatin accessibility changes at these peaks might reveal candidate TFs whose differential binding contributed to regulation of gene expression at those loci. Additionally, chromatin accessibility changes at these subsets of peaks had stronger correlation with gene expression changes in both the *Il10^stable^* versus *Il10^neg^* and *Il10^stable^* versus *Il10^recent^* comparisons than in the overall ATAC-seq peak atlas, further supporting that specific interrogation of these peaks would prove informative (Fig. 5c). We wanted to use our models to more specifically elucidate which motifs associated with accessibility changes at the genes of these clusters bI and bIV. We reasoned that comparing the performance of the models in predicting accessibility differences at specific peaks with individual motifs removed might reveal which motifs were specifically associated with increased or decreased accessibility at those peaks. For example, if removing a motif X from the model decreased its performance, i.e. the correlation between predicted and actual accessibility fold change values, at peaks belonging to cluster Y, this would suggest that TFs recognizing motif X are specifically active at cluster Y peaks. This analysis for cluster bIV peaks in the svn model associated Nfatc1, Rorc, and Maf family TF motifs with increased accessibility at cluster bIV peaks in *Il10^stable^* cells (Fig. 5d). For cluster bI peaks, the motifs of TCR responsive TFs Nr4a2/3, Batf, and AP-1 were associated with loss of accessibility in *Il10^stable^* cells relative to less-differentiated *Il10^recent^* cells (Fig. 5e). Reassuringly, peaks within each cluster with the relevant motifs had greater magnitude accessibility differences than all peaks in the cluster (Fig. 5d-e, y-axes), consistent with the notion that the corresponding TFs contributed to these accessibility changes. Indeed, cluster bIV gene loci such as *Il10* itself showed multiple peaks containing Maf motifs gaining accessibility alongside increased gene expression, while cluster bI loci such as *Ly75* showed decreased gene expression and accessibility at peaks containing Nr4a2/3 motifs (Extended Data Fig. 3A-b).

A hierarchical arrangement of Maf, Rorγt, Blimp1 activities in IL-10 induction in colonic Treg cells has been established ^23,47–51^. Thus, our epigenetic analysis seems to confirm a role for at least two of these TFs in inducing IL-10 expression and suggests they modulate expression of other co-regulated effector molecules. However, it has remained unknown whether these TFs are required to maintain IL-10 expression, let alone a larger gene expression program, in differentiated cells. Interestingly, our data comparing *Il10^recent^* and *Il10^stable^* Treg cell populations did not indicate discernable role for either Maf or Rorγt in the latter after their differentiation suggesting that these TFs might in fact be dispensable for maintaining this Treg cell state in general, and IL-10 expression in particular (Fig. 5b; also, note the lack of a significant Δcor for either Rorc or Maf motifs in Fig. 5e). We sought to formally test this possibility using both loss- and gain-of-function strategies. First, we ablated conditional *Maf* and *Rorc* alleles in IL-10 expressing cells in *Il10^tdTomato-CreER^Maf^FL^* and *Il10^tdTomato-CreER^Rorc^FL^* mice by treating them with tamoxifen (Extended Data Fig. 4a & 4b). Induced loss of these TFs in Treg cells, which had already acquired IL-10 expression, did not lead to substantial impairment of its maintenance, confirming the dispensability of these TFs for maintaining IL-10 expression (Extended Data Fig. 4a & 4b). Second, we inducibly expressed these TFs in *in vitro* activated Treg cells using retroviral gene transfer. The enforced expression of c-Maf and RORγt did not promote increased persistence of IL-10 expression, reinforcing the notion that these TFs play a non-redundant role only in the induction of IL-10 expression by Treg cells (Extended Data Fig. 4c & 4d). The higher per cell expression of IL-10 in Maf over-expressing cells is consistent with the known role of c-Maf in directly activating the *Il10* gene, likely through a motif in the *Il10* promoter identified in our ATAC-seq analysis (Extended Data Fig. 3a & 4c). At the same time, the effect of enforced Maf expression in causing loss of IL-10 expression in a small proportion of cells may reflect an unexpectedly nuanced role for this TF in regulating IL-10 that merits further study (Extended Data Fig. 4d).

### Altered TCR signaling and diminished TCR dependence in long-lived Il10^stable^ cells

The data thus far suggested that the transcriptional and epigenetic effects downstream of TCR signaling were distinct among different colonic Treg cell subsets, and that IL-10 expressing colonic Treg cells undergoing terminal differentiation were distinguished by progressively diminished TCR signaling. Supporting this notion, expression of a set of genes previously characterized as TCR activated was overall decreased, while expression of a TCR repressed gene set was overall increased in *Il10^stable^* versus *Il10^recent^* cells ^52^ (Extended Data Fig. 5a). While the TCR activated gene set also had decreased expression in *Il10^stable^* versus *Il10^neg^* cells, the TCR repressed genes did as well, suggesting that TCR signaling is somewhat attenuated in both *Il10^stable^* and *Il10^neg^* Treg cells, although in qualitatively distinct ways (Extended Data Fig. 5b). This latter possibility was consistent with the observed association of Nfatc1 motifs, contrary to other TCR signaling-dependent TF motifs, with increased accessibility of peaks at cluster IV genes in *Il10^stable^* versus *Il10^neg^* cells. Genes encoding proteins that propagate or amplify TCR signaling cascades were differentially expressed among *Il10^neg^*, *Il10^recent^*, and *Il10^stable^* cells, with the majority having decreased expression in the two IL-10 expressing Treg cell populations (Extended Data Fig. 5c). Conversely, several negative regulators of TCR signaling were highly expressed in *Il10^stable^* cells, exemplified by significantly higher expression of *Ubash3b* in *Il10^stable^* versus *Il10^recent^* cells (Extended Data Fig. 5d). Finally, components of the TCR complex such as *Cd247, Cd3e*, and *Tcrb,* and the co-receptor *Cd4*, were more highly expressed in *Il10^stable^* versus *Il10^neg^* Treg cells, with *Tcrb* being significantly increased in *Il10^stable^* versus *Il10^pos^* cells as well (Extended Data Fig. 5e). Given that these genes are transcriptionally repressed with TCR stimulation, this observation was also consistent with the posited attenuated TCR signaling in *Il10^stable^* cells ^53^.

These observations raised the possibility that the effector function of *Il10^stable^* Treg cells might be TCR independent and furthermore, that loss of the TCR would not diminish and might even increase the proportion of effector IL-10 expressing Treg cells undergoing terminal differentiation. We directly tested these possibilities through acute inducible ablation of the TCR in IL-10 expressing cells in *Il10^tdTomato-CreER^Trac^FL/FL^* (*Il10^iΔTCR^*) mice. Tamoxifen induced deletion of the conditional *Trac* allele leads to loss of the TCRα chain, and therefore, of the entire TCR signaling complex from the cell surface, enabling for identification of TCR-deleted cells by flow cytometric analysis of TCRβ cell surface expression ^54,55^ (Fig. 6b). These mice also harbored the *Gt(ROSA)26Sor^LSL-YFP^* recombination reporter allele, allowing us to track IL-10 tagged cells that lost or retained the TCR over time. We treated *Il10^iΔTCR^* mice with tamoxifen and analyzed YFP^+^ colonic Treg cells ten days later (Fig. 6a). For various molecules whose expression was enriched in the *Il10^stable^* Treg cells, we saw an unchanged or even increased proportion of cells expressing these markers among cells losing TCR expression (Fig. 6c-d). This included IL-10 (tdTomato) itself, as well as another immunomodulatory molecule CD39 (*Entpd1*), in addition to molecules involved in cell localization and tissue residency (CD69 and CCR9). However, this pattern was not universal, as the expression of several proteins encoded by transcripts highly expressed in *Il10^stable^* cells was TCR-dependent, such as CD25 (*Il2ra*) and CCR5 (*Ccr5*) (Fig. 6c-d). Finally, expression of proteins encoded by genes associated with the *Il10^neg^* Treg cell population, such as GITR (*Tnfrsf18*) and KLRG1 (*Klrg1*), were further diminished in IL-10 expressing cells after loss of the TCR (Fig. 6c-d). Overall, these data support the notion that selective diminution of or withdrawal from TCR signaling supports the terminal differentiation of *Il10^stable^* Treg cells and that conversely, ongoing TCR stimulation maintains some phenotypic plasticity of Treg cells to acquire distinct terminal differentiation fates.

**Fig. 6.**
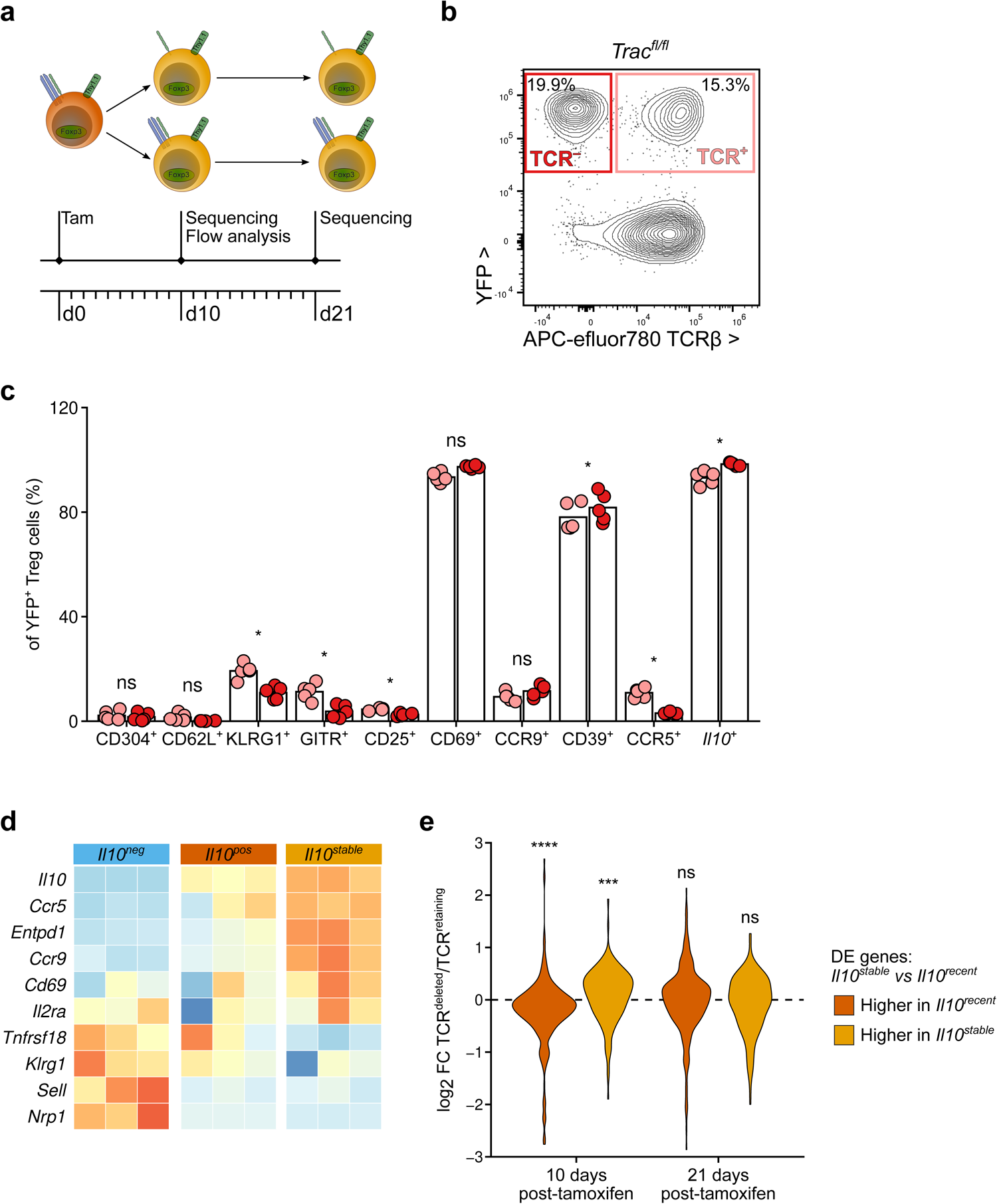
Features of *Il10^stable^* cells maintained in a TCR independent manner. a. Experimental schematic. 10-week-old male and female *Il10^iΔTCR^* mice were treated with tamoxifen on day 0 and cells were isolated from the LILP on day 10 for flow cytometric analysis (c) or TCR sufficient and deficient Treg cells were sorted on days 10 and 21 after tamoxifen administration and subjected to RNA-seq analysis (e). b. Representative 2D flow cytometry plots pre-gated on Treg cells (Thy1.1^+^CD4^+^CD90^+^CD5^+^) from the LILP of *Il10^iΔTCR^* mice. Plots show YFP expression (y-axis) and TCRβ cell surface expression (x-axis). c. Frequencies of cells positive for the indicated molecules among TCR sufficient (pink) or TCR deficient (red) YFP^+^ Treg cells isolated from the LILP of *Il10^iΔTCR^* mice. Paired two-sided t-tests corrected for multiple comparisons using the FDR method of Benjamini & Hochberg. Each point represents an individual mouse (n = 5, with TCR-sufficient and -deficient cells in each mouse) and represent data pooled from two independent experiments. d. Row Z-score normalized log_2_ expression across *Il10^neg^*, *Il10^recent^*, and *Il10^stable^* Treg cell samples for genes encoding the molecules assessed in (c). Data are from RNA-seq analysis presented in Fig. 2 and 3. e. Violin plots of log_2_ transformed gene expression fold change for TCR sufficient versus deficient samples at 10 or 21 days post-tamoxifen treatment. Genes differentially expressed between *Il10^stable^* and *Il10^recent^* Treg cell populations are divided into those with significantly increased expression in *Il10^stable^* Treg cells (yellow) or *Il10^recent^* cells (orange). *P*-values calculated by two-sided Kolmogorov-Smirnov test for log_2_ FC of genes with significantly increased expression in *Il10^stable^* Treg cells (yellow) or *Il10^recent^* cells (orange) versus of all genes are indicated. *P*-value > 0.05 ns, < 0.05 *, < 0.001 ***, < 0.0001 ****

To characterize more broadly the changes caused by TCR ablation in *Il10^stable^* colonic Treg cells, we performed RNA-seq analysis of TCR-sufficient and -deficient YFP^+^ Treg cells isolated from the colons of *Il10^iΔTCR^* mice 10 and 21 days after tamoxifen induced labeling. We assessed the expression of genes differentially expressed in *Il10^stable^* versus *Il10^recent^* Treg cells, i.e. genes with increasing or decreasing expression during the differentiation of *Il10^stable^* cells. Overall, the latter genes, those more highly expressed in *Il10^recent^* Treg cells, showed lower expression in TCR-deficient versus TCR-sufficient YFP^+^ Treg cells, whereas those which were increased in expression in *Il10^stable^* cells had overall higher expression in TCR deficient cells (Fig. 6e). Of note, this effect was more pronounced at day 10 rather than day 21 post TCR deletion, suggesting that at the latter timepoint, even the YFP “time-stamped” TCR sufficient cells had largely completed their terminal differentiation (Fig. 6e). Overall, this was consistent with the notion that terminal differentiation of IL-10 expressing effector Treg cells was constrained by ongoing TCR signaling and that its diminution or even withdrawal from it facilitated this process.

### Role of colonic Il10^stable^ Treg cells in preventing spontaneous inflammation

In contrast to multi-potential effector conventional T cells and their “stem-like” source cells, terminally differentiated effector T cells are often characterized as exhausted or dysfunctional: contributing minimally to clearing infectious agents, controlling tumor progression, or causing autoimmune disease ^56–59^. However, support for this characterization largely comes from studies of CD8^+^ T cells, and it remains unclear whether this conception extends to Treg cells. Given the apparent terminal differentiation of colonic *Il10^stable^* Treg cells, we wondered whether and to what extent these cells remain functional and contribute to immune regulation in this tissue. Considering the high and sustained expression of IL-10 by this population, and the known role of Treg cell derived IL-10 in preventing spontaneous colitis, one could assume that *Il10^stable^* Treg cells might be an important source of this cytokine in this context ^32,60^. However, previous studies of the role of Treg cell derived IL-10 using *Foxp3^Cre^Il10^FL/FL^* mice relied on constitutive ablation of IL-10 expression from Treg cells throughout the lifespan of the mice, including the critical early life period of microbial colonization. Therefore, the spontaneous colitis reported in these animals might be attributed to a requirement for Treg cell derived IL-10 during a critical developmental window, rather than its continuous production by a specialized colonic Treg cell population. Indeed, studies describing the unique contribution of Treg cells during the time of weaning and microbial community assembly support the former notion ^61,62^.

Therefore, we sought to assess the role of ongoing IL-10 production by Treg cells using tamoxifen inducible IL-10 ablation in healthy adult *Foxp3^CreER^Il10^FL^* mice. *Foxp3^CreER^Il10^FL/KO^* and littermate control *Foxp3^CreER^Il10^WT/KO^* mice were treated with two sequential doses of tamoxifen to achieve pronounced loss of IL-10 secretion by Treg cells and analyzed 16 days later (Extended Data Fig. 6ab and 7a). Unexpectedly, mice depleted of Treg cell derived IL-10 showed no signs of inflammatory disease, as assessed by weight loss or colon shortening (Extended Data Fig. 7b-c). This observation suggested that continuous production of IL-10 by Treg cells in the colon was dispensable for preventing colonic inflammation in lymphoreplete adult mice and appeared to support the notion that terminally differentiated colonic *Il10^stable^* Treg cells were non-functional, or at least did not play a non-redundant role in local immune regulation. Confirming that this observed dispensability of Treg cell derived IL-10 in adult animals was not simply due to the short duration of the experiment, no changes in colon length were observed upon extended IL-10 ablation in Treg cells for 5 weeks (Extended Data Fig. 6c & 6d).

This reasoning relies on the assumption that IL-10 production was the dominant effector modality of colonic *Il10^stable^* Treg cells. However, based on their gene expression program, IL-10 could constitute just one of multiple, potentially redundant immune regulatory mechanisms deployed by these cells (Fig. 2a). Thus, targeting these cells rather than just their production of IL-10 was required for proper testing of their function. We therefore developed a novel genetic model for selective ablation of IL-10 expressing Treg cells by engineering a modified *Foxp3* knock-in allele harboring a loxP-STOP-loxP (LSL) cassette upstream of the coding sequence for the human diphtheria toxin receptor (DTR) in the 3’ untranslated region (Extended Data Fig. 8a). In these *Foxp3^LSL-DTR^* mice, DTR expression by Foxp3^+^ Treg cells is conditional on expression of a Cre recombinase from a locus of interest. We therefore generated *Il10^tdTomato-Cre^Foxp3^LSL-DTR^* (*Il10^Foxp3-DTR^*) mice to allow for the specific depletion of IL-10 expressing Treg cells by diphtheria toxin (DT) administration and confirmed that treating mice with DT, but not heat-inactivated DT (boiled, bDT) resulted in loss of IL-10 expressing Treg cells while sparing other IL-10 expressing populations and other Foxp3^+^ cells, i.e *Il10^neg^* cells (Extended Data Fig. 8b and 8c). Notably, transient depletion of IL-10 expressing Treg cells in *Il10^Foxp3-DTR^* mice upon punctual administration of DT followed by a brief recovery revealed that these cells quickly repopulated the colon, suggesting that any progenitor population was spared by our depletion strategy and that it enabled specifically probing the function of *bona fide* terminally differentiated cells (Extended Data Fig. 8d and 8e).

In contrast to the loss of IL-10 production by Treg cells, depletion of IL-10 expressing Treg cells by DT treatment of *Il10^Foxp3-DTR^* mice for 16 days resulted in significant weight loss and colon shortening versus bDT treated controls (Fig. 7a-c). This was accompanied by increased frequencies of natural killer (NK) cells, neutrophils, and especially eosinophils in the colonic lamina propria (Fig. 7d-f). Increased abundance of the inflammatory chemokines responsible for recruiting these innate effector populations, CXCL9/10, CXCL1, and CCL11, suggested that loss of IL-10 expressing Treg cells resulted in a broadly heightened inflammatory state of the colon (Fig. 7g-j). Importantly, neither the inflammatory cell populations nor the pro-inflammatory chemoattractants were increased in the absence of Treg cell derived IL-10, suggesting that the *Il10^stable^* Treg cell population deployed additional functionally important immunoregulatory mechanisms alongside IL-10 (Extended Data Fig. 7d-j). Collectively, these results suggested that rather than being dysfunctional, IL-10 expressing Treg cells represented a specialized population of immunomodulatory effectors essential for preventing spontaneous inflammation in the colon.

**Fig. 7.**
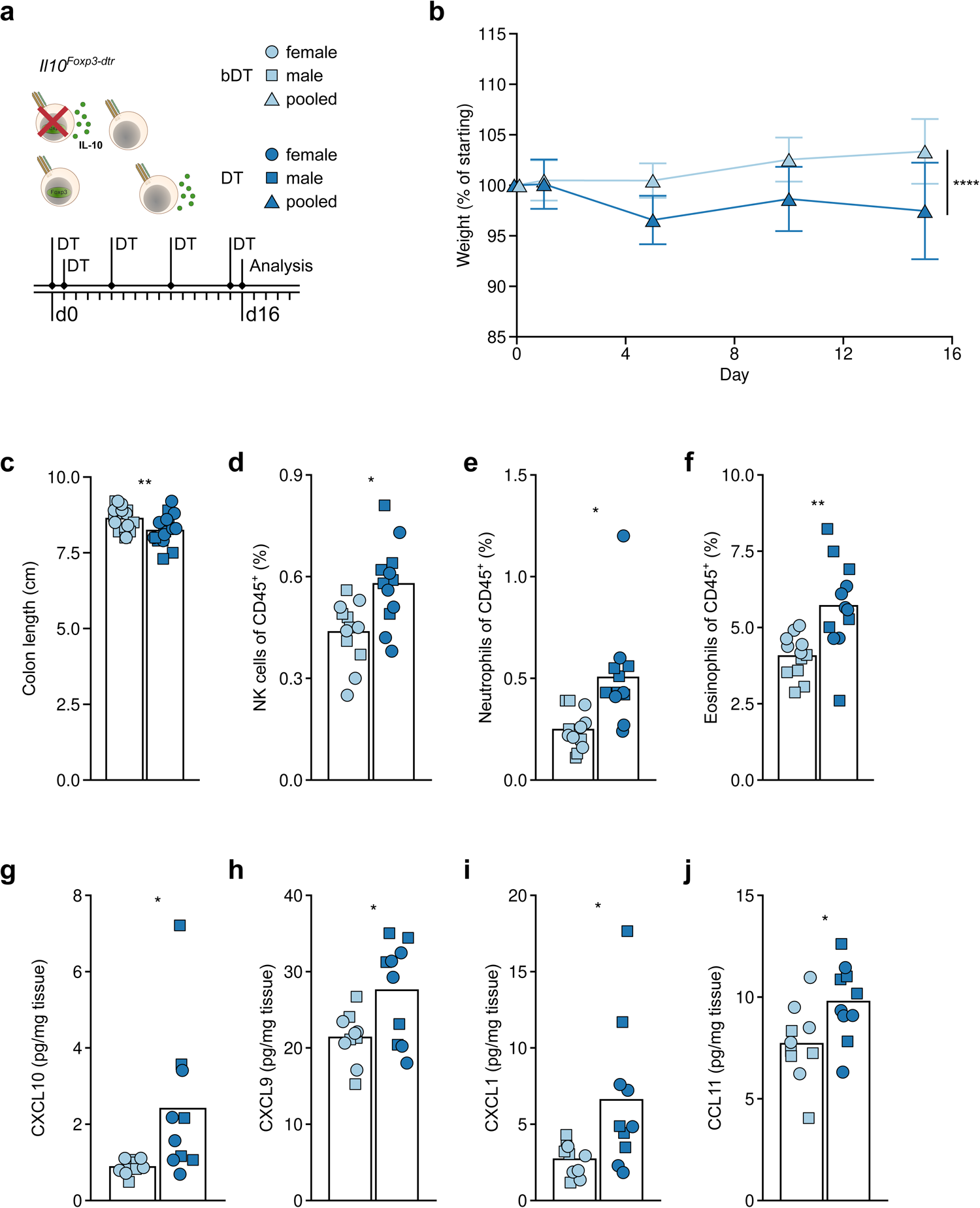
Acute ablation of *Il10* expressing Treg cells results in spontaneous colonic inflammation. 8- to 11-week-old *Il10^tdTomato-Cre^Foxp3^LSL-DTR^* (*Il10^Foxp3-DTR^*) mice were treated with active (DT) or heat-inactivated (boiled, bDT) diphtheria toxin over 16 days before analysis by flow cytometry. a. Diagram depicting targeted and punctual DT-induced ablation of IL-10 expressing Treg cells in *Il10^Foxp3-DTR^* mice (top left). Legend for plots throughout the figure with DT and bDT groups colored as light blue or dark blue, respectively, and female or male mice depicted as circles or squares, respectively; data representing pooled male/female samples are depicted as triangles (top right). Experimental treatment regimen timeline (bottom). b. Plot showing body weights of DT and bDT treated groups over the time course of the experiment. Weights are normalized to starting body weight for each mouse. Plots show mean ± SD and represent data pooled from five independent experiments (n = 3-4 mice per experiment per group, with n = 19 for each group overall and with each mouse weighed repeatedly for each time point). ANOVA for weight change as a function of treatment, sex, experimental replicate, and time with *p*-value for the effect of treatment determined by Tukey’s ‘Honest Significant Difference’ method. c. Plot showing colon lengths on day 16 of DT and bDT treated mice. Each point represents an individual mouse (n = 19 for each group) and data shown are pooled from five independent experiments. ANOVA for colon length as a function of treatment, sex, and experimental replicate with *p*-value for the effect of treatment determined by Tukey’s ‘Honest Significant Difference’ method. d-f. Cellular composition of the colonic lamina propria from DT and bDT treated mice was assessed by flow cytometry. Plots depict frequencies of NK cells (d), neutrophils (e), and eosinophils (f) among all live CD45^+^ cells. See Methods for cell gating strategies. Each point represents an individual mouse (n = 12 per group) and data shown are pooled from three independent experiments. ANOVA for frequency as a function of treatment, sex, and experimental replicate with *p*-value for the effect of treatment determined by Tukey’s ‘Honest Significant Difference’ method and then corrected for multiple comparisons (all parameters assessed by flow cytometry) using the FDR method of Benjamini and Hochberg. g-j. Protein was extracted from colonic tissue of DT and bDT treated mice and pro-inflammatory chemokines were quantified using a multiplexed bead assay. Plots depict abundance (normalized to weight of tissue) of CXCL10 (g), CXCL9 (h), CXCL1 (i), and CCL11 (j). Each point represents an individual mouse (n = 10 per group) and data shown represent tissue collected during three independent experiments. ANOVA for protein quantity as a function of treatment, sex, and experimental replicate with *p*-value for the effect of treatment determined by Tukey’s ‘Honest Significant Difference’ method and then corrected for multiple comparisons (all detectable chemokines measured by the multiplexed assay) using the FDR method of Benjamini and Hochberg. *P*-value < 0.05 *, < 0.01 **, < 0.0001 ****

One notable caveat to this conclusion was the possibility that the observed colonic inflammatory response was due a potentially non-specific effect of depleting a sizable population of colonic Treg cells, as IL-10 expressing Treg cells represented up to half of the overall colonic Treg cell population (Extended Data Fig. 8c). To test this potentiality, we generated *Il10^tdTomato-Cre^Foxp3^Thy1.1/GFP-DTR^* heterozygous female mice (hereafter referred to as *Foxp3^DTR-het^*) harboring roughly equal numbers of DT sensitive GFP-DTR- or DT resistant Thy1.1-expressing Treg cells due to the random X-inactivation. These mice allowed for depletion of approximately half of the overall pool of Treg cells regardless of their localization, activation, differentiation state, or IL-10 expression (Extended Data Fig. 9A & 9b). Contrary to the sequelae of depleting IL-10 expressing Treg cells, such “subset-unaware” halving of the colonic Treg cell population did not result in detectable weight loss or colon shortening, even though a mild increase in innate immune cell infiltration of the colon was observed (Extended Data Fig. 9c-g). This suggested that terminally differentiated IL-10 expressing Treg cells have a non-redundant function in supporting colon tissue and overall organismal health. To assess a role for these cells in settings of induced colonic injury and associated inflammation, mice were administered with DSS in drinking water for 2 days before and then throughout DT induced ablation of IL-10 expressing Treg cells (Extended Data Fig. 9h). This resulted in a rapid and severe weight loss in comparison to the control, non-depleted but DSS-treated group, extending the role of these cells in promoting tissue integrity at steady state into settings of inflammation (Extended Data Fig. 9i & 9j). Evaluation of the colonic tissues of these animals did not reveal any obvious worsening of histological signs of DSS induced pathology following IL-10 expressing Treg cell ablation except for the seemingly increased severity of Grade 3 inflammation on the simplified Geboes rubric, which indicates neutrophilic infiltration into the epithelial tissue^63^. This lack of a discernable histopathological difference may also be attributed to the severity of tissue damage in this model of colonic injury and inflammation at the late time-point analyzed.

### Terminal differentiation of IL-10 expressing Treg cells in the small intestine

The stability of IL-10 expression that distinguished terminally differentiated Treg cells in the colon was consistent with the well-documented abundance of IL-10 expressing Treg cells in that tissue ^32,47,48,64^ and raised a question as to whether the colon is unique in its ability to enable terminal differentiation of IL-10 expressing Treg cells. Given the phenotypic overlap between colonic and small intestinal (SI) Treg cells, as well as the integrated functions and lymphatic drainage of these tissues, we asked if this ability was also shared across these tissues. Thus, we assessed the stability of IL-10 expression by Treg cells in the SI lamina propria by analyzing these cells after their tamoxifen-induced “time-stamping” in *Il10^FM^* mice 21 days prior (Extended Data Fig. 10a & 10b). Similar to the colon, IL-10 expressing Treg cells were also highly abundant in the SI and virtually all were *Il10^stable^*. We further characterized these SI Treg cells by taking advantage of our *Il10^Foxp3-DTR^* mice, which allow identification of IL-10 expressing Treg cells in a manner compatible with intracellular and nuclear staining through the subset specific expression of the membrane bound GFP-DTR (Extended Data Fig. 10c-j). This analysis revealed substantial commonality between SI and LI IL-10 expressing Treg cells, including enriched expression of effector molecules such as CD39 and CTLA-4, and loss of CD62L expression (Extended Data Fig. 10e-g). At the same time, *Il10^stable^* Treg cells in the SI expressed either RORγt or GATA3, in contrast to their LI counterparts, which overwhelmingly were RORγt^+^ (Extended Data Fig. 10h & 10i). SI *Il10^stable^* Treg cells were also distinguished by their enriched expression of the activation/differentiation marker KLRG1 (Extended Data Fig. 10j). Collectively, the overall stability of IL-10 expressing SI Treg cells, paired with their heterogeneous TF and KLRG1 expression, raises the possibility that SI and LI tissues support terminal differentiation of distinct *Il10^stable^* Treg cell subsets whose shared and distinct mechanisms of differentiation and function in the SI versus colon remain to be elucidated.

## Discussion

Using genetic cell fate tracing and targeting we demonstrate that stable expression of the major immunomodulatory cytokine IL-10 defines a terminally differentiated effector Treg cell population in the colon with an essential non-redundant function in preventing spontaneous inflammation at this major barrier site. All IL-10 expressing Treg cells residing in the colon seemed to eventually adopt this fate following proliferative expansion, with the induction of IL-10 expression or accumulation of IL-10 expressing cells apparently dependent on the presence of antibiotic sensitive microbes. Although less well characterized, the *Il10^neg^* Treg cell population in the colon seemed to harbor distinct varieties of effector Treg cells. Importantly, our analysis of the consequences of antibiotic-induced depletion of the microbiota or bleomycin induced lung inflammation suggest that ongoing inflammatory exposure is not necessary to maintain nor sufficient to induce stability of IL-10 expression by activated Treg cells, respectively. The observation that alongside colon small intestine, but not other tissues afforded highly stable expression of IL-10 by Treg cells suggested distinct ability of the intestinal environment to support generation of *Il10^stable^* Treg cells.

Integrated analysis of transcriptomes and epigenomes of *Il10^neg^*, *Il10^recent^*, and *Il10^stable^* cells suggested attenuated TCR signaling alongside Maf and Rorγ(t) TF activities impart the distinct features and properties of terminally differentiated *Il10^stable^* Treg cells. The importance of Maf and Rorγ(t) for specifying the function of a population of colonic Treg cells has been demonstrated, supporting our results (Neumann et al., 2019; Ohnmacht et al., 2015; Sefik et al., 2015; Wheaton et al., 2017; Xu et al., 2018). The importance of TCR signaling in Treg cells for the induction of IL-10 expression is well established. However, our analysis of Treg cells post-activation strongly suggest that ongoing TCR signaling becomes an impediment to continued IL-10 expression and more broadly, terminal differentiation. This hypothesis is validated by experimental manipulation of TCR expression after activation and acquisition of IL-10 expression. That the identified population of IL-10 expressing Treg cells becomes independent of TCR signaling upon their terminal differentiation suggests acquisition of “innate-like” function in these cells. This “innate-like” functional state is reminiscent of unconventional and some conventional T cell populations, which have been shown to acquire TCR independent functionality when highly differentiated ^55,65–69^. This finding suggests that loss of TCR dependence for functional outputs represents a common feature of terminally differentiated anti-inflammatory, tissue supportive, and effector T cell populations. Furthermore, and in contrast to studies of CD8^+^ T cells, which suggest that salutary or maladaptive pathological immune responses are reliant on ‘stem-like’ populations, while their terminally differentiated progeny are deemed dysfunctional, our experiments suggest that terminally differentiated effector Treg cells play an essential functional role in inflammatory processes by maintaining immune balance ^56–59^.

While our experiments do not unambiguously prove that the observed differences in the properties of *Il10^neg^* and *Il10^stable^* cells indicate whether ongoing TCR stimulation is required to specify the *Il10^neg^* state or that attenuation of TCR signaling is required to calcify the *Il10^stable^* state, the analysis of cells from *Il10^iΔTCR^* mice favors the latter possibility. It is possible that TCR signaling is ongoing in both populations, but is qualitatively distinct, with diminished output from specific downstream signaling cascades in *Il10^stable^* cells. In this regard, Nr4a2 is a likely candidate TF favoring the *Il10^neg^* over the *Il10^stable^* Treg cell gene expression program, given that *Nr4a2* expression and apparent TF activity is specifically lost in *Il10^stable^* Treg cells. Likewise, *Tespa1* and *Themis* genes, encoding positive regulators of TCR signaling studied mostly in the context of thymic development, were differentially expressed between the two populations and are additional candidates affecting the two differentiation paths of effector Treg cells by modulating TCR signaling.

Interestingly, members of the Tec family of kinases, which are known to participate in and enhance proximal TCR signaling, are also differentially expressed across the colonic Treg cell populations analyzed herein ^70,71^. However, in contrast to other positive regulators of TCR signaling, these genes have high expression in *Il10^stable^* cells. Itk is known to play distinct roles in Treg cells: playing a negative role in thymic Treg cell differentiation, yet supporting peripherally generated Treg cell function ^72–74^. Whether the other Tec kinases have similar functions is unknown, but is not unlikely. How the continued expression of these kinases reconciles with the overall reduction in TCR signaling in *Il10^stable^* cells, as well as the contributions of Itk, Tec, and Txk to the phenotype and function, if any, of these cells remain to be explored.

Previous investigation of TFs involved in the specification of tissue Treg cells by the Feurer group identified Batf/BATF as an important player in facilitating their tissue supportive functions in both mouse and human ^25,26^. Interestingly, the Batf motif appears in our analysis as specifically associated with increased accessibility of gene loci with increased expression in *Il10^neg^* cells. This observation suggests that the *Il10^neg^* population in the colon shares features with Treg cells performing tissue supportive functions in other non-lymphoid organs. However, the *Il10^stable^* Treg cells we have characterized have high expression of a specific complement of genes encoding molecules known to support tissue function. While we have yet to test a role for Batf in *Il10^stable^* Treg cell function, our results suggest that tissue supportive functions are not exclusive to a specific population of Treg cells and are not exclusively controlled by Batf.

Signaling through ICOS, a co-stimulatory molecule for T cells, has been previously shown to be important for the stability and function of colonic Treg cells, but importantly, dispensable for IL-10 expression by T cells in the colon ^75^. In contrast to other genes associated with the phenotype and function of *Il10^stable^* Treg cells, ICOS was notably dependent on continued TCR signaling (data not shown). Therefore, at which stage ICOS signaling is important for colonic Treg cell development, and whether its continued expression in *Il10^stable^* cells is functionally relevant or simply represents a by-product of residual TCR stimulation remain to be established.

Our study identifies transcriptional programs and highlights upstream signaling pathways contributing to the specification of distinct populations of colonic effector Treg cells. In addition, we demonstrate that a subset of colonic Treg cells undergo functional specialization, assuming a stable terminally differentiated state, which is robust to environmental perturbations, and is not reliant on ongoing conditioning for its maintenance or elaboration of effector functions. That these terminally differentiated Treg cells exert an essential function in preventing severe local inflammation contrasts with the prevalent view of terminal differentiation of conventional T cells, which is thought to lead to a dysfunctional or exhausted state. The precise mechanisms by which these cells support tissue and organismal health remain unclear. A relatively moderate innate effector cell accumulation in the colon observed upon depletion of IL-10 expressing Treg cells raise the possibility that this subset acts on parenchymal cells or stromal cells. Importantly, given the dispensable role of ongoing production of IL-10 by Treg cells in controlling this inflammation in adulthood, their essential function must be reliant on the elaboration of a range of tissue supportive and suppressive molecules, likely including those which enriched expression in this subset, such as CD39, CTLA-4, galectin-3, and granzyme B. Furthermore, our studies offer a proof-of-principle for a genetic approach for studying the context-specific function and differentiation of pro- and anti-inflammatory T cells. Thus, our studies reveal that colonic IL-10 expressing Treg cells have an essential role in maintaining colonic health and do so by deploying a combination of immunoregulatory and other effector mechanisms beyond, or in addition to, IL-10. Our findings and approaches help advance the mechanistic understanding of chronic inflammatory conditions affecting the intestine and may assist the development of specific rational therapies for their treatment, which would benefit from a fuller understanding of the cell types and effector molecules dysregulated therein.

## Author Contributions

S.D. and A.Y.R. conceived the project and wrote the manuscript. S.D. designed, performed, and analyzed experiments, with help from G.B., Z.-M.W., and P.G. Y.P., C.K., and C.S.L. contributed to the development of the RNA-seq and ATAC-seq analysis strategy. A.G.L. generated the *Il10^tdTomato-Cre^* and *Il10^tdTomato-CreER^* mice. A.G. generated the *Foxp3^LSL-DTR^* mice. A.Y.R. contributed to data analysis.

## Acknowledgments

We thank M. Faire, J. Verter, E. Andretta, and A. Bravo for help with animal husbandry. We thank R. Bou Puerto for invaluable assistance. We thank the Memorial Sloan Kettering Integrated Genomics Core facility for performing all sequencing and the Single Cell Research Initiative for processing the scRNA-seq samples. We thank all the members of the Rudensky laboratory for input and A. Mendoza for critical reading of the manuscript. This work was supported by the NCI Cancer Center Support Grant P30 CA008748, NIH grant R37 AI034206 (A.Y.R.), and the Ludwig Center at the Memorial Sloan Kettering Cancer Center (A.Y.R.). A.Y.R. is an investigator with the Howard Hughes Medical Institute. S.D. was supported by the General Atlantic Fellowship. The content is solely the responsibility of the authors and does not necessarily represent the official views of the NIH.

## Declaration of Interests

A.Y.R. is an SAB member and has equity in Sonoma Biotherapeutics, RAPT Therapeutics, Surface Oncology, and Vedanta Biosciences, and is an SAB member of BioInvent and a co-inventor or has IP licensed to Takeda that is unrelated to the content of the present study. The remaining authors declare no competing interests.

## Extended Data Figure Legends

**Extended Data Fig. 1.**
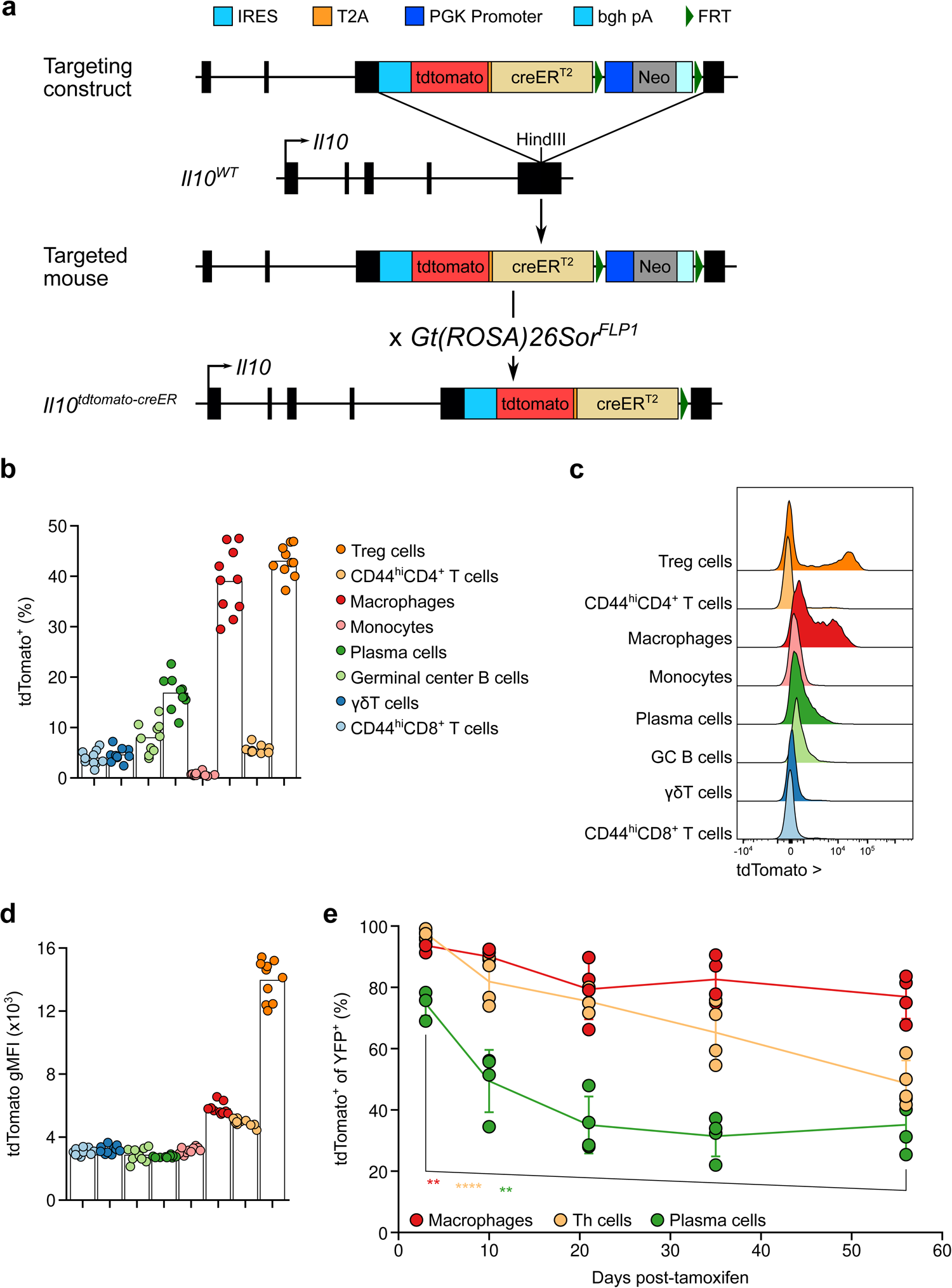
Characterization of *Il10^FM^* mice.

a. Strategy for generating *Il10^tdTomato-CreER^* mice. Insertion of an IRES-tdTomato-T2A-CreER cassette into the 3’ untranslated region of the *Il10* gene allows for co-expression of a fluorescent reporter and inducible recombinase with endogenous *Il10* expression. Elements are drawn to scale with the exception of FRT sites. See Methods for details.

b-d. Cells were isolated from the colonic lamina propria of 15-week-old *Il10^FM^* mice. Frequencies (b) and representative histograms (c) of tdTomato^+^ cells among different immune cells subsets, and their level of tdTomato expression (geometric mean fluorescent intensity, gMFI) (d). See Methods for cell gating strategies. (b, d) Each point represents an individual mouse (n = 10) and data are representative of two independent experiments. (c) Histograms are from a single mouse, representative of 20 mice from two independent experiments.

e. *Il10^FM^* mice were treated at 7 weeks of age with tamoxifen and analyzed 3 to 56 days later. Mice analyzed together were littermates and treated at different times with tamoxifen in order be analyzed on the same day. Frequencies of tdTomato^+^ cells among YFP^+^ macrophages, plasma cells, and Th cells isolated from the colonic lamina propria of mice at indicated days after tamoxifen treatment. Each point represents an individual mouse, lines indicate mean per tissue across time points, bars indicate standard deviation, and data are pooled from two independent experiments (n = 4 mice per time point, with different mice for each time point). ANOVA with Holm’s correction for multiple comparisons.

*P*-value < 0.01 **, < 0.0001 ****

**Extended Data Fig. 2.**
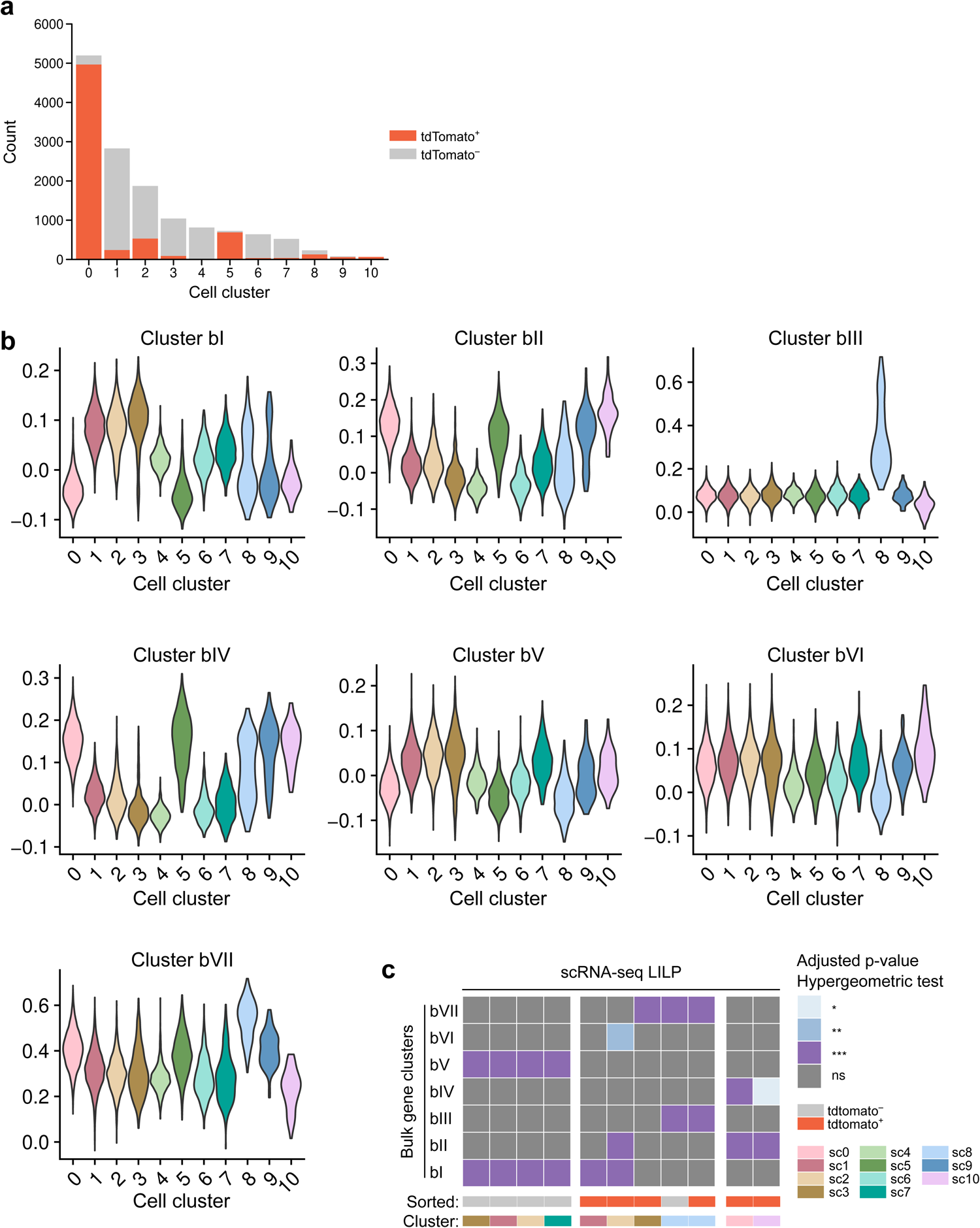
Expression of bulk RNA-seq gene clusters by scRNA-seq cell populations. Data are from scRNA-seq analysis presented in Fig. 3. a. Counts of cells originating from the tdTomato^+^ (red, IL-10 expressing) or tdTomato^−^ (grey, IL-10 non-expressing) samples assigned to each nearest neighbor cluster. b. The scRNA-seq cells were scored for expression of bulk RNA-seq gene clusters and violin plots detect per gene cluster score by each scRNA-seq cell cluster. See Methods for details. c. scRNA-seq cells were divided into subsets as in Fig. 3c, and statistical significance of enrichment of genes belonging to bulk RNA-seq k-means clusters bI-VII among genes with significantly higher expression in each scRNA-seq subset versus all other cells was determined. Shading indicates binned p-values of one-tailed (upper tail) hypergeometric test for enrichment, with *p*-values adjusted for multiple comparisons using Holm’s method. See Methods for details. *P*-value > 0.05 ns, < 0.05 *, < 0.01 **, <0.001 ***

**Extended Data Fig. 3.**
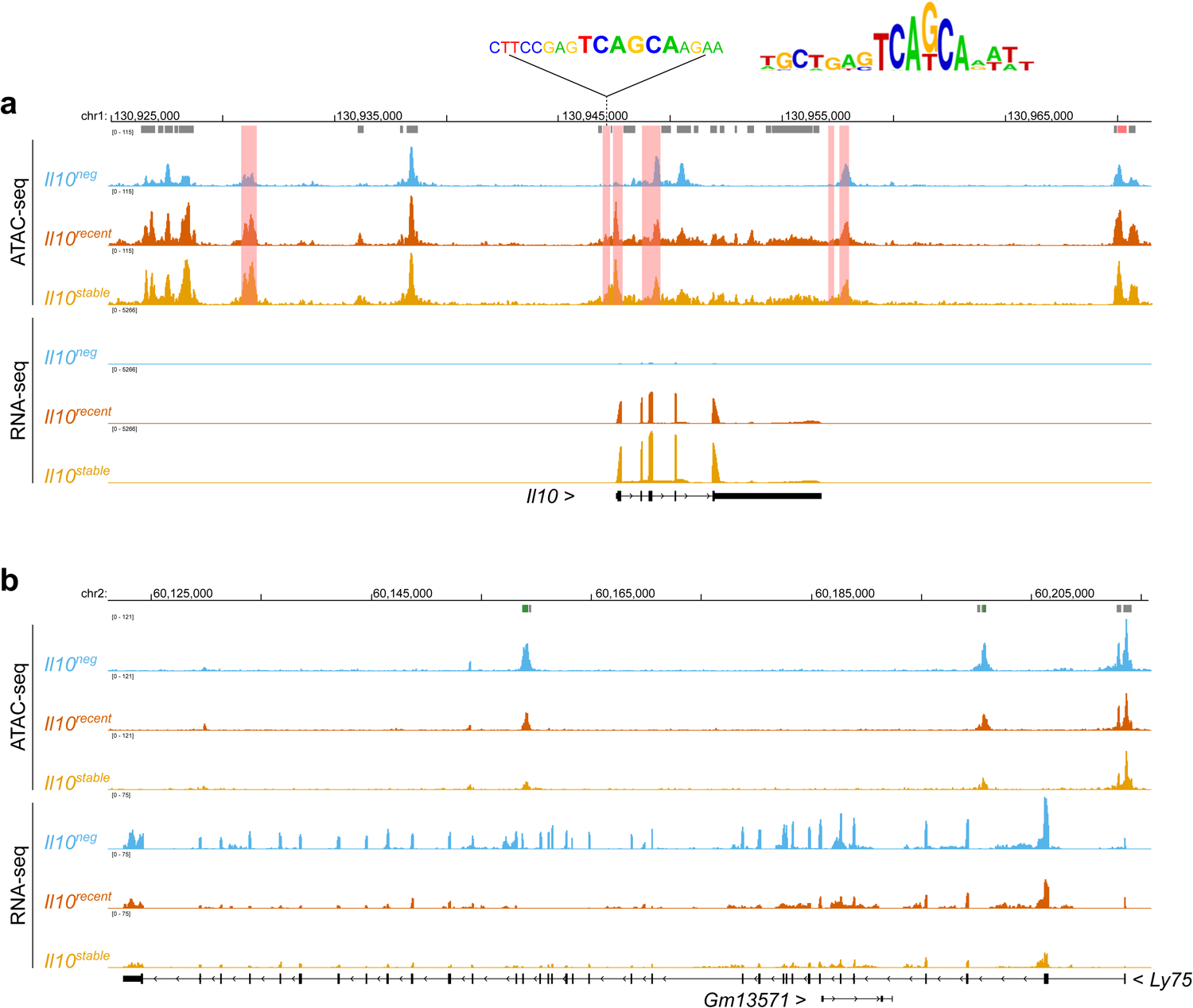
Chromatin accessibility and gene expression changes associated with Maf and Nr4a2/3 motifs. *Il10^neg^*, *Il10^recent^*, and *Il10^stable^* Treg cells, as defined in Fig. 1b, were sorted from the colons of 10-week-old *Il10^FM^* mice treated 21 days prior with tamoxifen and subjected to ATAC-seq and bulk RNA-seq analysis. See Methods for details. a, b. Representative tracks showing normalized pile-up of ATAC-seq and RNA-seq reads aligning to the *Il10* (a) and *Ly75* (b) gene loci. Coloring indicates cell states. Coordinates in a modified mm39 genome assembly are indicated. Grey bars indicate ATAC-seq peaks, orange bars indicate peaks containing at least one Maf motif, and green bars indicate peaks containing at least one Nr4a2/3 motif. Upper inset shows canonical Maf motif alongside the highest confidence identified Maf motif in the promoter region of *Il10*.

**Extended Data Fig. 4.**
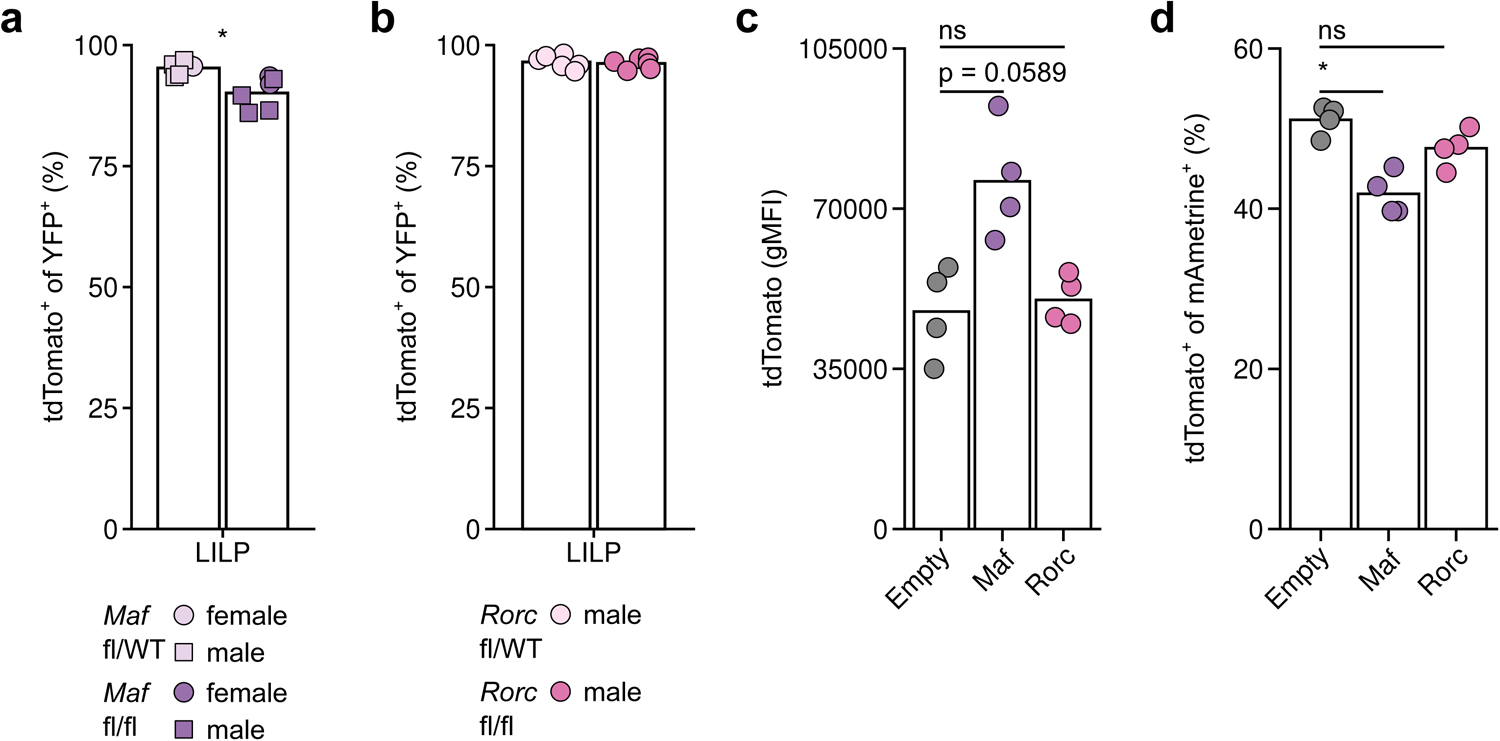
Modulating Maf and Rorγt expression in Treg cells minimally perturbs IL-10 expression. a-b. 8- to 12-week-old male and female *Il10^tdTomato-CreER^Maf^FL/WT^* or *Maf^FL/FL^* (a) or *Il10^tdTomato-CreER^Rorc^FL/WT^* or *Rorc^FL/FL^* mice were treated with tamoxifen on days 0, 4, 11, 18, 25, and 32 and cells were isolated from the LILP on day 35 for flow cytometric analysis. Plots show frequencies of tdTomato^+^ cells among YFP^+^ Treg cells (Thy1.1^+^CD4^+^TCRβ^+^). Each point represents an individual mouse (n = 6 per group) and data are pooled from two (Rorc) or three (Maf) independent experiments. c-d. IL-10 non-expressing Treg cells (Thy1.1^+^CD4^+^TCRβ^+^tdTomato^−^) were sorted from pooled secondary lymphoid organs and activated in vitro using anti-CD3 and anti-CD28. Cells were transduced with Cre-activated retroviral vectors expressing Maf or Rorγt cDNA subsequent to Cre expression, with mAmetrine fluorescence reporting on cDNA expression through an IRES. Plots depict geometric mean of fluorescence intensity for tdTomato (c) or frequencies of tdTomato^+^ cells among Treg cells (d) three days after transduction. Each point represents cells obtained from an individual mouse (n = 4) and data are from a single experiment. *P*-value > 0.05 ns, < 0.05 *

**Extended Data Fig. 5.**
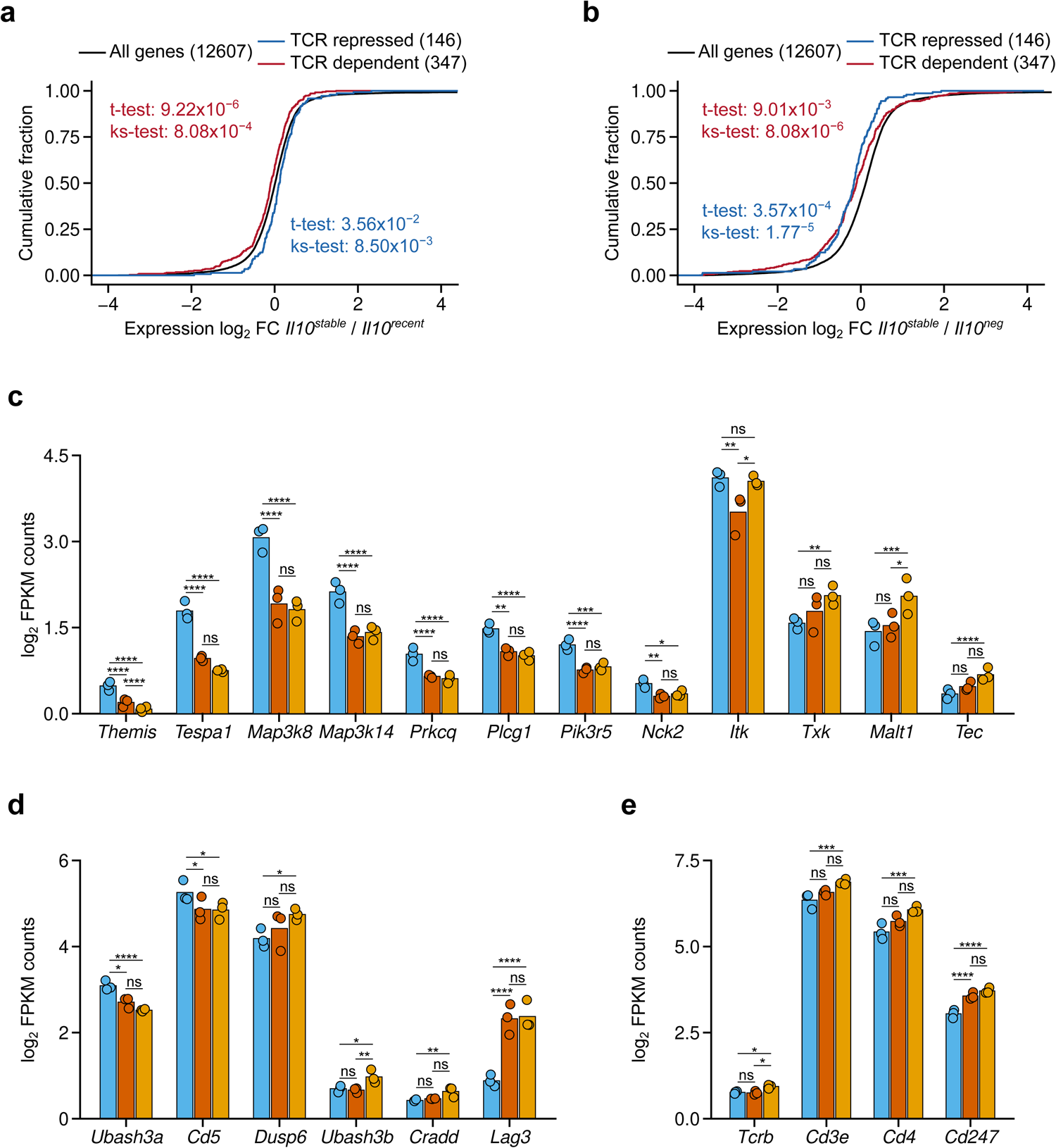
TCR dependent gene expression is attenuated in *Il10^stable^* colonic Treg cells. RNA was isolated from *Il10^neg^*, *Il10^recent^*, and *Il10^stable^* Treg cells, as defined in Fig. 1b from the colons of 10-week-old *Il10^FM^* mice treated 21 days prior with tamoxifen and sequenced. a, b. Cumulative distribution function plot of mean log_2_ transformed gene expression fold change for *Il10^stable^* versus *Il10^recent^* (a) and *Il10^stable^* versus *Il10^neg^* (b) samples. All genes (black) and genes with increased (red) or decreased (blue) expression in TCR sufficient versus deficient Treg cells. *P*-values calculated by two-sided *t*-test and two-sided Kolmogorov-Smirnov test for genes with increased (red) or decreased (blue) expression versus all genes are indicated. Data are from RNA-seq analysis presented in Fig. 2 and 3. c-e. Plots showing log_2_ transformed FPKM counts of indicated genes. Coloring indicates cell population. Negative binomial fitting with Wald’s significance test and the Benjamini & Hochberg correction for multiple comparisons. Significance testing and correction was performed on all genes. Data are from RNA-seq analysis presented in Fig. 2 and 3. *P*-value > 0.05 ns, < 0.05 *, < 0.01 **, <0.001 ***, < 0.0001 ****

**Extended Data Fig. 6.**
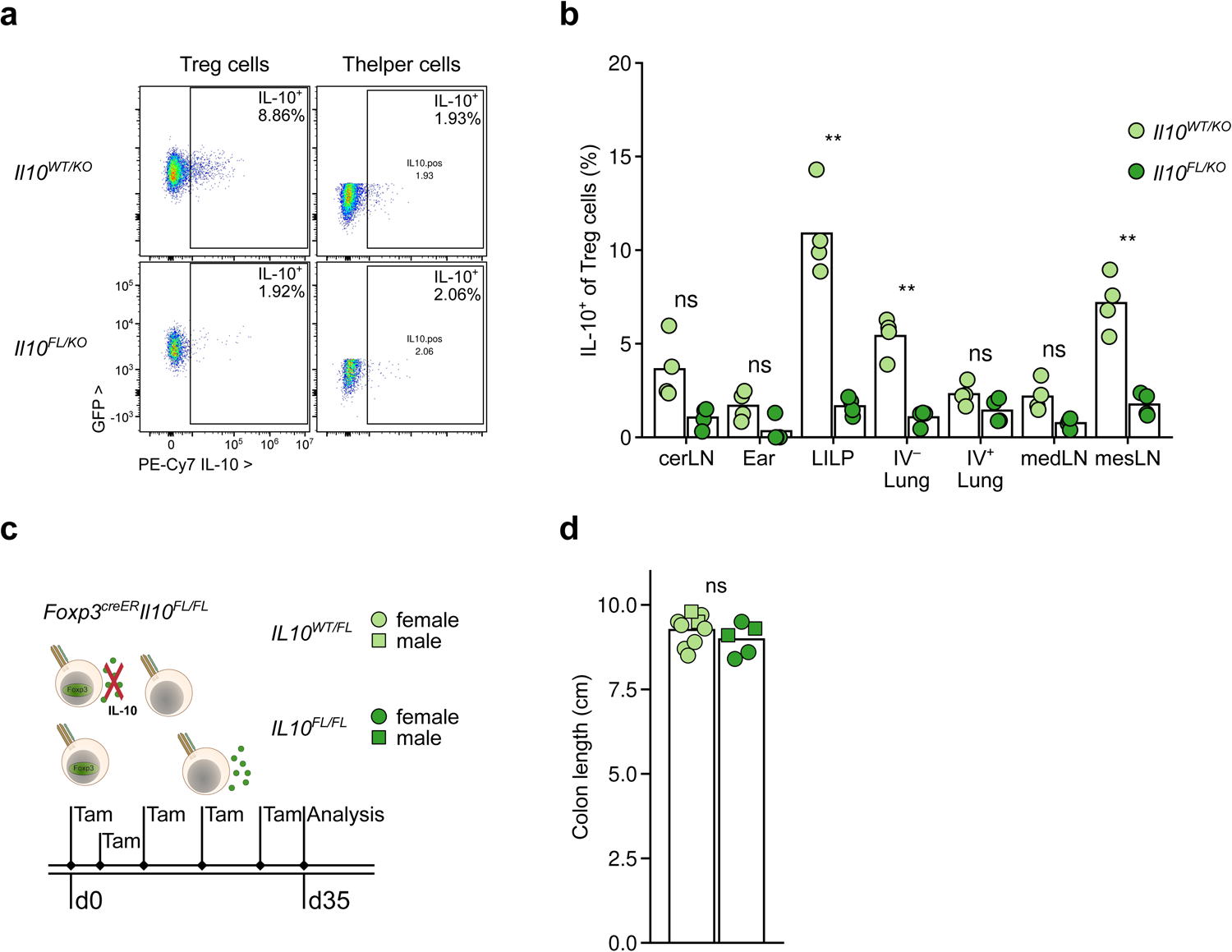
Analysis of function of Treg cell derived IL-10 in the colon. a, b. 8- to 10-week-old *Foxp3^CreER^Il10^FL/KO^* and littermate control *Foxp3^CreER^Il10^WT/KO^* mice were treated with tamoxifen on days 0 and 2 and cells were isolated from the indicated tissues on day 16. Cells were re-stimulated *ex vivo* to induce cytokine production, and intracellular cytokine accumulation was assessed by flow cytometry. Representative 2D flow plots showing IL-10 production by Treg cells (left; GFP^+^CD4^+^TCRβ^+^) and conventional activated helper CD4 T cells (Thelper) (right; GFP^−^ CD44^+^CD4^+^TCRβ^+^) in the colonic lamina propria from *Foxp3^CreER^Il10^WT/KO^* (top) and *Foxp3^CreER^Il10^FL/KO^* (bottom) mice (a) and plot showing frequencies of IL-10^+^ Treg cells in various tissues from *Foxp3^CreER^Il10^WT/KO^* (light green) and *Foxp3^CreER^Il10^FL/KO^* (dark green) mice (b). (a) Treg and conventional CD4 T cells (Tconv) gated cells from a single *Il10^WT/KO^* and *Il10^FL/KO^* mouse, representative of 9 mice of each genotype from three experiments. (b) Each point represents an individual mouse (n = 4 per genotype) and data are representative of three independent experiments. c-d. 8- to 10-week-old *Foxp3^CreER^Il10^FL/FL^* and littermate control *Foxp3^CreER^Il10^WT/FL^* mice were treated with tamoxifen on days 0, 4, 11, 18, 25, and 32 before analysis by flow cytometry on day 35. Legend for plots throughout figure with different treatment groups (IL-10 deleted, *Foxp3^CreER^Il10^FL/FL^* versus IL-10 retaining, *Foxp3^CreER^Il10^WT/FL^*) colored as dark green or light green, respectively, and female or male mice depicted as circles or squares, respectively; data representing pooled male/female samples are depicted as triangles (c, top right). Experimental treatment timeline (c, bottom). Plot showing colon lengths on day 35 of IL-10 deleted and IL-10 retaining mice (d). Each point represents an individual mouse (n = 5-9) and data shown are pooled from two independent experiments. ANOVA for colon length as a function of genotype, sex and replicate, with *p*-value for the effect of genotype determined by Tukey’s ‘Honest Significant Difference’ method. *P*-value > 0.05 ns, < 0.01 **, <0.001 ***

**Extended Data Fig. 7.**
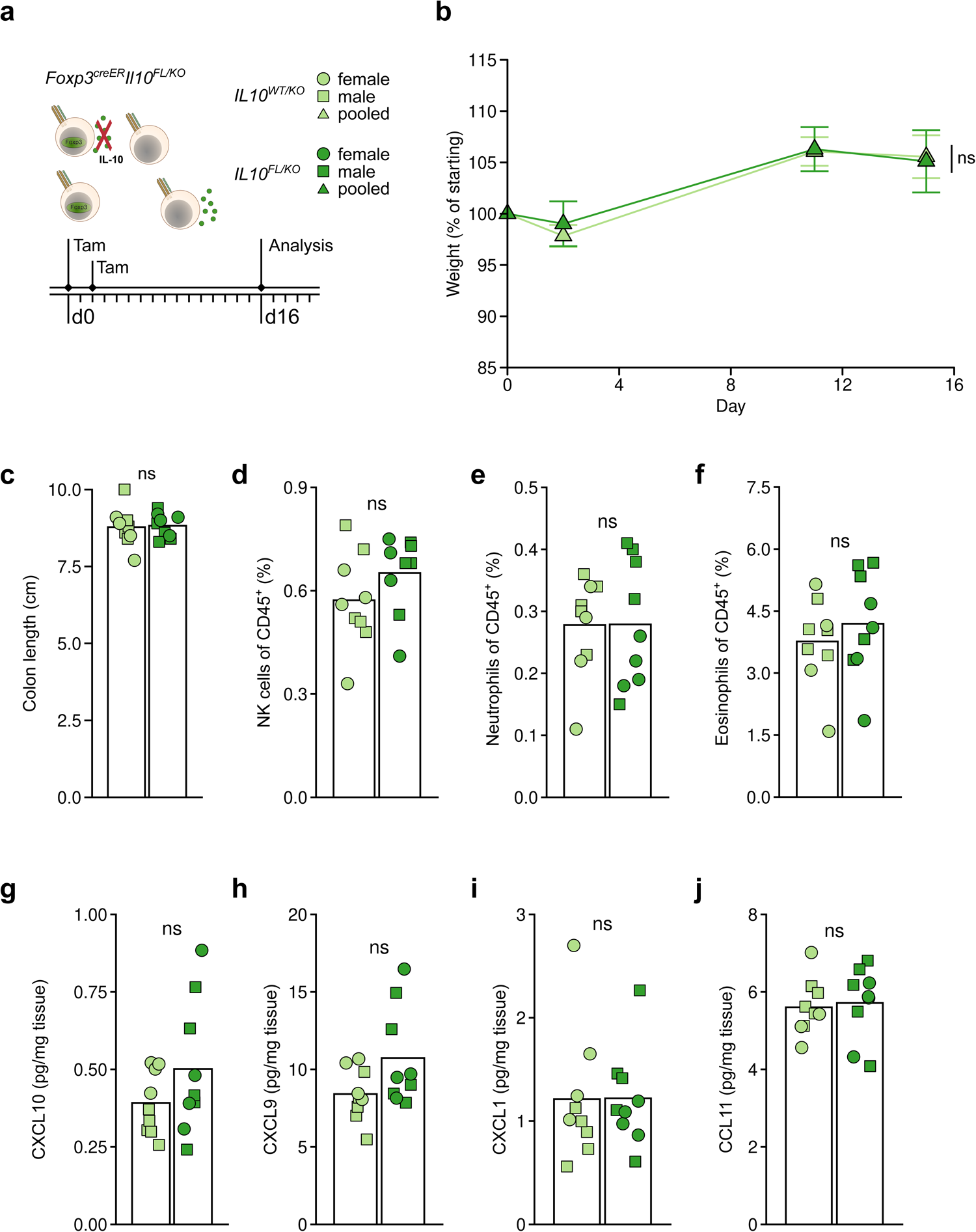
IL-10 expression by Treg cells in adult mice is dispensable for control of colonic inflammation at steady state. 8- to 10-week-old *Foxp3^CreER^Il10^FL/KO^* and littermate control *Foxp3^CreER^Il10^WT/KO^* mice were treated with tamoxifen on days 0 and 2 before analysis by flow cytometry on day 16. a. Legend for plots throughout figure with different treatment groups (IL-10 deleted, *Foxp3^CreER^Il10^FL/KO^* versus IL-10 retaining, *Foxp3^CreER^Il10^WT/KO^*) colored as dark green or light green, respectively, and female or male mice depicted as circles or squares, respectively; data representing pooled male/female samples are depicted as triangles (top right). Experimental treatment timeline (bottom). b. Plot showing body weights of mice harboring IL-10 deleted or IL-10 retaining Treg cells over time. Weights are normalized to starting body weight for each mouse. Plots show mean ± SD and represent data pooled from two independent experiments (n = 3-4 mice per experiment per group, n = 7 per group overall, with each mouse repeatedly weighed for each time point). ANOVA for body weight change as a function of genotype, sex, and time with *p*-value for the effect of genotype determined by Tukey’s ‘Honest Significant Difference’ method. c. Plot showing colon lengths on day 16 of IL-10 deleted and IL-10 retaining mice. Each point represents an individual mouse (n = 9) and data shown are pooled from three independent experiments. ANOVA for colon length as a function of genotype and sex, with *p*-value for the effect of genotype determined by Tukey’s ‘Honest Significant Difference’ method. d-f. Cellular composition of the colonic lamina propria from IL-10 deleted and IL-10 retaining mice was assessed by flow cytometry. Plots depict frequencies of NK cells (d), neutrophils (e), and eosinophils (f) among total live CD45^+^ cells. See Methods for cell gating strategies. Each point represents an individual mouse (n = 9) and data are pooled from three independent experiments. ANOVA for frequency as a function of genotype and sex with *p*-value for the effect of genotype determined by Tukey’s ‘Honest Significant Difference’ method and then corrected for multiple comparisons (all parameters assessed by flow cytometry) using the FDR method of Benjamini and Hochberg. g-j. Protein was extracted from colonic tissue of mice harboring IL-10 deleted or IL-10 retaining Treg cells and pro-inflammatory chemokines were quantified using a multiplexed bead assay. Plots depict abundance (normalized to weight of tissue) of CXCL10 (g), CXCL9 (h), CXCL1 (i), and CCL11 (j). Each point represents an individual mouse (n = 9 per group) and data shown represent tissue collected during three independent experiments. ANOVA for protein quantity as a function of genotype and sex with *p*-value for the effect of genotype determined by Tukey’s ‘Honest Significant Difference’ method and then corrected for multiple comparisons (all detectable chemokines measured by the multiplexed assay) using the FDR method of Benjamini and Hochberg. *P*-value > 0.05 ns

**Extended Data Fig. 8.**
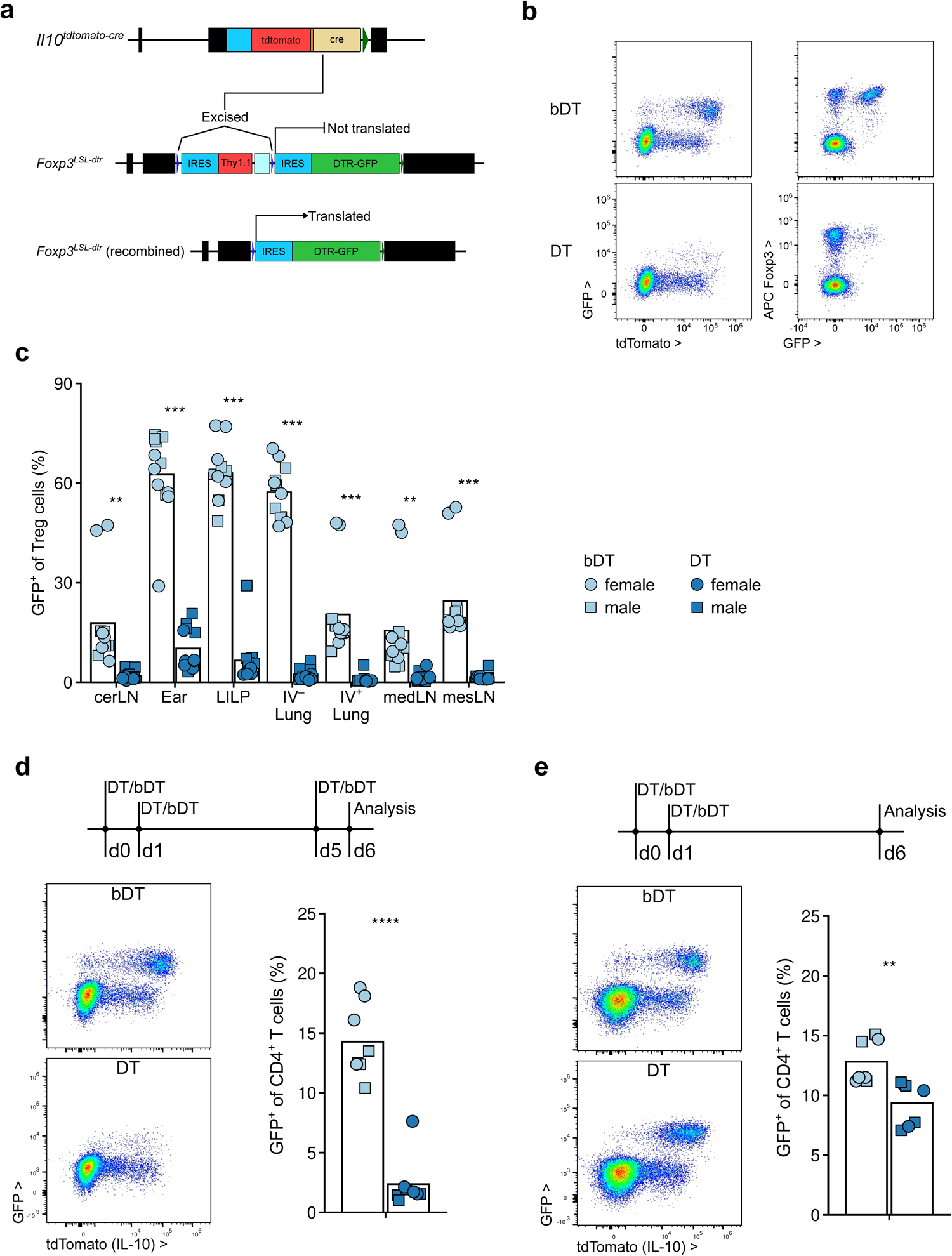
Analysis of IL-10 expressing Treg cell depletion model. a. Schematic of targeted *Il10* and *Foxp3* loci in *Il10^tdTomato-Cre^* and *Foxp3^LSL-DTR^* mice. Insertion of an IRES-tdTomato-T2A-nls-Cre cassette into the 3’ untranslated region of the *Il10* gene allows for co-expression of a fluorescent reporter and constitutively active Cre recombinase with preserved endogenous *Il10* expression. Insertion of an loxP-IRES-Thy1.1-BGHpA-loxP-IRES-GFP-DTR cassette into the 3’ untranslated region of the *Foxp3* gene allows for Cre-dependent diphtheria toxin receptor (DTR) and GFP reporter expression with preserved endogenous *Foxp3* expression. Bottom panel shows recombined (LSL cassette deleted due to Cre recombinase activity) allele permitting DTR expression. See Methods for details. b, c. 8- to 11-week-old *Il10^tdTomato-Cre^Foxp3^LSL-DTR^* (*Il10^Foxp3-DTR^*) mice were treated with active (DT) or heat-inactivated (boiled, bDT) diphtheria toxin over 16 days before analysis by flow cytometry. Representative 2D flow plots showing IL-10 (tdTomato) expression by GFP^+^ and GFP^−^ CD4^+^ T cells (left) and GFP expression versus Foxp3 protein staining in CD4^+^ T cells (right) from the colonic lamina propria from bDT (top) and DT (bottom) treated *Il10^Foxp3-DTR^* mice (b) and plots showing frequency of GFP^+^ cells among Treg cells in various tissues from bDT (light blue) and DT (dark blue) treated *Il10^Foxp3-DTR^* mice (c). (b) CD4^+^ T cells from a single DT and bDT treated mouse, representative of 12 mice for each group from three independent experiments. (c) Each point represents an individual mouse (n = 12 per group) and data shown are pooled from three independent experiments. d-e. *Il10^Foxp3-DTR^* mice were treated with DT or heat-inactivated bDT diphtheria toxin on days 0, 1, and 5 (d) or days 0 and 1 before analysis by flow cytometry on day 6 (e). Represent 2D flow plots (left) and summary plots (right) showing frequency of GFP^+^ cells among CD4^+^ T cells in LILP from bDT (light blue) and DT (dark blue) treated *Il10^Foxp3-DTR^* mice. Each point represents an individual mouse (n = 8 per group) and data shown are pooled from two independent experiments for each treatment regimen. *P*-value < 0.01 **, <0.001 ***, <0.0001 ****

**Extended Data Fig. 9.**
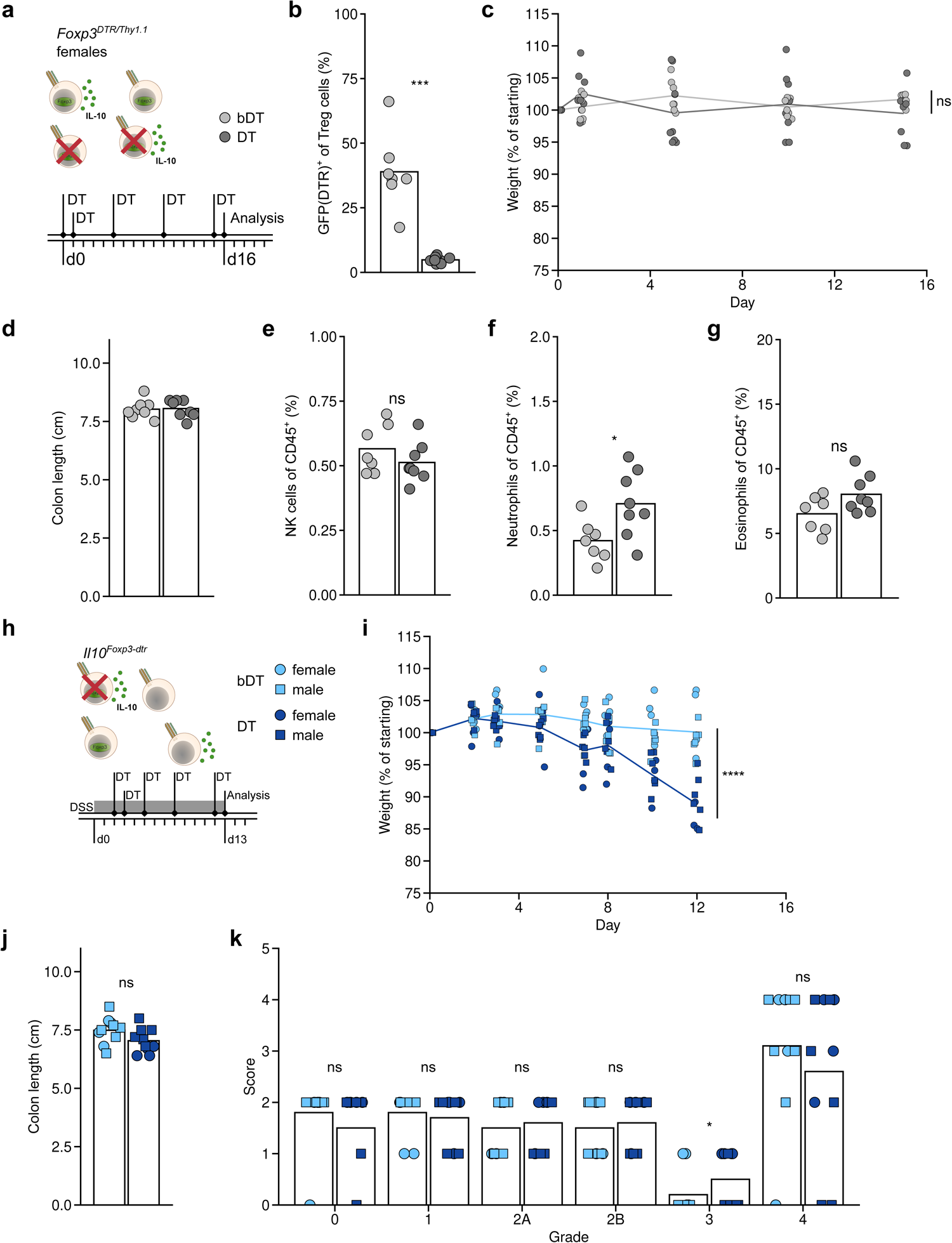
Further characterization of colonic IL-10 expressing Treg cell function at homeostasis and during colitis. a-g. 8- to 11-week-old *Il10^tdTomato-Cre^Foxp3^GFP-DTR/Thy1.^*^1^ (*Foxp3^DTR-het^*) mice were treated with active (DT) or heat-inactivated (boiled, bDT) diphtheria toxin over 16 days before analysis by flow cytometry. a. Diagram depicting DT-induced ablation of half of all Treg cells irrespective of IL-10 expression in *Foxp3^DTR-het^* mice (top left). Legend for plots throughout the figure with DT and bDT groups colored as light grey or dark grey, respectively. Experimental treatment regimen timeline (bottom). b. Plot showing frequency of GFP-DTR^+^ Treg cells among all Treg cells in the LILP of DT and bDT treated *Foxp3^DTR-het^* mice. c. Plot showing body weights of DT and bDT treated groups over the time course of the experiment. Weights are normalized to starting body weight for each mouse. Plots show mean ± SD and represent data pooled from two independent experiments (n = 3-4 mice per experiment per group, with each mouse weighed repeatedly for each time point). ANOVA for weight change as a function of treatment, experimental replicate, and time with *p*-value for the effect of treatment determined by Tukey’s ‘Honest Significant Difference’ method. d. Plot showing colon lengths on day 16 of DT and bDT treated mice. Each point represents an individual mouse (n = 8 for each group) and data shown are pooled from two independent experiments. ANOVA for colon length as a function of treatment and experimental replicate with *p*-value for the effect of treatment determined by Tukey’s ‘Honest Significant Difference’ method. e-g. Cellular composition of the colonic lamina propria from DT and bDT treated mice was assessed by flow cytometry. Plots depict frequencies of NK cells (e), neutrophils (f), and eosinophils (g) among all live CD45^+^ cells. See Methods for cell gating strategies. Each point represents an individual mouse (n = 8 per group) and data shown are pooled from two independent experiments. ANOVA for frequency as a function of treatment and experimental replicate with *p*-value for the effect of treatment determined by Tukey’s ‘Honest Significant Difference’ method and then corrected for multiple comparisons (all parameters assessed by flow cytometry) using the FDR method of Benjamini and Hochberg. h-k. 8- to 10-week-old *Il10^tdTomato-Cre^Foxp3^LSL-DTR^* (*Il10^Foxp3-DTR^*) mice were given DSS-laden (1.5% w/v) drinking water and then treated with active (DT) or heat-inactivated (boiled, bDT) diphtheria toxin over 11 days before analysis by flow cytometry on day 13 after DSS initiation. h. Diagram depicting targeted and punctual DT-induced ablation of IL-10 expressing Treg cells in *Il10^Foxp3-DTR^* mice (top left). Legend for plots throughout the figure with DT and bDT groups colored as sky blue or navy blue, respectively, and female or male mice depicted as circles or squares, respectively. Experimental treatment regimen timeline (bottom). i. Plot showing body weights of DT and bDT treated groups over the time course of the experiment. Weights are normalized to starting body weight for each mouse. Plots show mean ± SD and represent data pooled from four independent experiments (n = 2-4 mice per experiment per group, with n = 10 for each group overall and with each mouse weighed repeatedly for each time point). ANOVA for weight change as a function of treatment, sex, experimental replicate, and time with *p*-value for the effect of treatment determined by Tukey’s ‘Honest Significant Difference’ method. j. Plot showing colon lengths on day 13 of DT and bDT treated mice. Each point represents an individual mouse (n = 10 for each group) and data shown are pooled from four independent experiments. ANOVA for colon length as a function of treatment, sex, and experimental replicate with *p*-value for the effect of treatment determined by Tukey’s ‘Honest Significant Difference’ method. k. Histopathological scoring of colon sections collected on day 13 from DT and bDT treated mice following the Simplified Geboes Score rubric. See Methods for details. Each point represents an individual mouse (n = 10 for each group) and data shown are pooled from four independent experiments. ANOVA for score as a function of treatment, sex, grade, and experimental replicate with *p*-value for the effect of treatment determined by Tukey’s ‘Honest Significant Difference’ method. *P*-value > 0.05 ns, < 0.05 *, <0.001 ***

**Extended Data Fig. 10.**
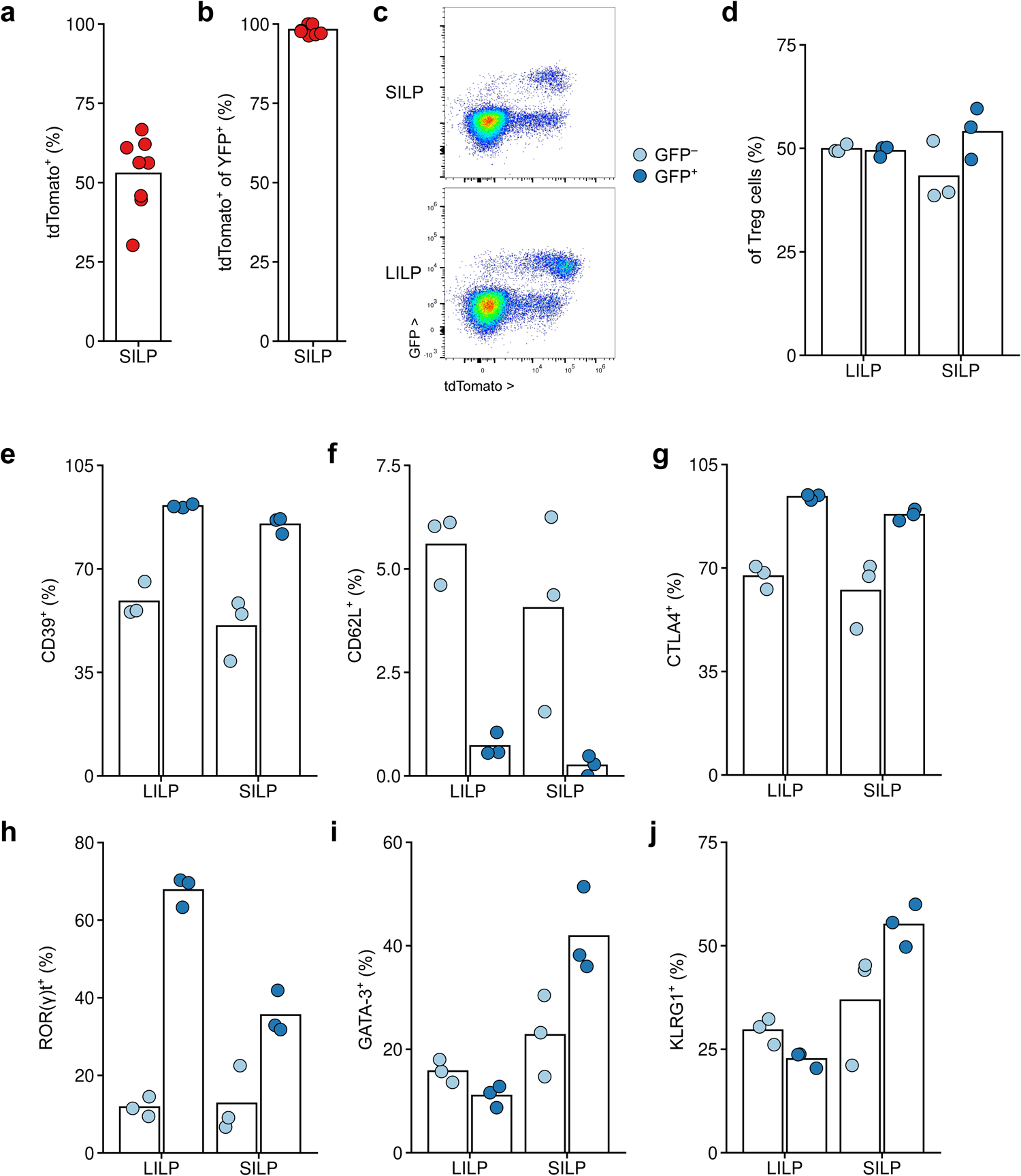
Characterization of small intestinal IL-10 expressing Treg cells. a-b. *Il10^FM^* mice were treated at 8 weeks of age with tamoxifen and analyzed 21 days later. Frequencies of tdTomato^+^ cells among Treg cells (a) and among YFP^+^ Treg cells (b) isolated from SILP. Each point represents an individual mouse and data are pooled from two independent experiments (n = 4 mice per replicate). c-j. Cells were isolated from the SILP and LILP of *Il10^Foxp3-DTR^* mice and analyzed by flow cytometry. c. Representative 2D flow cytometry plot showing GFP and tdTomato expression among CD4^+^ T cells from the SILP (top) and LILP (bottom). d. Plots showing frequencies of GFP^−^ (light blue) and GFP^+^ (dark blue) Treg cells from the LILP and SILP. e-j. Plots showing frequencies of CD39 (e), CD62L (f), CTLA4 (g), Rorγt (h), Gata3 (i), and KLRG1 (j) expression among GFP^−^ (light blue) and GFP^+^ (dark blue) Treg cells from the LILP and SILP of *Il10^Foxp3-DTR^* mice. d-j. Each point represents an individual mouse (n=3), with paired GFP^−^ and GFP^+^ cells from each mouse. Data are from a single experiment.

## Data availability

Bulk RNA-seq, ATAC-seq and single cell RNA-seq data will be available from the GEO database (accession number pending).

## Code availability

All custom code used in analysis of data and generation of plots is available upon request.

## Material availability

Mouse strains generated for this study are available upon request from the corresponding author with a completion of an MTA with MSKCC.

## Supplementary Figures

**Supplementary Fig. 1.**
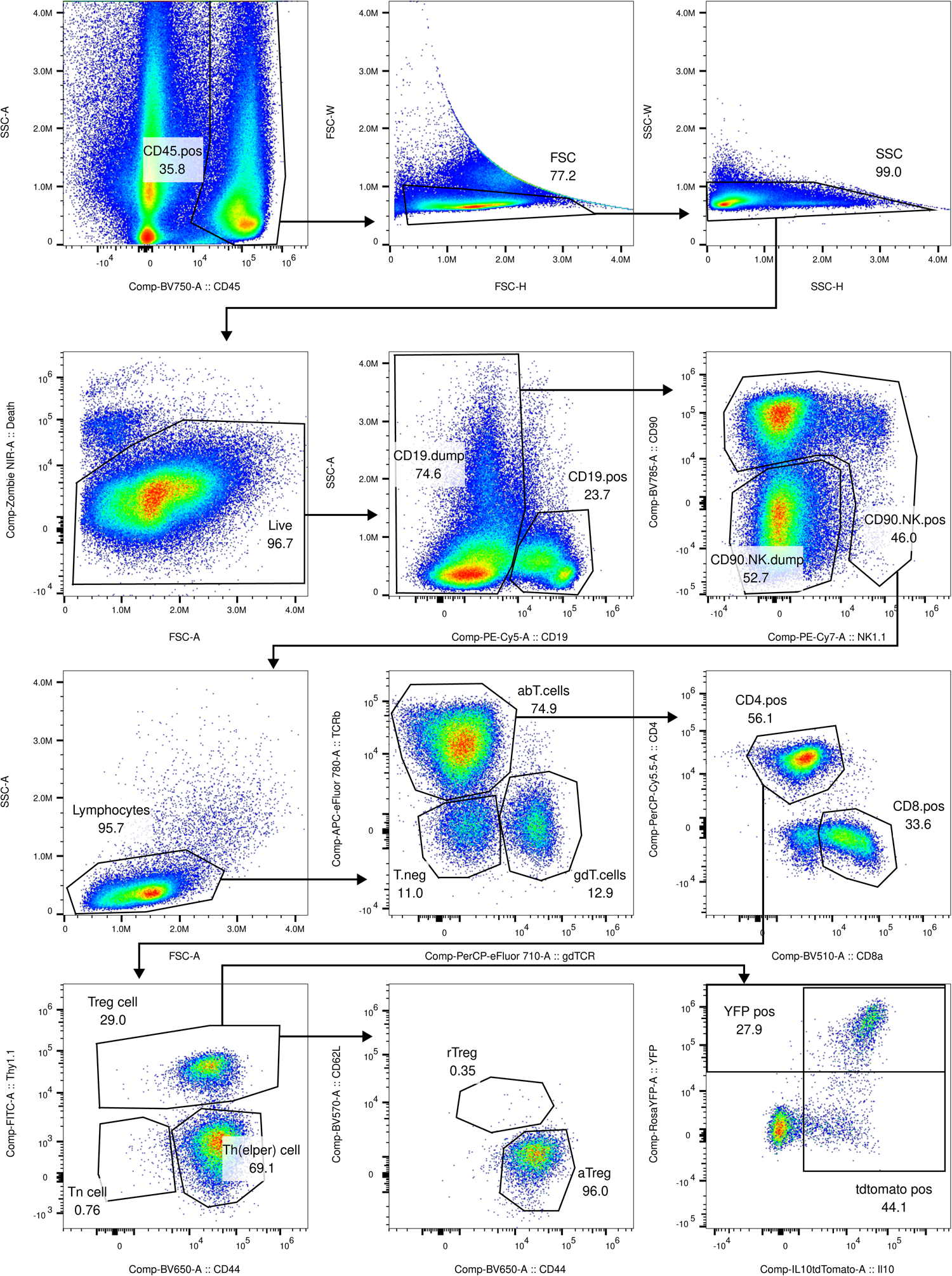
Gating strategy for Treg cells, Thelper cells, CD8 T cells and γδ T cells. Related to Fig. 1 and Extended Data Fig. 1.

**Supplementary Fig. 2.**
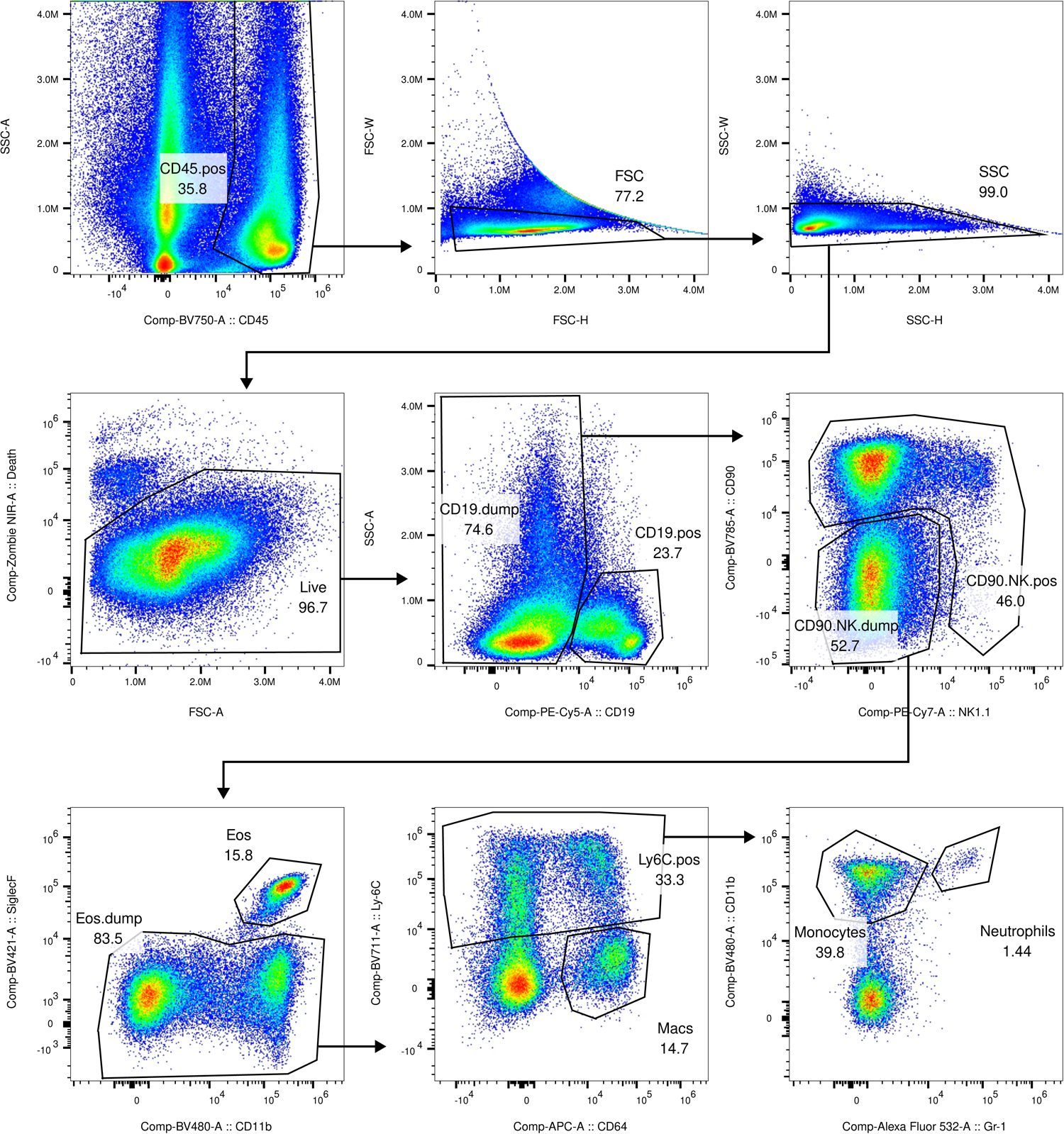
Gating strategy for Macrophages and Monocytes. Related to Fig. 1 and Extended Data Fig. 1.

**Supplementary Fig. 3.**
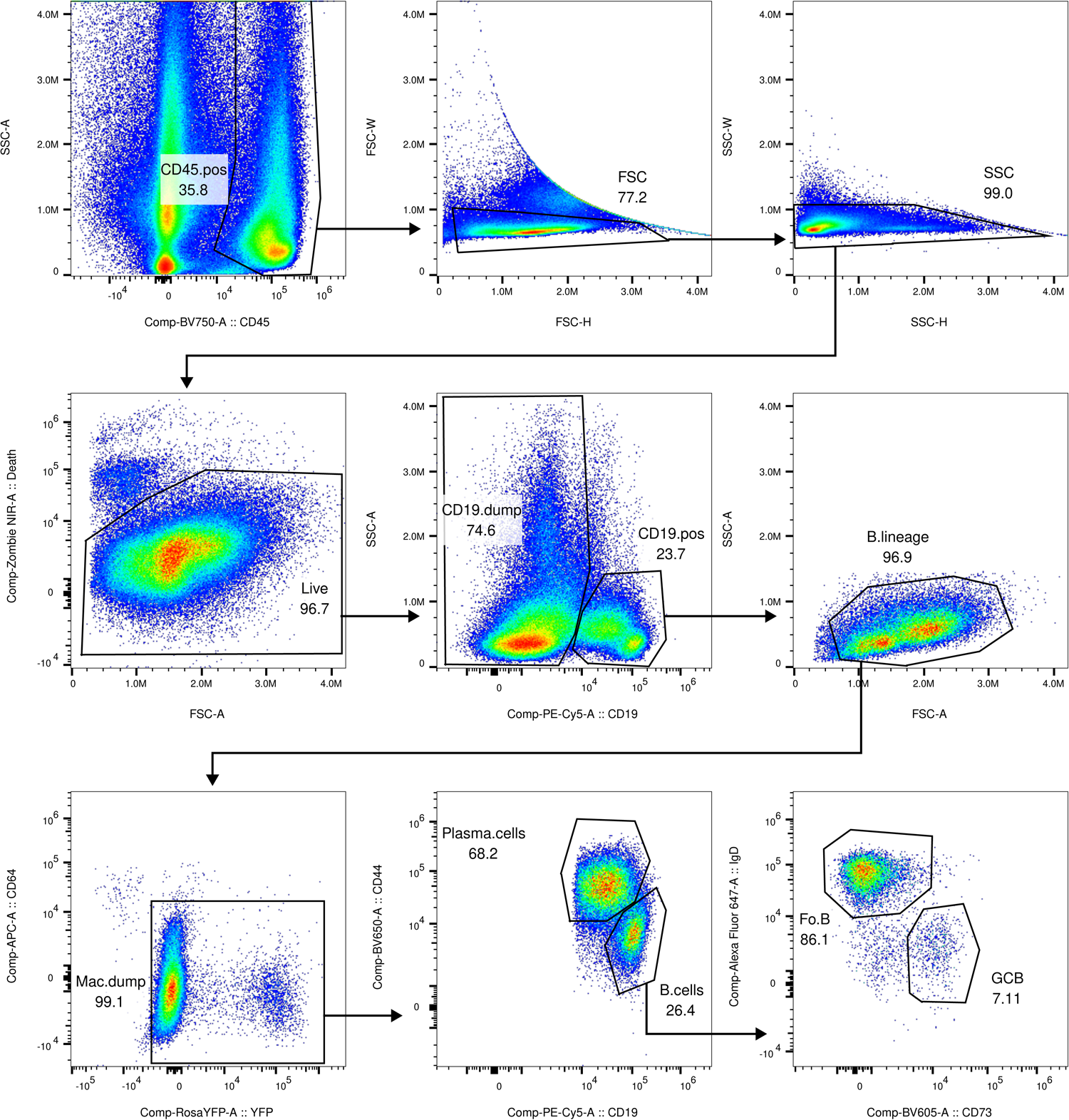
Gating strategy for GC B cells and Plasma cells. Related to Fig. 1 and Extended Data Fig. 1.

**Supplementary Fig. 4.**
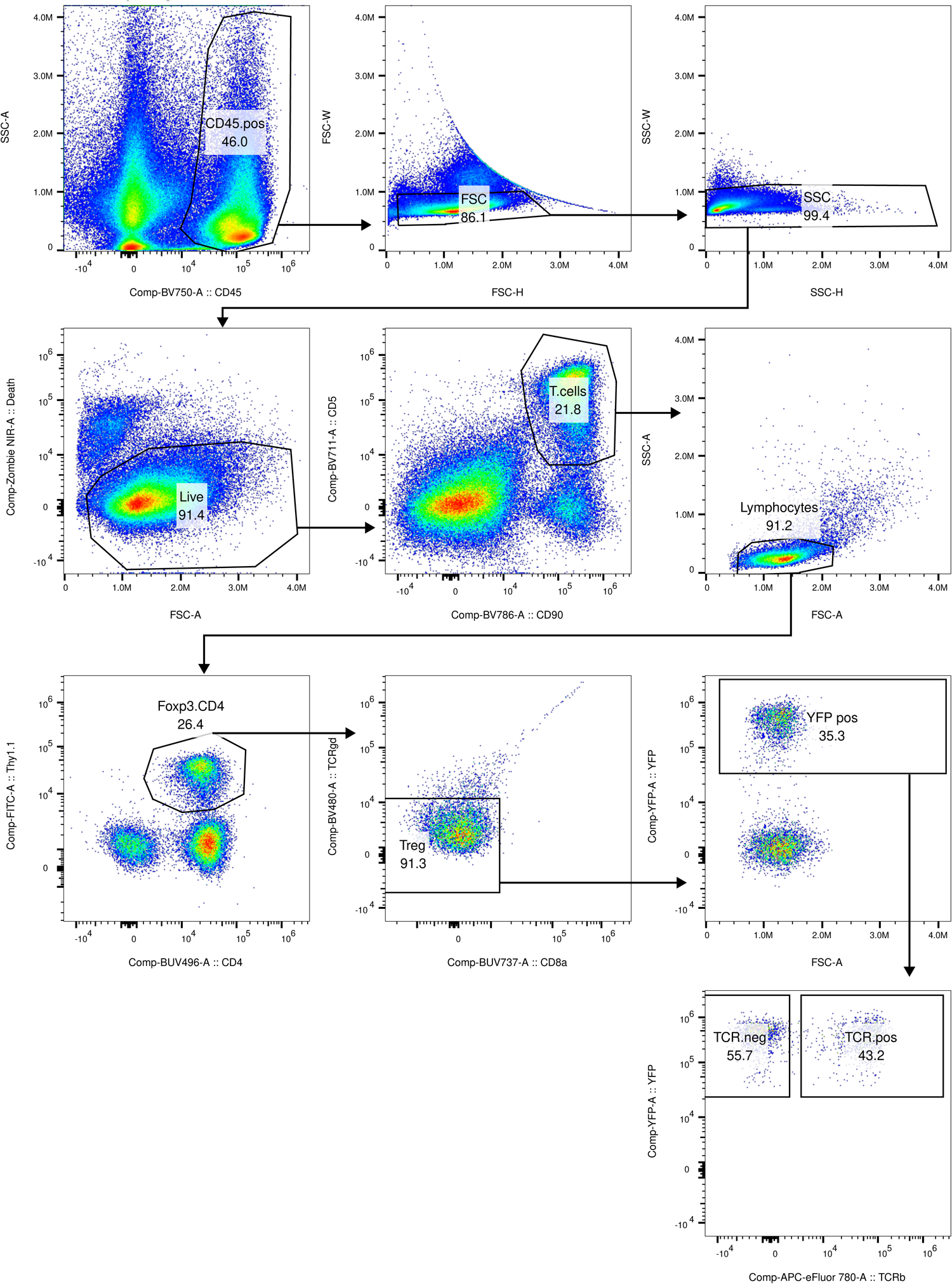
Gating strategy for TCR-deficient Treg cells. Related to Fig. 6.

**Supplementary Fig. 5.**
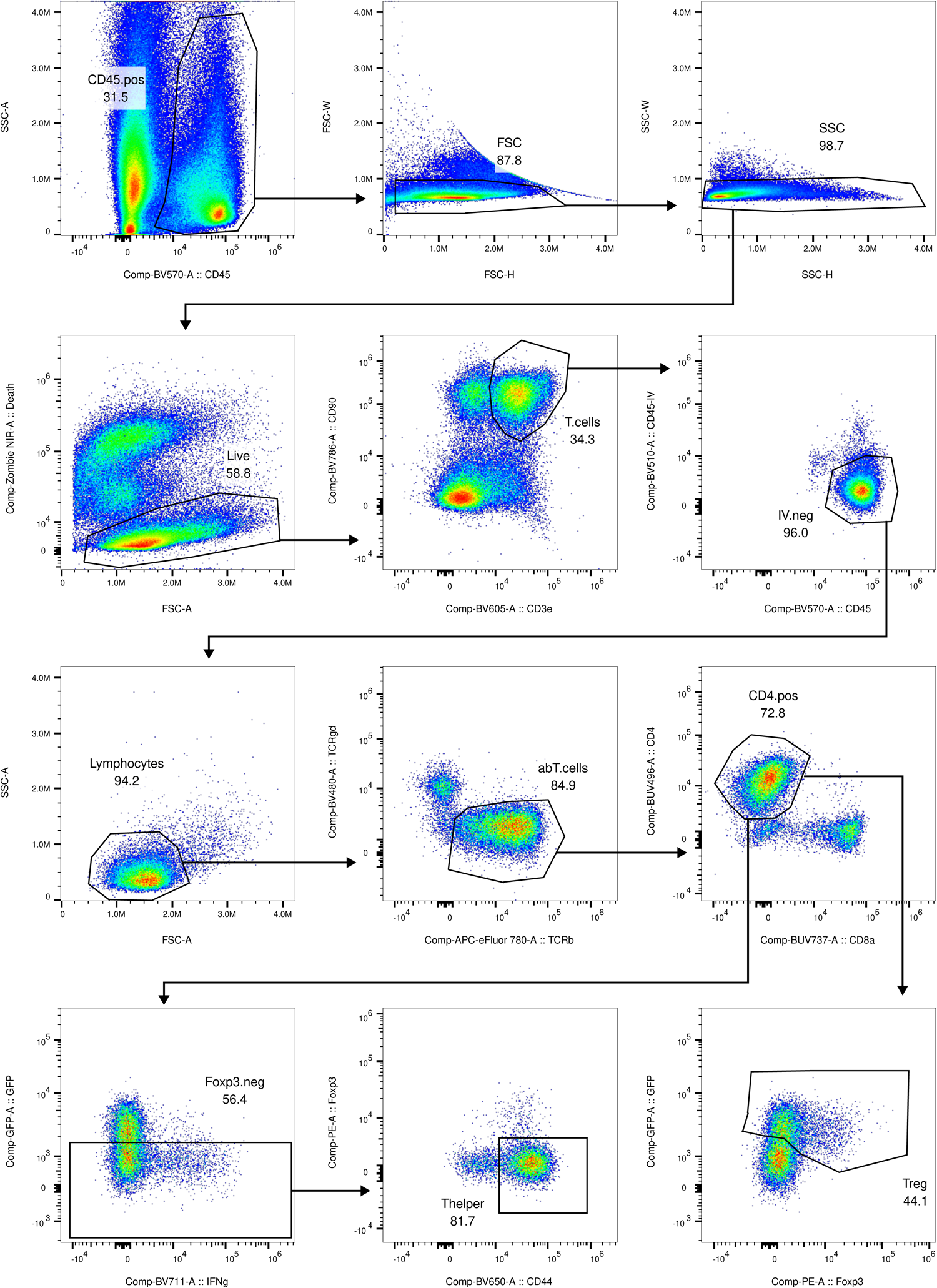
Gating strategy for IL-10 production after re-stimulation. Related to Extended Data Fig. 5.

**Supplementary Fig. 6.**
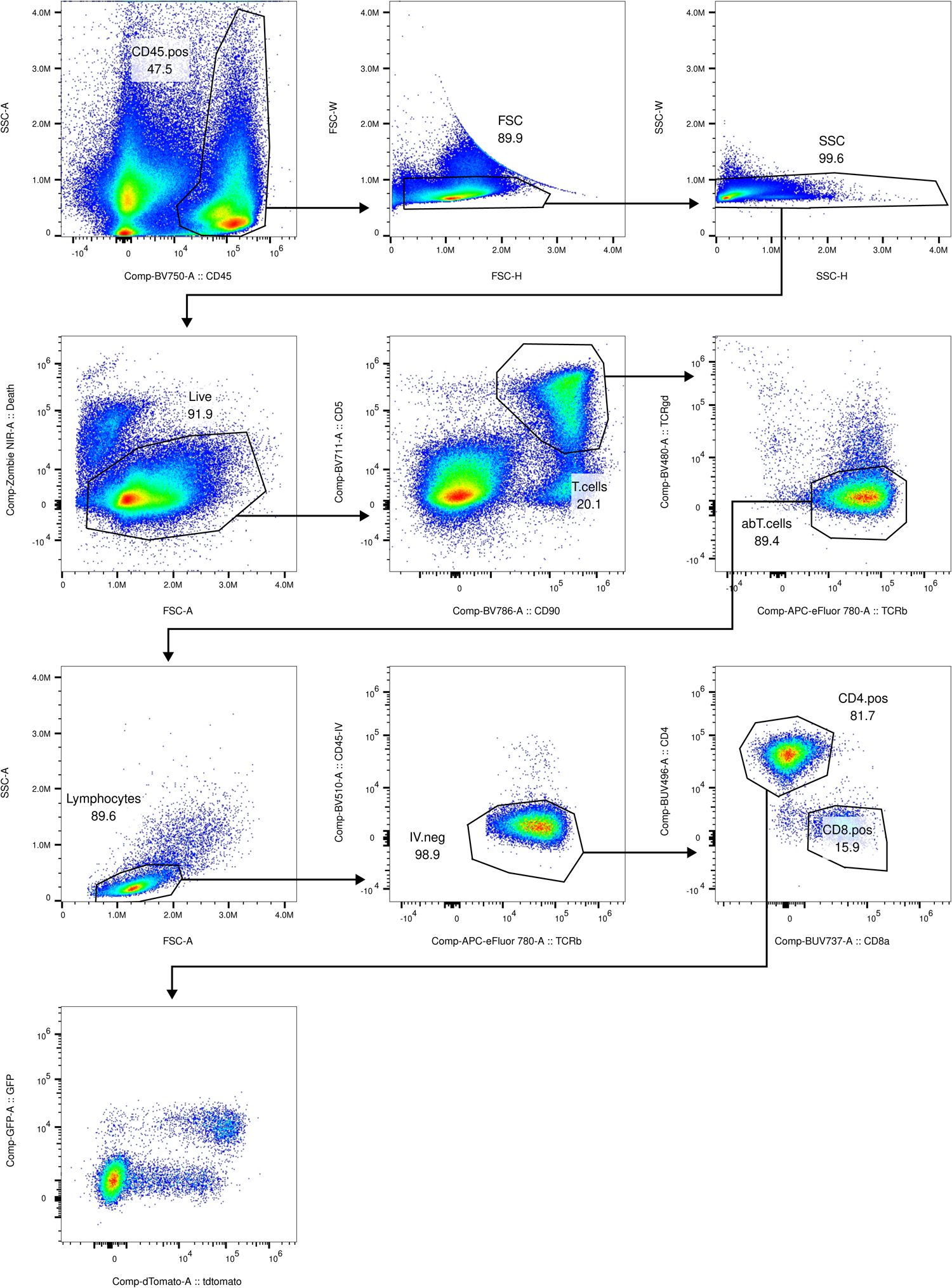
Gating strategy for GFP-DTR and tdTomato expression by CD4 T cells in *Foxp3^Il^*^10^*^-DTR^* mice. Related to Extended Data Fig. 5.

**Supplementary Fig. 7.**
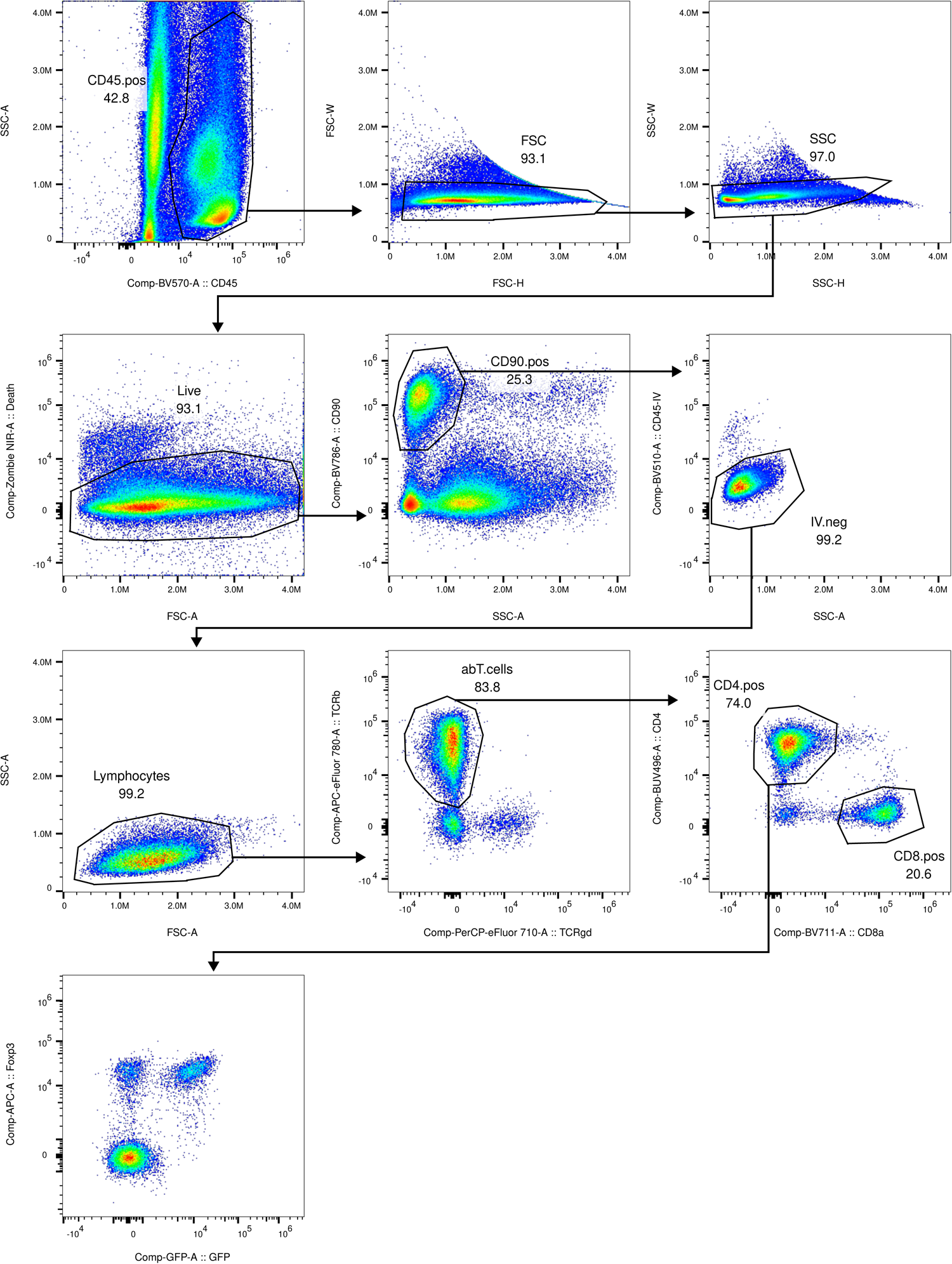
Gating strategy for GFP-DTR expression and Foxp3 staining in CD4 T cells in *Foxp3^Il^*^10^*^-DTR^* mice. Related to Extended Data Fig. 5.

**Supplementary Fig. 8.**
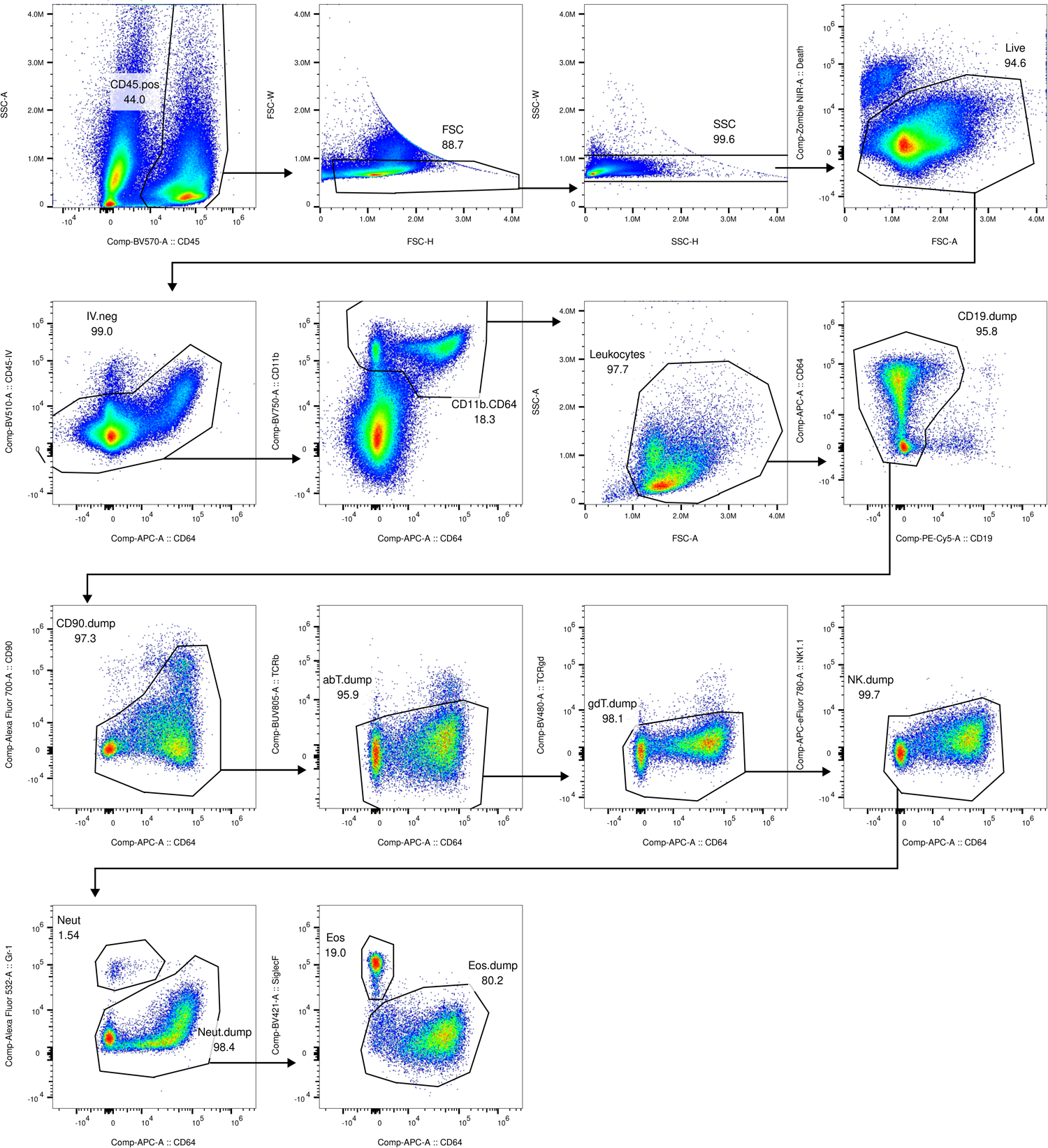
Gating strategy for Neutrophils and Eosinophils. Related to Fig. 7 and Extended Data Fig. 6.

**Supplementary Fig. 9.**
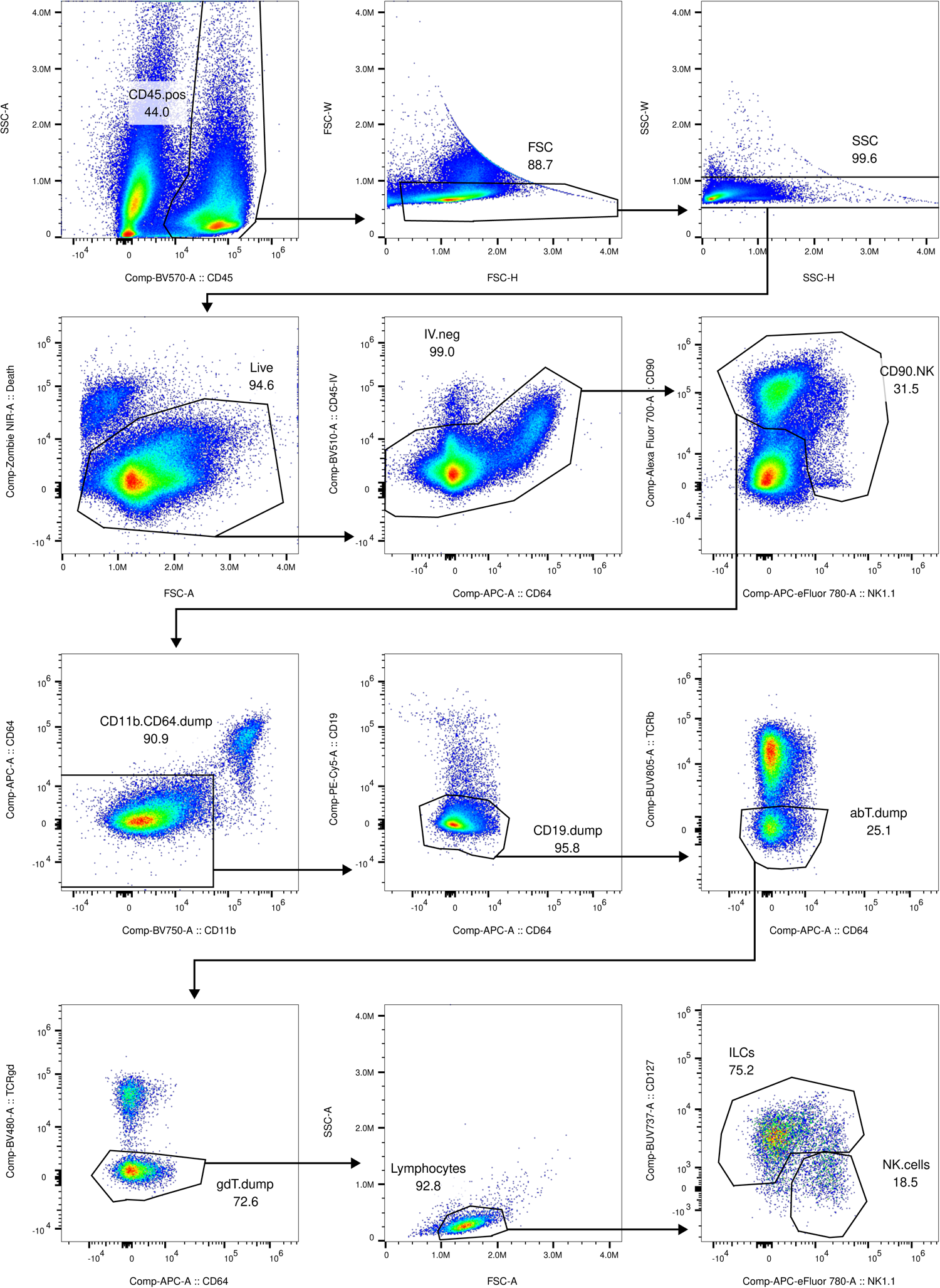
Gating strategy for NK cells. Related to Fig. 7 and Extended Data Fig. 6.

**Supplementary Fig. 10.**
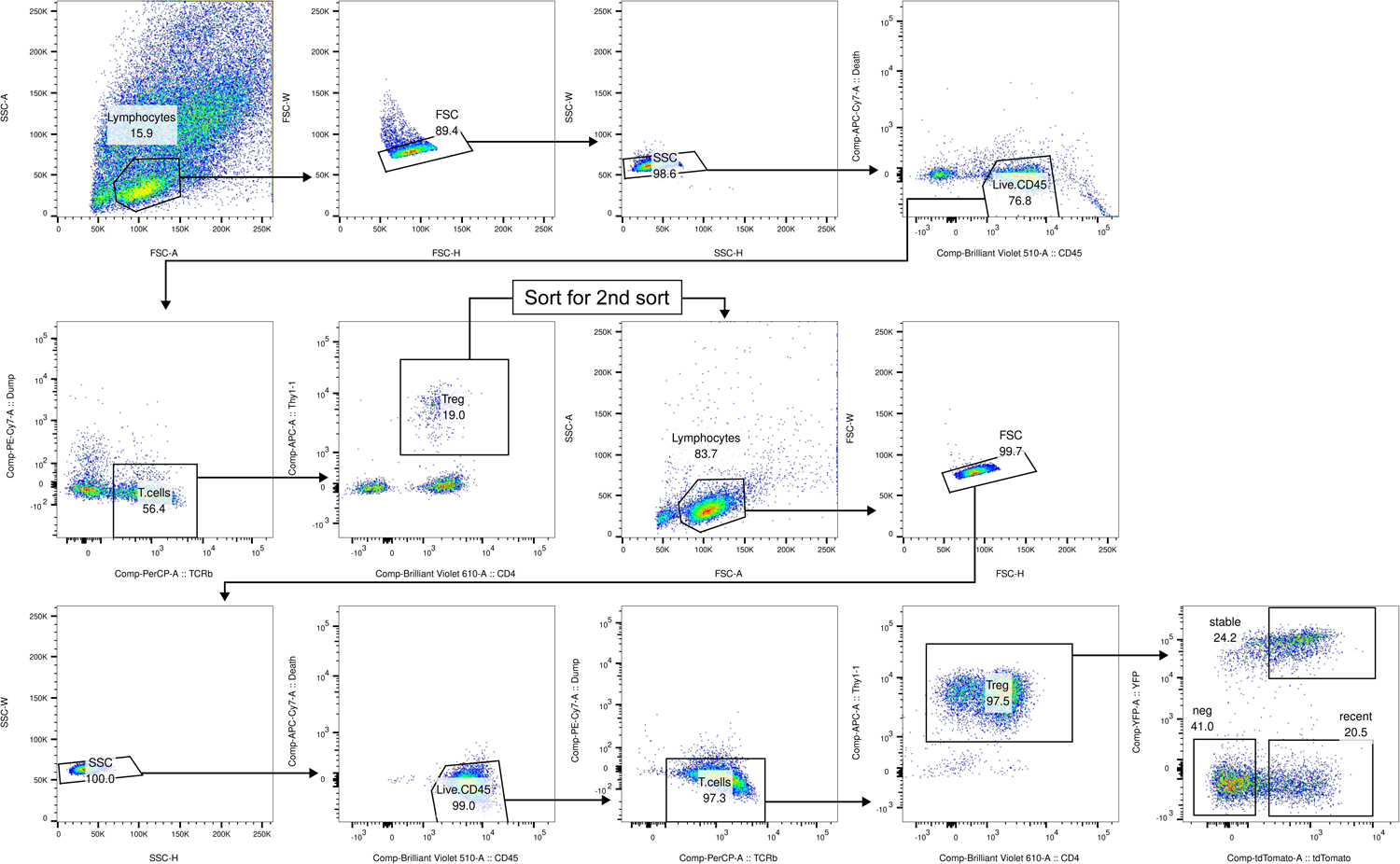
Gating strategy for sorting IL-10 expressing Treg cell populations. Related to all sequencing experiments.

## Tables

**Table S1. Antibodies used for flow cytometry.**

## Methods

### Mice

*Foxp3^Thy1.1^*, *Foxp3^GFP-DTR^*, *Foxp3^CreER-gfp^,* and *Il10^FL^* mice have been previously described and were maintained in house ^33,76–78^. *Gt(ROSA)26Sor^LSL-YFP^* and *Rorc^FL^* have been previously described and were purchased from Jackson Laboratories ^34,79^. *Tcra^FL^ and Maf^FL^* have been previously described ^55,80^. *Maf^FL^* mice were provided by D. R. Littman and *Tcra^FL^* mice were provided by M. Schmidt-Supprian. *Il10^FM^* mice were generated by intercrossing *Foxp3^Thy1.1^*, *Gt(ROSA)26Sor^LSL-YFP^*, and *Il10^tdTomato-CreER^* mice (see below) to homozygosity for each allele. Littermates were used in all experiments and were distributed among experimental groups evenly whenever possible, with different experimental groups co-housed. In experiments with different genotypes, all genotypes were represented in each litter analyzed. All mice were housed at the Research Animal Resource Center for Memorial Sloan Kettering Cancer Center and Weill Cornell Medicine. All studies were under protocol 08-10-023 and approved by the Sloan Kettering Institute Institutional Animal Care and Use Committee. All animals used in this study had no previous history of experimentation and were naive at the time of analysis. Both sexes were used in all experiments unless otherwise noted, as no sex differences in IL-10 expression were detected.

### **Generation of *Il10^tdTomato-CreER^* and *Il10^tdTomato-Cre^* mice.**

*Il10^tdTomato-CreER^* mice were generated by insertion of a targeting construct into the *Il10* locus by homologous recombination in embryonic stem cells on the C57BL/6 background. The targeting construct was generated by inserting sequence containing exons 2– 5 of the *Il10* gene into a plasmid backbone containing a PGK promoter driving expression of diphtheria toxin A subunit (DTA) followed by BGHpA sequence (modified PL452 plasmid). A SalI restriction enzyme site was simultaneously engineered into the *Il10* 3′ UTR between the stop codon and the polyadenylation site. The Clontech Infusion HD Cloning system was used to generate in the pUC19 plasmid backbone sequence encoding (in order from 5′ to 3′) encephalomyocarditis virus IRES; tdTomato; T2A self-cleaving peptide from *Thosea asigna* virus; Cre recombinase fused to the estrogen receptor ligand binding domain (CreER); followed by a FRT site-flanked PGK-Neomycin resistance gene (Neo)-BGHpA cassette. The IRES-tdTomato-T2A-CreERT2-FRT-Neo-BGHpA-FRT sequence was PCR-amplified and inserted into the SalI site in the *Il10* 3′ UTR in the modified PL452 backbone. The resulting plasmid was linearized with the restriction enzyme NotI before electroporation into embryonic stem cells. *Il10^tdTomato-CreER^* mice were bred to *Gt(ROSA)26Sor^FLP^*^1^ mice (MSKCC Mouse Genetics Core) to excise the Neo cassette and backcrossed to C57BL/6 mice to remove the *Gt(ROSA)26Sor^FLP^*^1^ allele. *Il10^tdTomato-Cre^* mice were generated in an identical manner except that the targeting vector contained a codon-optimized NLS-Cre encoding sequence after the T2A.

### Generation of *Foxp3^LSL-DTR^* mice

*Foxp3^LSL-DTR^* mice were generated by Biocytogen. First, a guide RNA targeting the 3’UTR of the *Foxp3* gene was designed and validated (GGAAAGTTCACGAATGTACCA). Then, a targeting vector was constructed including 1400bp homology upstream and downstream of an SspI site in the *Foxp3* 3’UTR. The following sequence was inserted into the SspI site: loxP-IRES-thy1.1-pA-loxP-IRES-DTR-eGFP. Cas9 protein, in vitro transcribed sgRNA, and the targeting vector were then micro-injected into C57BL/6N zygotes. Founder pups were then bred, and confirmed to have the proper integration by PCR and Southern Blot analysis.

### Mouse treatments

For tamoxifen treatment, mice were gavaged with 8 mg tamoxifen dissolved in 200 μL corn oil (Sigma). Tamoxifen was dissolved by gentle agitation at 37°C overnight. Aliquots were frozen (–80°C) and thawed as needed throughout an experiment. We found that freezing and storing at –80°C rather than –20°C greatly reduced the tendency of the tamoxifen to precipitate when thawed. For *Il10^iΔTCR^* experiments, mice were treated on days 0 or 11 and analyzed on day 21. For *Foxp3^iΔIl10^* experiments, mice were treated on days 0 and 2 and analyzed on day 16. For *Il10^iΔRorc^*, *Il10^iΔMaf^*, and *Foxp3^iΔIl10^* (long term) experiments, mice were treated on days 0, 4, 11, 18, 25, and 32 and analyzed on day 35. Diptheria toxin (DT) was reconstituted in sterile PBS at 1 mg/ml and frozen at –80°C in single use aliquots. Aliquots were thawed and diluted in 995 μL PBS. For inactivated control (bDT), this 1 mL dilution was heated at 95 – 100°C for 30 minutes. Both active and control DT were filtered through 0.22 μm syringe-driven filters. Mice were injected intraperitoneally with 200 μL of this dilution for the first two doses (1000 ng DT) or with 200 μL of a 1:1 dilution with PBS for subsequent doses (500 ng DT), except for experiments depicted in Extended Data Fig. 9h-k, where 1000ng DT was administered for each dose. Bleomycin was dissolved in PBS at a concentration of 5.7 U/mL, sterile-filtered, and frozen (–80°C) in single use aliquots. Aliquots were diluted with sterile PBS immediately before use. mice were anesthetized with isofluorane (3% in O_2_, 3 L/min, Covetrus) and 0.1 U bleomycin in 35 μL PBS was administered intranasally using a micropipette. Mice were exposed to isoflurane for at least 5 minutes before delivery of bleomycin, and mouth was gently pressed shut during delivery to prevent swallowing. Bleomycin was given drop-wise, with pauses between drops to ensure inhalation. For antibiotic treatment, a solution of 1 g/L ampicillin sodium salt, 1 g/L kanamycin sulfate, 0.8 g/L vancomycin hydrochloride, 0.5 g/L metronidazole, and 2.5 g/L sucralose (Splenda) was prepared in acidified, reverse-osmosed water and sterile-filtered. Control solution only contained sucralose but was otherwise treated similarly. Solutions were replaced every seven days for the duration of the experiment. For chemically induced colitis (Figure 1), 15 g dextran sulfate sodium salt (DSS, MW ∼ 40,000) was dissolved in 50 mL distilled deionized water, and then sterile filtered. This solution was then diluted in 450 mL acidified, reverse-osmosed water, resulting in a final concentration of 3% (w/v) DSS. Control groups received 50 mL sterile filtered distilled deionized water diluted into 450 mL acidified, reverse-osmosed water. For these experiments, female mice were used, since male mice proved highly sensitive to even lower concentrations of DSS ^81^. For chemically induced colitis (Extended Data Fig. 9), 7.5 g dextran sulfate sodium salt (DSS, MW ∼ 40,000) was dissolved in 50 mL distilled deionized water, and then sterile filtered. This solution was then diluted in 450 mL acidified, reverse-osmosed water, resulting in a final concentration of 1.5% (w/v) DSS. Solution was replaced after seven days.

### Cell isolation for flow cytometry

Mice were injected retro-orbitally with 1.5 μg anti-mouse CD45.2 (Brilliant Violet 510 conjugated, BioLegend 109838) in 200 μL sterile PBS 3 min prior to euthanasia to label and exclude blood-exposed cells. All centrifugations were performed at 800x g for 3 min at 4 °C. Secondary lymphoid organs were dissected and placed in 1 mL wash medium (RPMI 1640, 2% Fetal Bovine Serum (FBS), 10 mM HEPES buffer, 1% penicillin/streptomycin, 2 mM L-glutamine). Tissues were then mechanically disrupted with the back end of a syringe plunger, and then passed through a 100 μm, 44% open area nylon mesh. For skin and lung, both ears and all lung lobes were collected. Ears were peeled apart to expose the dermis and cut into 6 total pieces. Tissues were then placed in 5 mL snap-cap tubes (Eppendorf 0030119401) in 3 mL wash medium supplemented with 0.2 U/mL collagenase A, 5 mM calcium chloride, 1 U/mL DNase I, along with three ¾ inch ceramic beads (MP Biomedicals, 116540424-CF) and shaken horizontally at 250 RPM for 45 min at 37°C for the lung and for two rounds of 25 min for skin, replacing collagenase solution in between. Digested samples were then passed through a 70 μm strainer (Milltenyi Biotec, 130-095-823) and centrifuged to remove collagenase solution. Lungs were were then treated with ACK buffer (155 mM ammonium chloride, 10 mM potassium bicarbonate, 100 nM EDTA pH 7.2) to lyse red blood cells, and then washed by centrifugation in 40% Percoll^TM^ (ThermoFisher, 45-001-747) in wash medium to remove debris and enrich for lymphocytes. For colon, cecum and large intestine were dissected, and after removal of fat and the cecal patch, opened longitudinally and vigorously shaken in 1x PBS to remove luminal contents. Colon was then cut into 1 to 2 cm pieces, placed in a 50 mL screw-cap tube with 25 mL wash medium supplemented with 5 mM EDTA and 1 mM dithiothreitol and shaken horizontally at 250 RPM for 15 to 20 min at 37°C. After a 5 sec vortex, epithelial and immune cells from the epithelial layer were removed by filtering the suspension through a tea strainer. Remaining tissue was placed back in 50 mL tubes, washed with 25 mL wash medium, strained again, and replaced in 50 mL tubes. 25 mL wash medium supplemented with 0.2 U/mL collagenase A, 4.8 mM calcium chloride, 1 U/mL DNase I was added, along with four ¾ inch ceramic beads, and tissues were shaken horizontally at 250 RPM for 35 min at 37°C. Suspension was then passed through a 100 μm strainer, centrifuged to remove debris and collagenase solution, and then washed by centrifugation in 40% Percoll^TM^ in wash medium. Small intestine was processed as the colon, except that the pieces were cleaned by shaking in corn starch, then rinsed with PBS before EDTA treatment. All enzymatically digested samples were washed by centrifugation in 5 mL wash medium.

### Flow cytometry

To assess cytokine production after *ex vivo* restimulation, single cell suspensions were incubated for 4 hours at 37°C with 5% CO_2_ in the presence of 50 ng/mL PMA and 500 ng/mL ionomycin with 1 μg/mL brefeldin A and 2 μM monensin to inhibit ER and Golgi transport. For flow cytometric analysis, cells were stained in 96 well V-bottom plates with antibodies and reagents used at concentrations indicated in Table S1. All centrifugations were performed at 900x g for 2 min at 4 °C. Staining with primary antibodies was carried out in 100 μL for 25 min at 4°C in staining buffer (PBS, 0.2 % (w/v) BSA, 2 mM EDTA, 10 mM HEPES, 0.1% (w/v) NaN_3_). Cells were then washed with 200 μL PBS and then concurrently stained with Zombie NIR™ Fixable Viability dye and treated with 20 U/mL DNase I in DNase buffer (2.5 mM MgSO_4_, 0.5 mM CaCl_2_, 136.9 mM NaCl, 0.18 mM Na_2_HPO_4_, 5.36 mM KCl, 0.44 mM KH_2_PO_4_, 25 mM HEPES) for 10 min at room temperature. Cells were washed with 100 μL staining buffer, resuspended in 200 μL staining buffer, and passed through a 100 μm nylon mesh. For cytokine staining, cells were fixed and permeabilized with BD Cytofix/Cytoperm per manufacturer instructions. Intracellular antigens were stained overnight hour at 4°C in the 1x Perm/Wash buffer. Samples were then washed twice in 200 μL 1x Perm/Wash buffer, resuspending each time, resuspended in 200 μL staining buffer, and passed through a 100 μm nylon mesh. All samples were acquired on an Aurora cytometer (Cytek Biosciences) and analyzed using FlowJo v10 (BD Biosciences).

### Histopathological analysis

∼1cm sections of colon were fixed in 4% PFA for > 48 hours. Tissue embedding, sectioning, and staining was carried out by Histowiz Inc (New York). A blinded pathologist scored sections based on the Simplified Geboes Score rubric^63^:

Grade 0 (no inflammatory activity):

0.0 No abnormalities
0.1 Presence of architectural changes
0.2 Presence of architectural changes and chronic mononuclear cell infiltrate

Grade 1 (Basal plasma cells):

1.0 No increase
1.1 Mild increase
1.2 Marked increase

Grade 2A (Eosinophils in lamina propria):

2A.0 No increas
2A.1 Mild increase
2A.2 Marked increase

Grade 2B (Neutrophils in lamina propria):

2B.0 No increase
2B.1 Mild increase
2B.2 Marked increase

Grade 3 (Neutrophils in epithelium):

3.0 None
3.1 < 50% crypts involved
3.2 > 50% crypts involved

Grade 4 (Epithelial injury in crypt and surface epithelium):

4.0 None
4.1 Marked attenuation
4.2 Probable crypt destruction: probable erosions
4.3 Unequivocal crypt destruction: unequivocal erosion
4.4 Ulcer or granulation tissue

### Flow cytometric identification of immune cell populations

Generally, the following populations were identified with the following markers (all immune cells were first gated as CD45^+^ and ZombieNIR^−^ and excluded of doublets):

Treg cells: CD90.2^+^CD5^+^SSC^lo^FSC^lo^TCRβ^+^TCRγδ^−^CD4^+^CD8α^−^Thy1.1^+^
Th cells (CD44^hi^CD4^+^ cells): CD90.2^+^CD5^+^SSC^lo^FSC^lo^TCRβ^+^TCRγδ^−^CD4^+^CD8α^−^Thy1.1^−^CD44^+^
Macrophages: CD11b^+^CD64^+^CD90.2^−^CD19^−^NK1.1^−^Gr-1^−^SiglecF^−^Ly6C^−/lo^
Monocytes: CD11b^+^CD64^−/lo^CD90.2^−^CD19^−^NK1.1^−^Gr-1^−^SiglecF^−^Ly6C^hi^
Plasma cells: CD19^lo^CD44^hi^CD64^−^CD11b^−/lo^CD11c^−/lo^CD90.2^−^NK1.1^−^SiglecF^−^Gr-1^−^
Germinal center B cells: CD19^hi^CD44^lo^CD73^+^IgD^−^CD64^−^CD11b^−/lo^CD11c^−/lo^ CD90.2^−^NK1.1^−^SiglecF^−^Gr-1^−^
γδT cells: CD90.2^+^SSC^lo^FSC^lo^TCRβ^−^TCRγδ^+^
CD8^eff^ cells (CD44^hi^CD8^+^ T cells): CD90.2^+^SSC^lo^FSC^lo^TCRβ^+^TCRγδ^−^CD4^−^CD8α^+^CD44^hi^CD62L^−^
NK cells: NK1.1^+^CD90.2^+/–^SSC^lo^FSC^lo^TCRβ^−^TCRγδ^−^CD19^−^CD64^−^CD11b^−^CD127^−^
Neutrophils: Gr-1^+^CD11b^+^CD64^−/lo^CD90.2^−^CD19^−^NK1.1 SiglecF^−^
Eosinophils: SiglecF^+^CD11b^+^CD64^−/lo^CD90.2^−^CD19^−^NK1.1^−^

### Cell sorting for sequencing

Cell isolation was performed as described above, except that samples were not washed with 40% Percoll^TM^. Staining was performed as described above, except buffer contained 2 mM L-glutamine and did not contain NaN_3_, with staining volume adjusted to 500 μL and washes adjusted to 5 mL and staining performed in 15 mL screw-cap tubes. 1 μg ‘HashTag’ antibodies (BioLegend, 155801, 155803, 155805, 155807) were added to extracellular antigen stain for scRNA-seq sorting, and scRNA-seq sort samples were not DNase I treated. Samples were resuspended in Wash buffer supplemented 5mM EDTA for sorting. Samples were double sorted, with the first sort enriching for all Thy1.1^+^ Treg cells, and the second sort separating *Il10^neg^*, *Il10^recent^*, and *Il10^stable^*, or tdTomato^+^ versus tdTomato^−^ cells. For bulk RNA-seq, samples were sorted directly into Trizol-LS^TM^, per the manufacturer’s instructions, in 1.5 mL microcentrifuge tubes. For scRNA-seq, samples were sorted into PBS with 0.04% BSA (w/v) in 1.5 mL Protein LoBind tubes (Eppendorf, 0030108442). For ATAC-seq, samples were sorted into wash medium in 1.5 mL Protein LoBind tubes. All sorting was performed on an Aria II (BD Biosciences).

### Preparation of reference genome

The mm39 mouse genome assembly and NCBI RefSeq annotation information (GTF file) were downloaded from the UCSC Genome browser ^82–85^. In order to account for the presence of the *Il10^tdTomato-CreER^*, *Foxp3^Thy1.1^*, and *Gt(ROSA)26Sor^LSL-YFP^* targeted mutations, the corresponding sequences were inserted into the appropriate locations of the mm39 genome using the ‘reform’ script, creating the ‘reformed mm39’ genome ^86^. The GTF file was modified to appropriately extend the *Il10*, *Foxp3*, and *Gt(ROSA)26Sor* transcript and gene annotations, and to shift all other affected annotations, resulting in a ‘reformed GTF’ using a custom R script, relying on the ‘GenomicRanges’ and ‘rtracklayer’ packages ^87–89^. The reformed mm39 and reformed GTF were used for bulk RNA-seq and ATAC-seq alignment and analyses after generating a STAR genome index using STAR (version 2.7.3a) with the following commands ^90^.

ATAC:

STAR --runMode genomeGenerate --runThreadN 4 --genomeDir mm39_100 --genomeFastaFiles mm39_reformed.fa

RNA:

STAR --runMode genomeGenerate --runThreadN 4 --genomeDir mm39_100_RNA --genomeFastaFiles mm39_reformed.fa --sjdbGTFfile mm39_reformed.gtf

### Bulk RNA-seq

5,000 cells were sorted per population per replicate for bulk RNA-seq, with each replicate pooled from two mice. RNA was extracted and libraries prepared using SMARTer Stranded RNA-Seq Kits according the manufacturer’s protocols (Takara) by the Integrated Genomics Operation (IGO) Core at MSKCC. Paired-end 50bp reads (20 to 30 million per sample) were sequenced on an Illumina HiSeq 3000 by IGO.

### Bulk RNA-seq data processing

Samples were processed and aligned using Trimmomatic (version 0.39), STAR (version 2.7.3a), and samtools (version 1.12), with the following steps, where *Sample* stands in for each *Il10^neg^*, *Il10^recent^*, or *Il10^stable^* replicate ^90–92^.

TrimmomaticPE *Sample*_R1.fastq.gz *Sample*_R2.fastq.gz -baseout *Sample*.fastq.gz

ILLUMINACLIP:TruSeq3-PE.fa:2:30:10 LEADING:3 TRAILING:3 SLIDINGWINDOW:4:15 MINLEN:36

STAR --runThreadN 6 --runMode alignReads --genomeLoad NoSharedMemory --readFilesCommand zcat --genomeDir mm39_100_RNA --readFilesIn *Sample*_1P.fastq.gz *Sample*_2P.fastq.gz -- outFileNamePrefix *Sample* --outSAMtype BAM Unsorted --outBAMcompression 6 -- outFilterMultimapNmax 1 --outFilterMismatchNoverLmax 0.06 --outFilterMatchNminOverLread 0.35 -- outFilterMatchNmin 30 --alignEndsType EndToEnd

samtools sort -@ 4 -n -o *Sample*.bam *Sample*Aligned.out.bam

samtools fixmate -@ 4 -rm *Sample*.bam *Sample*.fixmate.bam

samtools sort -@ 4 -o *Sample*.resort.bam *Sample*.fixmate.bam

samtools markdup -@ 4 -l 1500 -r -d 100 -s *Sample*.resort.bam *Sample*.duprm.bam

samtools index -@ 4 -b *Sample*.duprm.bam

This procedure resulted in retention of all uniquely aligning reads, with PCR and optical duplicates removed, to be used for downstream analysis. Reads aligning to genes derived from the reformed GTF were then counted using a custom R script relying on the ‘GenomicAlignments’, ‘GenomicRanges’, and ‘GenomicFeatures’ packages with default counting parameters ^88^. Differential expression analysis was carried out using the ‘DESeq2’ package, with the formula ‘∼ Celltype + Replicate’ where Celltype was either *Il10^neg^*, *Il10^recent^*, or *Il10^stable^* and replicates were the separate samples from which each of the three populations were sorted ^93^. Fragments per kilobase mapped (FPKM) normalized counts were extracted using the *fpkm* function of DESeq2. Differential expression analysis and statistical testing was performed for all pairwise comparisons of ‘Celltype’: *Il10^neg^*, *Il10^recent^*, *Il10^stable^*. Differential expression analysis was performed on all genes, but genes with FPKM counts below the mean FPKM count of *Cd8a* (a gene functionally not expressed in Treg cells), genes with 0 counts in the majority of samples, or genes corresponding to immunoglobulin or TCR variable, diversity, or junction segments were eliminated for subsequent analyses. This process did not remove any significantly differentially expressed genes, except immunoglobulin or TCR variable, diversity, or junction segments, whose differential expression was not interpretable. K-means clustering was performed with R using per-gene Z-score normalized counts of genes differentially expressed (adjusted *p*-value < 0.05) in any pairwise comparison between the three cell populations, with 7 clusters chosen on the basis of preliminary hierarchical clustering. TCR activated and repressed genes were defined as genes which lost and gained expression in Treg cells ablated of the *Tcra* gene ^52^.

### scRNA-seq

Uniquely ‘hash-tagged’ samples from different tissues were pooled during sorting as separate tdTomato^+^ and tdTomato^−^ samples. The tdTomato^+^ sample had 48,000 cells (35,000 from LILP, 1,100 from lung, 11,000 from mesLN, and 900 from medLN) and the tdTomato^−^ sample had 60,000 cells (30,000 from LILP, 10,000 each from lung, mesLN, and medLN). Samples were centrifuged and resuspended in 30 μL PBS with 0.04% BSA (w/v). Libraries were then prepared following the 10x Single Cell 3’ Reagent Kit v3 (10X Genomics) following the manufacturer’s instructions, incorporating the BioLegend TotalSeq^TM^-A HTO protocol. Samples were sequenced on an Illumina NovaSeq platform by IGO.

### scRNA-seq processing

Reads from the tdTomato^+^ and tdTomato^−^ samples were processed, aligned to the mm10 genome, and demultiplexed using Cell Ranger software (10X Genomics) with default parameters. Reads for the HTOs of the tdTomato^+^ and tdTomato^−^ samples were processed and demultiplexed using Cell Ranger software with default parameters. Samples were further processed and analyzed with a custom R script relying on the ‘Seurat’ (version 4) package ^94^. First, genes detectable in fewer than 0.1% of cells were removed. Second, HTO identities (i.e. LILP, Lung, mesLN, medLN) were assigned using the *HTODemux* function and cells without an unambiguous HTO identity or determined to be a doublet were excluded ^95^. Then, cells with mitochondrial genes accounting for > 10% of gene counts, presumed to be dead or dying, as well as cells in the top or bottom 2% of total counts were eliminated. This latter cut off was chosen on the basis of a percentile rather than an arbitrary absolute value to account for different cell numbers, and different median UMI counts across the two samples. Afterwards, tdTomato^+^ and tdTomato^−^ samples were merged and analyzed together. First, the top 2000 variable genes were identified and scaled. A PCA was performed on these genes and the top 30 principal components were used to assign k-nearest neighbors, generate a shared nearest neighbor graph, and then optimize the modularity function to determine clusters, at resolution = 0.5 ^96^. On the basis of these original clusters, a small population of cells dominated by high type I interferon signaling was excluded and subsequent analyses were performed only on cells with LILP or mesLN HTO identities. The remaining cells had gene counts scaled again. A PCA was performed on the 3000 most variable genes and the top 30 principal components were used to assign k-nearest neighbors, generate a shared nearest neighbor graph, and then optimize the modularity function to determine clusters, at resolution = 0.5. The shared nearest neighbor graph was used as input for the python-based algorithm ‘Harmony’ (600 iterations) in order to generate a two dimensional force directed layout for visualization ^97^. The 30 PCs were used as input for the python-based algorithm ‘Palantir’ in order to determine ‘pseudotime’ and ‘entropy’ values ^42^. In order to reconcile scRNA-seq clusters and bulk RNA-seq populations, the ‘Seurat’ function *AddModuleScore* was used to assign scores for each bulk gene cluster to each cell. Mean scores for every cell cluster were calculated. For enrichment testing, the phyper function of R was used, with q = genes in a given bulk RNA-seq k-means cluster and also significantly over- or under-expressed in a given scRNA-seq subset (combination of cell cluster and tdTomato^+^ or tdTomato^−^ identity); m = genes in a given bulk RNA-seq k-means cluster; n = all other genes with detectable expression in a given scRNA-seq subset; and k = all genes significantly over- or under-expressed in a given scRNA-seq subset.

### ATAC-seq

40,000 cells were sorted per population per replicate for ATAC-seq with replicates 1 and 2 originating from a single mouse each and replicate 3 representing two pooled mice. ATAC-seq libraries were prepared as previously described, with some modifications ^98^. Cells were pelleted in a fixed rotor benchtop centrifuge at 500x g for 5 min at 4°C. Cells were then washed in 1 mL cold PBS and pelleted again. Supernatant was aspirated and cells were resuspended in 50 µL ice-cold cell lysis buffer (10 mM Tris-Cl pH 7.4; 10 mM NaCl; 3 mM MgCl_2_; 0.1% NP-40) to disrupt plasma membranes. Nuclei were pelleted at 1000x g for 10 min. Supernatant was aspirated and nuclei were resuspended in 40 µL transposition reaction mixture (Illumina Tagment Kit: 20 µL TD buffer; 2 µL TDE1; 18 µL ddH_2_O). Samples were incubated in a ThermoMixer at 1100 RPM for 45 min at 42°C. DNA was then purified usng a MinElute Reaction Cleanup Kit, according to the manufacturer’s instructions. DNA was eluted in 10 µL buffer EB. Libraries were then barcoded and amplified with NEBNext® High-Fidelity Master Mix and primers from ^99^ (50 µL reaction with 10 µL DNA and 2.5 µL of 25 µM primers – 1 cycle of 5 min at 72°C, 30 sec at 98°C; 5 cycles of 10 sec at 98°C, 20 sec at 63°C, 1 min at 72°C). A qPCR on the product determined that an additional 7 cycles (10 sec at 98°C, 20 sec at 63°C, 1 min at 72°C) were required. Library was purified and size selected with AMPure XP beads: 45 µL of PCR product was incubated with 18 µL beads and supernatant was collected (beads bound larger > ∼2000 bp fragments). Supernatant (63 µL) was then incubated with an additional 63 µL of beads for 5 minutes, supernatant was removed, beads were washed twice with 75% ethanol, and DNA was eluted into 50 µL H_2_O by incubating for 2 minutes. Samples were QC-checked and quantified on an Agilent BioAnalyzer by IGO. Paired-end 50bp reads, 20 to 30 million per sample, were sequenced on an Illumina HiSeq 3000 by IGO.

### ATAC-seq data processing

Samples were processed and aligned using Trimmomatic (version 0.39), STAR (version 2.7.3a), and samtools (version 1.12), with the following steps, where *Sample* stands in for each *Il10^neg^*, *Il10^recent^*, or *Il10^stable^* replicate.

TrimmomaticPE *Sample*_R1.fastq.gz *Sample*_R2.fastq.gz -baseout *Sample*.fastq.gz

ILLUMINACLIP:TruSeq3-PE.fa:2:30:10 LEADING:3 TRAILING:3 SLIDINGWINDOW:4:15 MINLEN:36

STAR --runThreadN 6 --runMode alignReads --genomeLoad NoSharedMemory --readFilesCommand zcat --genomeDir mm39_100 --readFilesIn *Sample*_1P.fastq.gz *Sample*_2P.fastq.gz -- outFileNamePrefix *Sample* --outSAMtype BAM Unsorted --outBAMcompression 6 -- outFilterMultimapNmax 1 --outFilterMismatchNoverLmax 0.06 --outFilterMatchNminOverLread 0.35 --

outFilterMatchNmin 30 --alignIntronMax 1 –alignEndsType Local

samtools sort -@ 4 -n -o *Sample*.bam *Sample*Aligned.out.bam

samtools fixmate -@ 4 -rm *Sample*.bam *Sample*.fixmate.bam

samtools sort -@ 4 -o *Sample*.resort.bam *Sample*.fixmate.bam

samtools markdup -@ 4 -l 1500 -r -d 100 -s *Sample*.resort.bam

*Sample*.duprm.bam samtools sort -@ 4 -n -o *Sample*.byname.bam *Sample*.duprm.bam

samtools index -@ 4 -b *Sample*.duprm.bam

This procedure resulted in retention of all uniquely aligning reads, with PCR and optical duplicates removed, to be used for downstream analysis. Then, peaks were called across the 3 replicates of each cell population individually using Genrich (version 0.5), with the following command, where *Celltype* stands in for each cell population, *Sample* (r1-3) represent the three replicates, and X is 0.002% of the mean number of uniquely aligned reads for each cell population ^100^.

Genrich -t *Sample*_r1.byname.bam,*Sample*_r2.byname.bam,*Sample*_r3.byname.bam - o ./Gen_out/Peak/Celltype.narrowPeak -j -d 25 -g 5 -v -q 0.01 -a X

Peak atlases for each population were then concatenated, sorted, and clustered using bedtools (version 2.27.1) to identify overlapping peaks with the following command ^101^.

bedtools cluster -d -1 -i combined_sort.narrowPeak > clustered.narrowPeak

A custom R script was then used to merge the atlases, according to the following principles. If all the peaks in a cluster entirely overlapped, defined as all peak summits falling within the maximal start and minimal end positions of the cluster, the merged peak was defined as the mean start, summit, and end of all peaks in that cluster. This was the case for ∼82% of all peaks. In the other cases, clusters had multiple distinct summits. These clusters were divided into distinct peaks, with one for each distinct summit, and the boundaries defined by the most proximal downstream and upstream start/end positions within the cluster. Peaks assigned to regions of the assembly not corresponding to any chromosome, and peaks with a width > 3,500 bases were eliminated. The *getfasta* function of bedtools and a custom R script were used to identify and eliminate peaks with > 70% repetitive elements. The final combined atlas contained 70,323 peaks. The R packages ‘GenomicRanges’ and ‘ChIPpeakAnno’ were used to assign peaks to the closest gene according to the following principles ^88,102,103^. Peaks 2,000 bases upstream or 500 bases downstream of a transcription start site were considered ‘promoter’ peaks. Non-promoter peaks within the body of a gene were considered ‘intragenic’ peaks. All other peaks 100,000 bases up or downstream of a gene body were considered ‘intergenic’. All other peaks were not assigned to any specific gene. Differential accessibility analysis was carried out using the ‘DESeq2’ package, with the formula ‘∼ Celltype + Replicate’ where Celltype is either *Il10^neg^*, *Il10^recent^*, or *Il10^stable^* and replicates are the separate samples from which each of the three populations were sorted ^93^. Differential accessibility analysis and statistical testing was performed for all pairwise comparisons of ‘Celltype’: *Il10^neg^*, *Il10^recent^*, *_Il10_stable*.

### Motif identification and model generation

Motif discovery in peaks and model creation were carried out as previously described with some modifications ^44^. Motifs for all mouse transcription factors (TFs) were downloaded from CisBP v2.00 ^104,105^. TFs with mean FPKM > 1 in any cell population were used for further analysis. This resulted in 319 motifs for 188 TF encoding genes. The ‘AME’ software from the MEME suite (version 5.3.0) was used to identify which of these motifs were enriched in the sequences corresponding to the combined peak atlas ^106,107^. At this stage the best motif for each TF encoding gene, defined as being detected in the highest fraction of peaks, was chosen for further analysis. Then, the ‘Tomtom’ software from the MEME suite was used to determine closely related motifs ^108^. A custom R script was used to group motifs according to the follow principles. Motifs were grouped if their ‘Tomtom’ E-value was < 0.00001 and if they were in the same protein family (according to CisBP). Then, groups were merged on the basis of overlapping members until each gene belonged to at most one group. This resulted in 142 motif groups. Finally, motifs not significantly enriched in the peak atlas (‘AME’ adjusted p-vale < 0.01) were excluded. This resulted in 76 TF encoding genes organized into 58 groups. Then, the ‘FIMO’ software from the MEME suite was used to identify individual instances (*p*-value < 0.0001) of each of the 76 motifs across the entire peak atlas ^109^. Then, motif families occurring in fewer than 2% of peaks were eliminated. This resulted in a final set of 57 motifs within 40 groups. Finally, a peak-by-motif matrix was generated wherein 1 indicated at least one instance of a motif belong to that family and 0 indicated no motif.

The ‘ridge’ package in R was used to fit a linear ridge regression for the log_2_ transformed fold change (log_2_ FC) in accessibility at each peak as a function of the peak-by-motif matrix ^45,46^. This package applies an algorithm for semi-automatically determining the optimal ridge parameter(s) to use in order to maximize model performance. This was done separately for the *Il10^stable^* vs *Il10^recent^* log_2_ FC (svr model) and *Il10^stable^* vs *Il10^neg^* log_2_ FC (svn model). Motif families with significant coefficients in the models (*p*-value < 0.001) were used in subsequent analyses. At the same time, linear ridge regressions were fit as above, except with each motif individually removed from the matrix, generating a series of ‘zeroed-out’ models. Then, for sets of peaks of interest (e.g. those associated with a specific cluster of genes), the correlations between the actual log_2_ FC and the log_2_ FC predicted by the svr or svn model as well as the correlations between the actual log_2_ FC and the log_2_ FC predicted by the ‘zeroed-out’ svr or svn models were determined. Decreased correlation for the ‘zeroed-out’ model was assumed to be indicative of the ‘zeroed-out’ motif disproportionately contributing to the model’s predictiveness at those specific peaks, and therefore potentially regulating accessibility.

### Plotting RNA-seq and ATAC-seq tracks

The UCSC utility faCount was used to determine the effective genome size of the ‘reformed mm39’ genome ^110^. The bamCoverage function of deeptools (version 3.5.1) was used to generate ‘bigwig’ files for ATAC- and RNA-seq tracks, with the following command ^111^. bamCoverage -b *Sample*.duprm.bam --effectiveGenomeSize *x* -bs 1 --maxFragmentLength *y* -- scaleFactor *z* -o *Sample*.normdt.bw

Above, *Sample* stands in for each *Il10^neg^*, *Il10^recent^*, or *Il10^stable^* replicate; *x* was the effective genome size as defined above; *y* was the equivalent value determined by STAR during alignment (*(2^winBinNbits)*winAnchorDistNbins*) for ATAC-seq or the default for RNA-seq; and *z* was the inverse of the size factors determined by DESeq2.

### TCR deletion RNA-seq

5,000 cells were sorted per population per replicate for bulk RNA-seq. RNA was extracted and libraries prepared using SMARTer Stranded RNA-Seq Kits according the manufacturer’s protocols (Takara) by the Integrated Genomics Operation (IGO) Core at MSKCC. Paired-end 50bp reads (20 to 30 million per sample) were sequenced on an Illumina HiSeq 3000 by IGO.

### TCR deletion RNA-seq data processing

Samples were processed and aligned using Trimmomatic (version 0.39), STAR (version 2.7.3a), and samtools (version 1.12), with the following steps, where *Sample* stands in for each d10 or d21 TCR^+^ or TCR^−^ replicate ^90–92^.

TrimmomaticPE *Sample*_R1.fastq.gz *Sample*_R2.fastq.gz -baseout *Sample*.fastq.gz

ILLUMINACLIP:TruSeq3-PE.fa:2:30:10 LEADING:3 TRAILING:3 SLIDINGWINDOW:4:15 MINLEN:36

STAR --runThreadN 6 --runMode alignReads --genomeLoad NoSharedMemory --readFilesCommand zcat --genomeDir mm39_100_RNA --readFilesIn *Sample*_1P.fastq.gz *Sample*_2P.fastq.gz -- outFileNamePrefix *Sample* --outSAMtype BAM Unsorted --outBAMcompression 6 -- outFilterMultimapNmax 1 --outFilterMismatchNoverLmax 0.06 --outFilterMatchNminOverLread 0.35 -- outFilterMatchNmin 60 --alignEndsType EndToEnd

samtools sort -@ 4 -n -o *Sample*.bam *Sample*Aligned.out.bam

samtools fixmate -@ 4 -rm *Sample*.bam *Sample*.fixmate.bam

samtools sort -@ 4 -o *Sample*.resort.bam *Sample*.fixmate.bam

samtools markdup -@ 4 -l 1500 -r -d 100 -s *Sample*.resort.bam *Sample*.duprm.bam

samtools index -@ 4 -b *Sample*.duprm.bam

This procedure resulted in retention of all uniquely aligning reads, with PCR and optical duplicates removed, to be used for downstream analysis. Reads aligning to genes derived from the reformed GTF were then counted using a custom R script relying on the ‘GenomicAlignments’, ‘GenomicRanges’, and ‘GenomicFeatures’ packages with default counting parameters ^88^. Differential expression analysis was carried out using the ‘DESeq2’ package, with the formula ‘∼ Conditon’ where Condition is each unique combination of timepoint and whether cells lost or retained cell surface TCR ^93^. Fragments per kilobase mapped (FPKM) normalized counts were extracted using the *fpkm* function of DESeq2. Differential expression analysis and statistical testing was performed for TCR^+^ versus TCR– replicates at each timepoint. Differential expression analysis was performed on all genes, but genes with FPKM counts below the mean FPKM count of *Cd8a* (a gene functionally not expressed in Treg cells), genes with 0 counts in the majority of samples, or genes corresponding to immunoglobulin or TCR variable, diversity, or junction segments were eliminated for subsequent analyses. This process did not remove any significantly differentially expressed genes, except immunoglobulin or TCR variable, diversity, or junction segments, whose differential expression was not interpretable. K-means clustering was performed with R using per-gene Z-score normalized counts of genes differentially expressed (adjusted *p*-value < 0.05) in any pairwise comparison between the three cell populations, with 7 clusters chosen on the basis of preliminary hierarchical clustering. TCR activated and repressed genes were defined as genes which lost and gained expression in Treg cells ablated of the *Tcra* gene ^52^.

### *In vitro* transduction

To produce virus, HEK293T cells grown to 70% confluency in complete DMEM (10% FBS) were transfected with 25ug modified MIGR1 vector, 20ug pCL-Eco helper plasmid, and 135uL Fugene transfection reagent (Promega). The vector structure was as follows: 5’ LTR – MESV psi – gag – lox71 (sense) – IRES-mAmetrine (antisense) – Maf/Rorc/empty (antisense) – lox66 – 3’ LTR. This enabled Cre mediated swapping of the Maf/Rorc/empty – IRES – mAmetrine reading frame into the translated, sense orientation. 48 hours after transfection, the virus was concentrated using RetroX concentrator reagent (Takara). Meanwhile, Treg cells were sorted as CD4+TCRβ+Thy1.1+tdTomato– and cultured in complete RPMI (10% FBS) containing 1000U/mL IL-2, on anti-CD3/CD28 (2ug/mL) pre-coated plates. 24 hours post-culture, cells were ‘spin-fected’ at 1250xg for 90 minutes at 32°C in complete RPMI containing concentrated virus, 8ug/mL polybrene and 1000U/mL IL-2. Three days later, cells were isolated and analyzed by flow cytometry.

### Statistical analysis and data plotting

All statistical tests were carried out in R using base R or the indicated packages, or by the indicate software or python algorithms. Plots were generated in R, using the ggplot2, patchwork, RColorBrewer, ggrepel, and Seurat packages ^94,112–115^. Inkscape (version 1.0.2) was used to modify and assemble plots for presentation ^116^.

## Notes

### Summary of Updates

New/Revised Figures: Extended Data Figures 3-10

